# What Could Go Wrong? Promoting Success by Planning for Failure in Label-Free Biosensor Assay Development

**DOI:** 10.1101/2025.09.16.676687

**Authors:** Sajida Chowdhury, Samantha M. Grist, Avineet Randhawa, Maggie Wang, Karyn Newton, Lauren S. Puumala, Yuting Hou, Lukas Chrostowski, Sudip Shekhar, Karen C. Cheung

**Affiliations:** Department of Electrical and Computer Engineering, The University of British Columbia; Centre for Blood Research, The University of British Columbia; School of Biomedical Engineering, The University of British Columbia; Stewart Blusson Quantum Matter Institute, The University of British Columbia

**Keywords:** Label-free biosensors, root cause analysis, assay optimization, silicon photonic biosensors, lightweight experimental workflow, continuous improvement

## Abstract

Label-free biosensors offer powerful platforms for detecting molecular interactions, but developing robust assays on these systems presents several challenges due to the complexity of the testing systems and intersection of disciplines. In this work, we describe a lightweight, 3-step experimental workflow supporting troubleshooting and root cause analysis that we developed during the design and optimization of silicon photonic biosensor assays in an academic research setting. Because such environments often lack the resources and formal quality-management infrastructure available in industry, our approach emphasizes practicality and ease of adoption. Drawing from our own assay failures and successes, we identify common failure modes, propose structured troubleshooting workflows, and provide case studies illustrating the application of established frameworks, including the Five Whys, the Plan-Do-Check-Act cycle, Open-Narrow-Close and the Ishikawa (fishbone) diagram. By applying this framework to our research, we increased our assay yield by a factor of 2, from ~45% to ~90%. This paper aims to support other teams engaged in label-free assay development by equipping them with practical tools and experimental best practices.

**Highlights:**

- We doubled assay yield from ~45% to ~90% by planning for failure, not just success.
- Tailored troubleshooting frameworks help pinpoint and solve various assay failures.
- Applying innovative problem-solving to workflows accelerates biosensor development.
- Systematic troubleshooting improves assay development across a range of lab settings.
- A 3-step workflow highlighting what could go wrong supports continuous improvement.

## 1. Introduction

Label-free biosensors provide real-time, high-sensitivity detection of molecular interactions and have been applied in diverse domains, from clinical diagnostics to environmental monitoring [1]–[6]. These platforms are compatible with a variety of assay formats and receptor types, enabling flexible experimental design, and they can allow analysis of binding kinetics, giving insight into the dynamics and affinities of molecular interactions [7]. Additionally, some platforms support multiplexing, high-throughput screening, or integration with automated pipelines [8]. In certain cases, sensors can also be regenerated and reused for multiple experiments [9], increasing efficiency and cost-effectiveness.

Despite their advantages, developing robust assays on label-free platforms is challenging due to the sensitivity of the systems to a wide array of experimental variables [10]. Small inconsistencies in fluidic delivery, sensor surface functionalization, or reagent preparation can lead to ambiguous or non-reproducible results. Instrumentation issues, such as bubble formation and fluidic blockages or microfluidic channel misalignment can further challenge measurement reliability. Additionally, human factors including protocol adjustments across trials, mislabeling or improper storage of reagents, inconsistent or sparse documentation, and variable experience levels among team members can exacerbate these issues. Given the cost of reagents and the complexity of instrumentation, failed assays carry a high burden in terms of time and resources. Root cause analysis (RCA) frameworks [11] therefore have particular value in the biosensor field, where they can reduce inefficiencies and improve reproducibility.

RCA is a structured problem-solving approach that seeks to identify the underlying causes of a failure in order to implement targeted, preventive solutions. Rather than only addressing symptoms, it focuses on eliminating the fundamental drivers of the problem [11]. For example, re-running experiments to correct inconsistent binding signals addresses only the symptom, whereas identifying unstable fluid delivery, reagent degradation or poor surface functionalization (and the gaps in protocols and training that permitted these issues to occur) targets the root cause. In industrial and clinical settings, RCA is often formalized into systematic processes that involve defining the problem, collecting evidence, analyzing contributing factors, and developing targeted interventions. This approach has been investigated across a range of areas, from diagnosing bearing failures in motors [12], to preventing medical errors in health care [13], identifying missed diagnostic opportunities in post-endoscopy cancer [14], and improving reliability in CNC plasma cutting systems [15]. Despite this versatility, structured strategies for troubleshooting and RCA in academic research laboratories remain underreported [16]. Several factors may contribute to this gap: academic projects typically prioritize novelty and rapid publication over process optimization; formal quality-management systems are rarely resourced or incentivized; and high personnel turnover can disrupt the continuity required for systematic analyses. As a result, many research groups rely on ad hoc trial-and-error troubleshooting, which may resolve immediate symptoms but leave the true root cause unaddressed, ultimately impeding reproducibility and knowledge transfer.

To provide structured alternatives supporting RCA, several frameworks can be applied and adapted to lightweight formats in academic laboratories and other environments (e.g., startup companies) that lack dedicated resources for RCA. These frameworks provide organized approaches to identify underlying issues, prioritize interventions, and implement changes that prevent recurrence:

- **Five Whys [17]**: Repeatedly asking “why?” to trace a problem to its root cause. This simple but effective method can facilitate rapid diagnosis across a wide array of assay challenges from inconsistent binding signals to unexpected baseline drift, making it a valuable first-line tool for RCA in label-free biosensor development.
- **Plan-Do-Check-Act (PDCA) cycle [18], [19]**: An iterative process for continuous improvement that emphasizes planning interventions, testing them, evaluating outcomes, and refining protocols. PDCA is well suited to complex, multi-step assay workflows where incremental adjustments can be helpful.
- **Open-Narrow-Close (ONC) [20]**: A problem-solving approach that encourages broad exploration of possible causes (Open), methodical filtering to the most plausible hypotheses (Narrow), and targeted testing to confirm root cause (Close). ONC is particularly suited to situations with multiple interacting failure sources.
- **Ishikawa (fishbone) diagram [21], [22]**: A visual framework that maps potential contributing factors across different categories. This method provides a holistic view of assay performance issues and facilitates team-based discussions around troubleshooting.

In this manuscript, we present a lightweight, data-driven workflow for label-free assay development that integrates these established frameworks. The workflow is designed to support interdisciplinary teams with varying levels of experience in wet-lab experimentation, data analysis, and biosensor engineering. Drawing on our own successes and failures in developing silicon photonic (SiP) biosensor assays, we demonstrate how systematic troubleshooting can improve assay reproducibility and yield from ~45% to ~90%. To contextualize our best practices, we report several examples of failures and challenges that led to the development and continuous improvement of this workflow; in doing so, we aim to describe some of the many aspects of biosensor development that can go wrong, and also provide specific examples of how our workflows can help turn failures into valuable learning experiences. Because tools cannot provide benefit if the overhead is too high for researchers to use them, we describe an approachable workflow that balances proven best-practice frameworks with a lightweight format. By reflecting on the process of resolving complex assay issues and sharing the challenges, pitfalls and solutions we encountered, this work aims to provide practical guidance that can streamline the development of label-free biosensor assays while supporting the training of highly qualified personnel and translation of research results.

## 2. Methods

### 2.1 Experimental workflow for continuous improvement in assay development and troubleshooting

Figure 1 describes the 3-step data-driven experimental workflow that our team has developed for label-free assay development supporting effective troubleshooting and RCA. In developing this workflow, we aimed to improve assay yields while maintaining a lightweight approach. Development and characterization of label-free biosensors is challenging because of the complexity of the systems required for their development. For example, for SiP biosensors (Figure 1(a)), the system comprises: (i) the SiP chip itself; (ii) the photonic readout system used to acquire data from the sensors (typically including a swept-tunable laser and optical detectors as well as readout electronics); (iii) the integrated microfluidics to deliver reagents and samples to the sensor surface; (iv) the functionalization chemistry that confers specific detection; (v) the thermal control and/or reference sensors typically used to improve system stability; (vi) data acquisition and analysis software; (vii) the fluidic control system; and (viii) the reagents used for sensor functionalization and detection. In addition to the complexity of the physical system, the interdisciplinary team required for this type of development may confer additional complexity: some team members may have never worked in a wet-lab environment before, while others may not have programming or photonics training. In this kind of environment, effective documentation and communication between team members are critical to support collaboration. Our workflow includes strategies for documentation, logging, and data analysis that we have found to be particularly important for these types of environments (Figure 1(a-b)).

**Figure 1.**
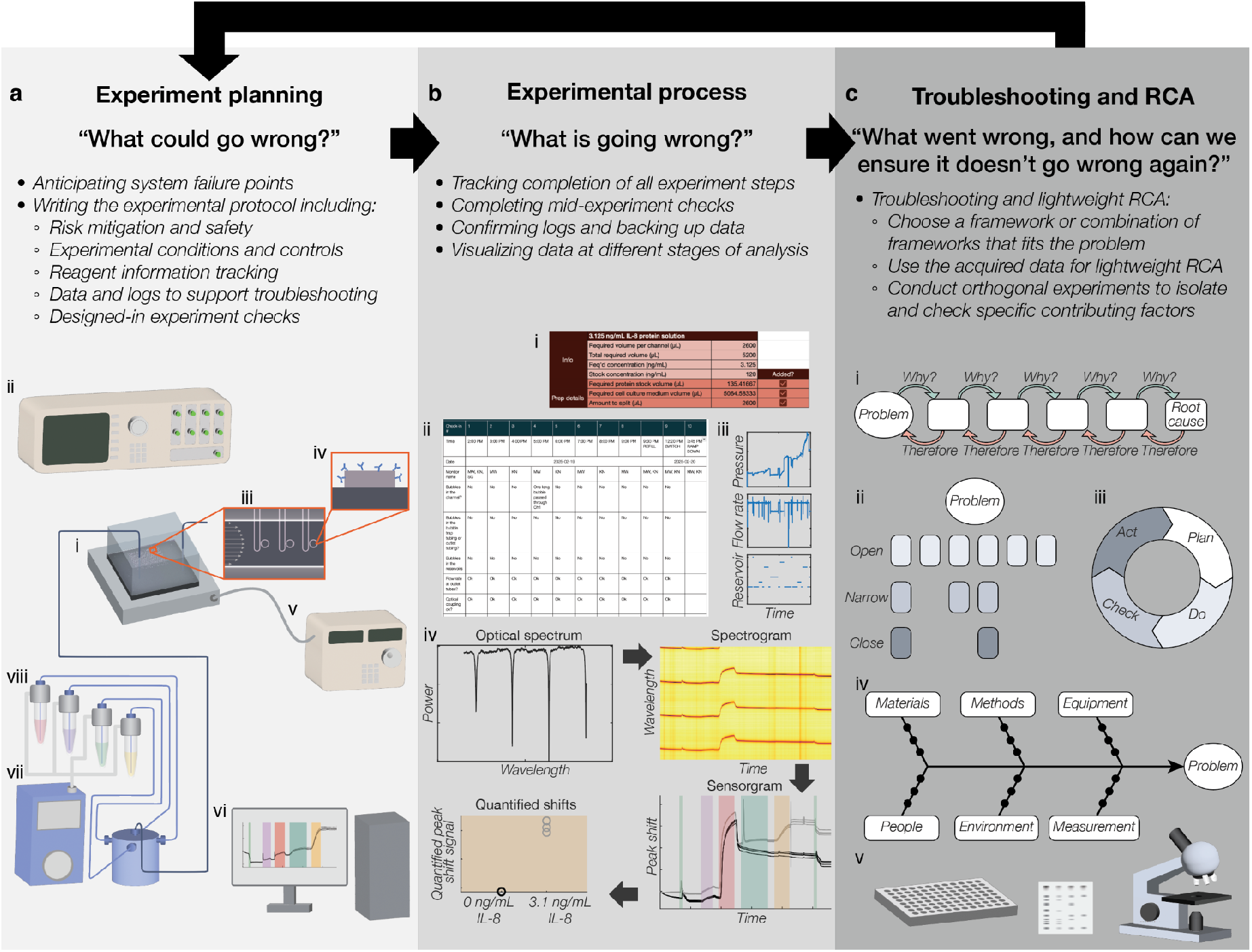
Data-driven 3-step experimental workflow for iterative label-free assay development supporting effective troubleshooting and root cause analysis (RCA). The workflow is composed of three steps **(a-c)**: experiment planning, experimental process, and analysis, troubleshooting, and RCA. **(a)** In the experimental planning phase, we ask “what could go wrong” with all components of the setup and system, and write the experimental protocol with all potential failure points in mind. For SiP biosensors, the system is composed of **(i)** the SiP chip, **(ii)** photonic readout system, **(iii)** integrated fluidics, **(iv)** functionalization chemistry, **(v)** thermal control and/or referencing, **(vi)** data acquisition and analysis software, **(vii)** fluidic control system, and **(viii)** assay and reagents. Each of these components has the potential to fail in several ways, which are considered during experiment planning. **(b)** In the experimental process phase, we ask “what is going wrong” and log and track as many factors as possible that could confound our downstream analysis. This tracking includes **(i)** checklists for reagent preparation and experiment setup, **(ii)** frequent system checks of potential failure points during automated experiments, **(iii)** data logs for parameters like temperature, pressure, flow rate, and reagent/reservoir being delivered, **(iv)** visualization of the data at various stages of analysis (for SiP sensors, this includes the individual raw spectrum at each time point, the raw data spectrogram (with power visualized on a colormap scale), the sensorgram depicting resonance peak shift signal vs. time, and the quantified signal). **(c)** In the analysis, troubleshooting, and RCA phase, we ask “what went wrong, and how can we ensure it doesn’t go wrong again”, and employ tools like **(i)** the Five Whys RCA framework, **(ii)** the Open-Narrow-Close problem-solving strategy, **(iii)** the Plan-Do-Check-Act (PDCA) framework for continuous improvement, **(iv)** Ishikawa (fishbone) diagrams to support RCA, **(v)** orthogonal measurements like ELISA, dot and Western blots, and microscopy support (i-iv).

Academic research laboratories have unique challenges including high researcher turnover (e.g., as students graduate and finish summer research terms), needs for training new team members, and limited administrative resources. Because of these constraints, strategies for troubleshooting, RCA, and project management that are best practices in industry and government often do not translate well to academic research labs. Similar to how the Scrum project management framework has been adapted to the LabScrum framework [23], we sought lightweight tools for troubleshooting and RCA. In developing this workflow, we have implemented multiple frameworks (Figure 1(c)) and describe their application to specific problems that we encountered in Section 3.

Our iterative workflow builds upon classical concepts of iterative experimental design using an experimental cycle [24] and Kolb’s experiential learning cycle [25], [26] and aims to leverage these potential issues into learning experiences for the team. “Things going wrong” are inevitable, and by designing a workflow around this fact, we aim to make the most of every issue that arises: each “thing that goes wrong” is not a failure but a valuable piece of data from which to learn and improve our experiments, processes, and training. Our workflow (Figure 1) is thus structured around three questions: (1) in the experiment planning phase, we ask “what could go wrong?” and design detailed experimental protocols with this in mind; (2) in the experimental process phase, we constantly ask “what is going wrong?” by tracking, logging, and conducting experimental checks; (3) in the data analysis, retrospective, and troubleshooting phase, we ask “what went wrong, and how can we ensure it doesn’t go wrong again?” and use lightweight frameworks to investigate. These learnings are then factored into the planning stage of the next experiment. By approaching our experiments in this way, we seek to learn from both best practices and our team’s experience in a cycle of continuous improvement.

### 2.2 Supporting best practices for troubleshooting

During the experiment-planning phase (Figure 1(a)), our efforts are organized into a comprehensive experimental protocol document. In addition to implementing the first half of the experimental cycle (formulating the research question and hypothesis, outlining the experimental groups and controls, and designing the methods), this document guides the researcher through the process of brainstorming everything that could go wrong with the experiment and how to mitigate (a) the chance of this specific failure point going wrong in the first place and (b) the chance of this failure point impacting team safety, equipment functionality, and experimental results. Thus, our protocols have sections including the goals, summary of previous results, hypotheses and expectations, controls and experimental groups, hypothesis-testing strategy, required reagents and equipment, risk mitigation, and detailed protocol. Within the detailed protocol we include spots for information to be filled in during the experiment, including specific photonic chips and microfluidic gaskets and moulds used for the assay, timing of various steps, ambient environmental conditions, fluid volumes (e.g., for reagent spotting), and microscope images of the SiP chip at various stages of the experiment. This information supports downstream troubleshooting, promoting as much consistency as possible between experiments and allowing us to retroactively look for relationships between variable factors in the experiment and results. To better illustrate our protocols, we have included an adapted diagram depicting the experimental cycle and how we use documentation for its implementation (Figure S1) as well as an example protocol document and spreadsheet as a part of the Supplementary Information (SI).

To further support this kind of troubleshooting and mitigate reagent challenges, reagents are also tracked. In addition to recording the vendor, product, and lot numbers, we record the receipt and opening dates of each reagent, how it has been stored and aliquoted, and when each aliquot was opened and thawed. We also use planning spreadsheets with checkboxes to be selected as each reagent is added and each solution is mixed during solution prep. This kind of logging not only reduces the chance of human error during solution preparation, but also supports troubleshooting by clearly documenting exactly how each solution was prepared in each trial. By planning all of this information ahead of time and clearly indicating the factors that need to be checked and documented during the experiment itself, we seek to promote experimental workflows that are organized and less prone to error, reducing the load on the researcher during the experiment so they can focus on carrying out the steps (like following a recipe) rather than the parallel tasks of planning, troubleshooting, and experimentation.

During the experimental process (Figure 1(b)), we continually check in on the factors we identified by asking “what is going wrong?”. In doing so, we follow the experimental protocol exactly (continually checking our work) and fill in all required fields of the protocol and reagent preparation sheets (Figure 1(bi)). We log important information relevant to troubleshooting (Figure 1(biii)), such as the measured photonic chip temperature, the pressures, flow rates, and reagent reservoirs being delivered during the assay, time-lapse images of the microfluidic channels to monitor for bubbles, and logs of when automated alignment of the optical I/O is performed during long assays. We also define experiment check-in points (Figure 1(bii)), where researchers log all checks in a table in order to monitor the experiment for factors that can go awry. For example, for our SiP biosensor assays we typically perform hourly checks during the critical first ~6-12h of the assay and check for key aspects of the microfluidic and optical I/O systems as well as check on the raw data and stability. Since bubbles have a high probability of disrupting assay results, we check for bubbles in the channel as well as input and output tubing. We check the pressures and flow rates being delivered by our microfluidic flow-control system to monitor for potential blockages or flow sensor issues. We also perform a qualitative check of the effluent flow rates at the system outputs to make sure that they match the logged flow rate data and check for leakage along the fluidic path of the system. We check that data acquisition software is still running as expected, optical I/O is working seamlessly, and that the data and time-lapse imaging files are being generated. Finally, we perform quick checks that the data are as expected by viewing the raw optical spectra and spectrogram to look for unexpected instability or coupling (I/O) issues. In our analysis frameworks, we build in the ability to view the data at various stages of analysis (Figure 1(biv)). Although the quantified binding shift signal may be the final metric of interest, the ability to view intermediate stages of analysis is critical for troubleshooting and understanding system issues. These include the sensorgram (tracked resonance peak position vs. time for each sensor after curve-fitting and peak-tracking the individual resonance peaks), spectrogram (a 2D image format of the raw data visualizing wavelength vs. time, with optical power represented on a colourmap scale and each resonance peak visible as a dark lines in the spectrogram), and raw optical spectrum (power vs. wavelength for each timepoint and sensor) formats.

During the final phase of analysis, troubleshooting, and RCA (Figure 1(c)), we analyze the data and prepare an experimental summary document. In preparing this document, we ask “what went wrong, and what can we do to make sure it doesn’t go wrong again?”. To help us answer these questions, we use tools like the Five Whys RCA framework (Figure 1(ci)), the ONC problem-solving strategy (Figure 1(cii)), the PDCA cycle framework for continuous improvement (Figure 1(ciii)), and fishbone diagrams to support RCA (Figure 1(civ)). We apply the Five Whys to help dig deeper to find underlying root causes [27]. We use the ONC strategy to consider a range of potential contributing factors to an issue and identify the highest-likelihood factors to analyze with RCA [20]. We use the cyclic PDCA framework to promote iterative continuous improvement and test out potential solutions [19], and use fishbone diagrams to support RCA to map potential contributing factors and encourage the researcher to generate hypotheses for improvement [21]. Each of these frameworks provide a structured methodology, inviting the researcher to dig into the experimental data and additional supporting logs to plan follow-up tests in order to determine what went wrong. As a part of these frameworks, the researcher often circles back to the experiment planning phase to plan the next experiment to test specific hypotheses about the issue’s root cause.

A key part of troubleshooting and RCA is to isolate contributing factors to narrow down to the root cause. Orthogonal assays are an important means to do this, allowing us to isolate key potential failure points and test contributing factors in a higher-throughput manner than that permitted with our SiP biosensor assays. Our team employs several types of orthogonal measurements in our SiP biosensor assay development (Figure 1(cv)). We use the enzyme-linked immunosorbent assay (ELISA) and other immunoassays like dot blots and Western blots to test and validate reagents (e.g., antibodies, antigens, and amplification reagents like streptavidin-HRP). We use various forms of microscopy to assess sensor functionalization and microfabrication, and ellipsometry to assess and validate sensor functionalization. We also design and use other simple tests to check and validate the performance of individual pieces of equipment (e.g., flowing dye solutions using our automated flow control system). Together, these types of orthogonal measurements allow us to efficiently execute the troubleshooting and RCA frameworks to support continuous improvement. Each cycle of learning is incorporated into the answers to “what could go wrong” and their mitigating factors for the next experiment to iteratively improve our assay yield.

### 2.3 Technical methods

In the examples of implementing our workflow covered in our Results and Discussion section, we focus on the development of a sandwich assay to detect interleukin-8 (IL-8), an 8-9 kDa cytokine. In developing this assay, we encountered several challenges that we used as learning experiences to apply and improve our workflow. SI Section 2 briefly describes the technical methods used for this assay and the associated orthogonal experiments that we present.

## 3. Results and discussion: examples of troubleshooting and RCA

### 3.1 The Five Whys RCA framework permits identification and correction of fluidic delivery system issues

In the first example of applying our workflow, the Five Whys framework enabled us to address the root cause of an issue with low binding signals (Figure 2(a)) that we encountered during a series of IL-8 detection assays using covalent, polydopamine (PDA)-mediated functionalization with multiple rounds of detection (Figure 2(b), Figure S2). Between each round of detection in these IL-8 assays, the sensor was regenerated using a pH 2.2 glycine-HCl solution that disrupts the binding of the IL-8 with the covalently attached capture antibody on the sensor surface, permitting another round of binding. This assay did not use enzymatic amplification, since sufficient signal was observed from the SA-HRP binding stage after assay optimization (further discussion in Section 3.4). After analyzing data from a series of assays, we noticed that we were seeing unexpectedly low or negative signal during the SA-HRP assay stage (Figure 2(a)). Using retrospective analysis of our data and logs and targeted troubleshooting investigation, we followed the Five Whys framework to find the root cause of the issue (Figure 2(c)).

**Figure 2.**
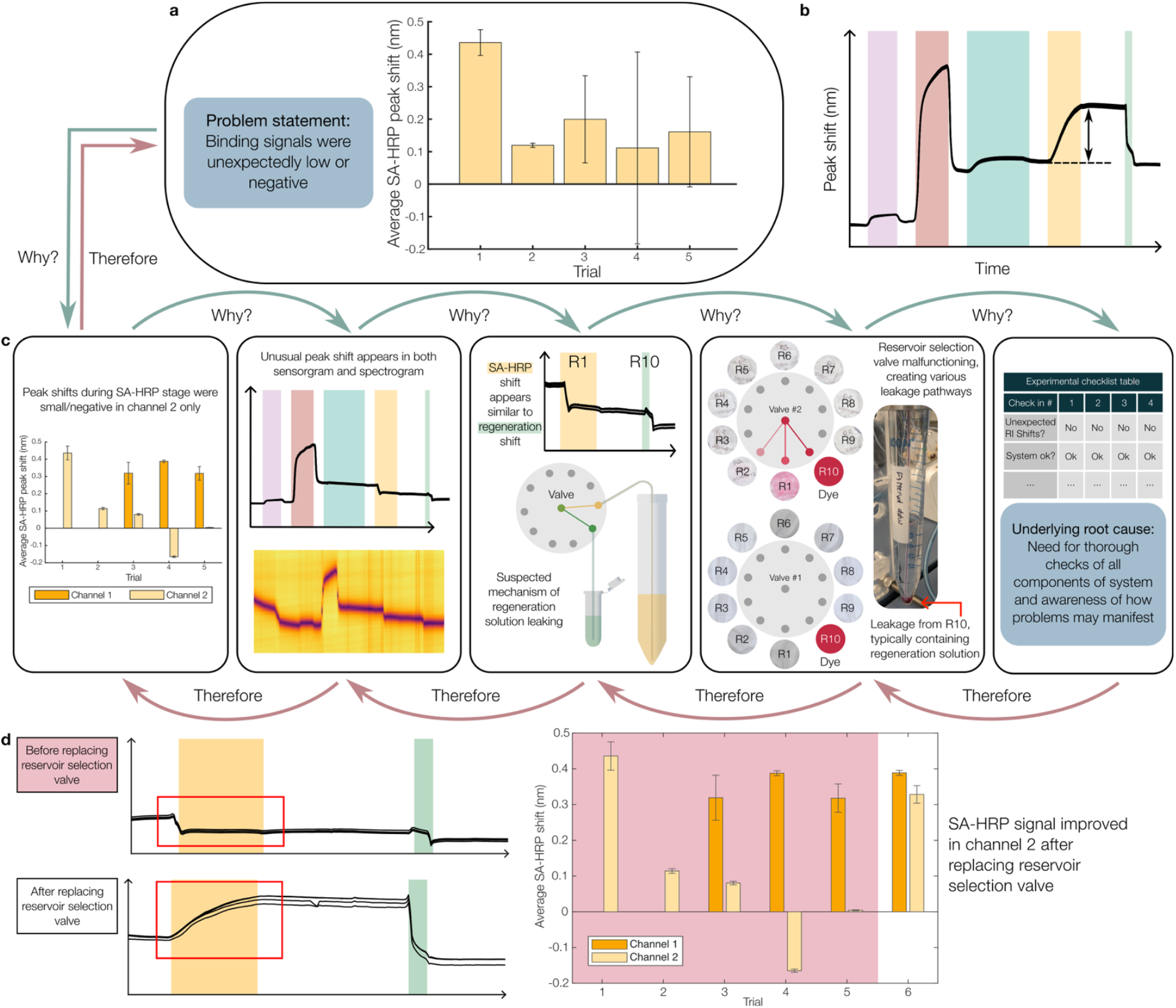
An example of applying the Five Whys framework to determine the root cause of unexpected, microfluidic channel-specific low signal. **(a)** First evidence of the problem: the average signal from the SA-HRP amplification assay stage decreased after trial 1, and variability in signal increased. Bar plot shows SA-HRP binding signals in the first binding cycle with an IL-8 concentration of 3.125 ng/mL. The bars depict averaged signal across two microfluidic channels; however, no data were available for channel 1 in trials 1 and 2 as it was used as a negative control. **(b)** Expected sensorgram for one binding cycle, highlighting the SA-HRP signal that was quantified in the bar plot. The assay stages are overlaid with different colours with purple indicating the BSA challenge stage, red indicating the IL-8 sample stage, teal indicating the IL-8 detection antibody stage, yellow indicating the SA-HRP stage, and green indicating the regeneration stage as described with schematics of the assay in Supplementary Figure S2 and SI section 2.1. Assays were typically run with 3-6 binding cycles. **(c)** Illustration of the process of applying the Five Whys to this issue and determining its root cause: leakage within a rotary reservoir-selection valve within our commercial fluidic control system that led to our regeneration solution contaminating the fluid delivered from the SA-HRP reservoir. **(d)** Zoomed-in view of sensorgrams during the SA-HRP and regeneration stages before and after troubleshooting the issue. Difference in SA-HRP peak shift is highlighted by the red box. The bar plot shows quantified peak shifts from both channels, peak shifts from the five trials before troubleshooting highlighted in red. The mean (bar heights) and standard deviation (error bars) of the signals from 4 sensors in each microfluidic channel and trial are depicted.

Our first “why” prompted us to look for trends in this signal, and we were able to isolate it to one of our microfluidic channels. To determine why the signals were low, we looked at both the analyzed and raw data, and identified an unexpected negative shift during the SA-HRP step. This shift appeared quite similar visually to the shifts we see during the regeneration stage of these assays, so we hypothesized that there was a malfunction occurring within our fluidic system that was delivering the regeneration solution instead of SA-HRP, or causing them to mix together. To get to our next “why”, we tested this hypothesis. After examining the logs of the delivered reservoir, flow rate, and pressures, we determined that there was no error in the fluidic protocol or software that would deliver the wrong solution at that stage, so we developed a method to test for leakage within the system using food dye (SI Section 2.2). We were able to identify several leakage pathways within the reservoir selection valve, including one between the regeneration solution and SA-HRP reservoirs. The same testing was conducted on the other channel’s valve and no evidence of leakage was found. We kept careful documentation of all of our findings, and this documentation was extremely helpful to support correspondence with the vendor of the component about the issue. After replacing the component, the symptom of the issue was addressed and the two channels showed similar signal (Figure 2(d)).

Through this process, we identified the root cause of the issue as being not only the failure of the component itself, but also a lack of thorough checks of all the components within the system and intuition on how possible problems may manifest. With so many parts of the system and our experiments that have potential to fail, our team first focused on the possibility of human error and reagent or protocol issues after trials 2-4 and did not consider the possibility of failure of the automated fluid control system as a high-likelihood cause. The RCA conducted after trial 5 allowed us to identify the root cause of the issue and implement several changes to our workflows to help our team catch similar issues earlier. These solutions will help to catch future system issues, even in cases when the mechanism of failure may not yet be known to us. We sought discussion and feedback from the vendor on the completeness of our experimental and maintenance protocols to ensure that any of our practices were likely to cause early failure of the system. The vendor’s generous guidance and feedback was extremely valuable, and having our detailed experimental protocol documents and system maintenance logs already prepared was critical to take advantage of this discussion. Although we did not identify workflows that were likely to increase the chance of component failure, the vendor’s guidance did allow us to improve our maintenance and system cleaning procedures, and we have been vigilant in following the updated protocol with each of our assays. Another change to our experimental workflow was adding additional checks for unexpected refractive index (RI) changes during solution delivery, which will allow us to identify this kind of issue faster in the future. Additionally, the documented protocol that we defined to test for leakage allows us to check the status of the component if we see similar evidence to this issue, as well as during annual system checks.

The takeaways from this process include the importance of rigorous documentation and thorough checks during the experiment. The specific indicators of this problem were irregularities with the spectrogram, which could also be caused by other factors such as issues with the reagents or bubbles within the channels. Having detailed notes and collecting evidence allows an RCA framework to begin to be applied and aids in designing follow-up experiments to be implemented as part of the RCA process, allowing the actual problem to be uncovered quickly. When the problem has been identified, we must first isolate the parts that are within our control and make changes that can help us moving forward. In the context of implementing a solution to this leakage problem, we needed to balance the benefit of routine testing (which could completely control the issue) with the resources required for its implementation. This comes with a tradeoff as the procedure we developed for thorough testing of all leakage pathways takes up a significant amount of time and effort (~1 full day), and the vendor has indicated that the likelihood of failure of this component is very low. The procedure that we implemented (thorough data checks alongside annual component testing, with additional checks implemented as-needed if anomalies are seen) aims to balance these factors. More broadly, this example illustrates the importance of considering all potential failure points in the system (even commercial equipment with low likelihood of failure) and how those failures would manifest (what to look for in the data and how to specifically test for those failures to diagnose the issue quickly). Any component can fail, whether it is a piece of equipment, software, or a certain reagent within its guaranteed period. Our goal as researchers is to be aware of this fact (which we cannot control) and design our protocols to (1) minimize the chance of failure and (2) quickly diagnose issues when they arise (which is the root cause within our control).

### 3.2 Iteratively reducing microfluidic blockage issues with the Plan-Do-Check-Act (PDCA) framework

Blockages represent an important source of variability and failed experiments in microfluidic assays. Complete or partial obstruction of the flow path elevates system pressure, disrupts fluid delivery, and in severe cases, halts reagent transport entirely, leading to unusable data. Pressures exceeding 1000 mbar are not supported by our fluidic control system. Once this threshold is reached, the effective flow rate progressively declines and can reach 0 μL/min. These failures not only compromise assay yield but also waste valuable time and reagents. The troubleshooting framework that we found most applicable to reducing the incidence and impact of microfluidic blockages was the PDCA cycle. In Figure 3 we demonstrate an example of applying the PDCA workflow to iteratively reduce blockage issues in the microfluidic channels interfaced with our SiP biosensors, illustrating three cycles of iterative improvement.

**Figure 3.**
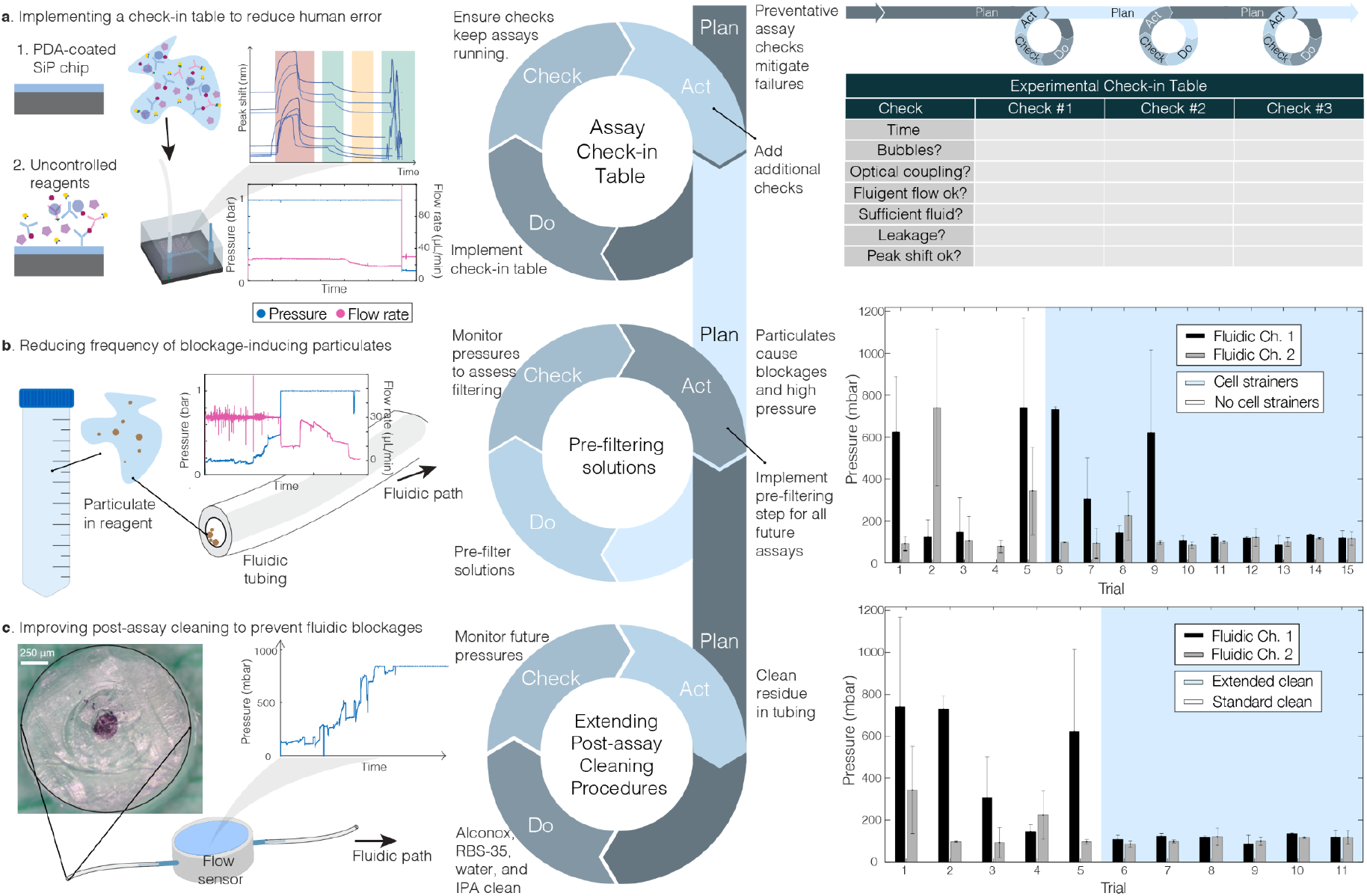
Plan-Do-Check-Act (PDCA) workflow for iterative label-free assay development supporting mid-assay troubleshooting, and three example iterations. The Plan-Do-Check-Act is composed of four phases: firstly, identifying an issue and creating an actionable ‘Plan’. econdly, during the ‘Do’ phase, implement a solution and verify it during the ‘Check’ phase of PDCA. Lastly, during the ‘Act’, implement changes to standardize assay procedures. **(a)** shows a human error example where assay clamps were not unblocked which led to leakage, high pressures, and uncontrolled reagents reaching the photonic ring resonators. An assay check-in table was implemented which also permits PDCA each hour of the assay. Plan: Define what needs to be monitored during the assay and establish check-in intervals. Decide what counts as normal vs abnormal. Do: Perform the assay and complete the check-ins at each scheduled time. Check: Compare logged values and notes against expected outcomes and identify deviations. Act: Fix issues found (e.g., adjust tubing if leaks are present, trim tubing if blockage present, re-make refill samples) and update the protocol and/or schedule if problems repeat. **(b)** Particulates in reagents were identified as a source of flow path blockages and elevated pressures during assays. (Middle) A PDCA cycle was applied: reagents were pre-filtered with cell strainers to remove particulates, and pressures were monitored to evaluate effectiveness. (Right) Bar plots show reduced average supplied pressures when cell strainers were used (333.0 ± 289.5 mbar (n = 5 experiments) prior to cell strainer use and 182.2 ± 177.5 mbar (n = 10 experiments) after). Errors and error bars represent one standard deviation of the average pressures from each replicate experiment. **(c)** (Left) Even after implementing reagent filtering, on-chip immunoassays sometimes resulted in residue buildup within fluidic tubing, contributing to blockages and increased pressures, as visualized by flow sensor and pressure read-outs as well as an image depicting a blockage (dark blue) within clear tubing. (Middle) A PDCA cycle was applied, in which post-assay cleaning steps were extended. (Right) Bar plots show reduced pressures and improved stability following extended cleaning. Error bars once again represent one standard deviation of the average pressures from each replicate experiment.

The PDCA cycle is a spiraling cycle consisting of the ‘Plan’ phase, where problems and potential issues are anticipated and pre-emptive mitigation strategies are planned; the ‘Do’ phase, where mitigation strategies are conducted; the ‘Check’ phase, where mitigation strategies are evaluated; and the ‘Act’ phase, where strategies are modified or edited as required [28]. The PDCA cycle can be applied to label-free biosensor assay development in order to continuously improve assay yield, performance, and replicability by iteratively planning and testing the impact of assay variables and other contributing factors.

Figure 3(a) demonstrates an iteration of the PDCA framework to mitigate human error, which often occurs during long and complex experiments. In one trial utilizing IL-8 detection with enzymatic amplification using 4-chloro-1-naphthol (4CN), an error led to unexpected binding signal shifts. At the beginning of fluidic testing in many of our assays, the effluent tubing is clamped to prevent backflow of reagents across the SiP chip when reagent reservoirs are being switched or installed. In this assay, the effluent tubing was accidentally left clamped from the beginning of the assay, which led to the unexpected assay results. With the fluidic path clamped, the automated fluid control system (which was operating in a constant-flow regime with a feedback flow rate sensor) increased the driving pressure in the assay reservoirs up to the maximum controllable delivery pressure of 1 bar. This resulted in leakage occurring at a few points in the fluidic path where connectors were improperly tightened. This leakage in turn resulted in uncontrolled mixing and delivery of the reagents reaching the SiP sensors located within the microfluidic channels, and unusable assay results.

To mitigate this kind of issue in the future, the team hypothesized that assay checks, implemented using a dedicated assay check-in table in the protocol document, could reduce the likelihood of this issue as well as several other potential failures (including other human errors). In the planning phase, we designed the check-in table contents based on assay variables which were already being measured and observed in a less rigorous manner as part of our process to first consider “what could go wrong”. These checks included: microfluidic flow rate and pressure, presence of bubbles, presence of leakage, observation of optical coupling, observation of peak shift, observation of leakage, and observation of leakage into the vacuum line. In implementing and evaluating this check-in table (the ‘Do’ and ‘Check’ stages) we found that the table allowed us to catch several issues early and allow us to correct them before they impacted the assay results, establishing the success of this intervention. We also sought to continuously improve this check-in table in the ‘Act’ stage, by incorporating additional checks to accommodate other failures as we encountered them. When we encountered issues with flow rate sensors occasionally reporting erroneous values (which can occur with calorimetric flow rate sensors due to blockage or film buildup disrupting the sensor’s mechanism of action), we added in a qualitative check of the flow rate observed at the outlet as well as increased the frequency and rigour of our post-experiment cleaning protocols. In another assay trial involving multiple binding cycles, a reagent was depleted before it was needed again which led to air being introduced to the microfluidic channel. This kind of depletion can occur not only due to errors in initial filling, but also due to inaccuracies in flow rate measurement when a flow rate sensor is used as part of a feedback control system to deliver constant flow to a microfluidic system. Accordingly, in addition to increasing our reagent buffer volumes after this issue, we added a check of the remaining quantity of each reagent to the check-in table. It is interesting to note that the check-in table itself can be seen as an example of a PDCA cycle during each hourly check of the assay: problems can be quickly identified, a solution applied during the check, and evaluated during the next hour if needed. We found overall that the use of this check-in table reduced the frequency of human error during assays and permitted more frequent mid-assay solutions, improving our team’s experimental yield and throughput.

Although the first PDCA cycle reduced incidence of human error that can contribute to microfluidic blockage and a suite of other issues and allowed us to find blockage issues mid-experiment, there are many other factors not related to human error that can contribute to blockages within microfluidic systems. Although finding blockages mid-experiment (before they impact assay results) affords the opportunity to troubleshoot, the troubleshooting process itself has potential to disrupt the running assay. We thus sought to apply additional cycles of the PDCA framework to further mitigate the root causes of blockage formation. For example, Figure 3(b) (Left) demonstrates an assay where we observed a continual pressure climb and subsequently, a complete loss of reagent flow. In this case, delivery of reagent to the chip became halted and uncontrolled, reducing overall assay yield. Pressure increases were attributed to particulates or microfluidic blockages within the fluidic tubing after discussion with the fluidic control system vendor. Thus, we applied the PDCA framework to address the accumulation of residues and trapped particulates inside the tubing and components of the fluidic system, which can lead to increased pressures during experiments. Protein residues and particulates can adhere to the interior of the polymer tubing that connects system components and accumulate at fluidic junctions where the internal diameter narrows (e.g., the entrance of flow sensors, fluidic tubing adaptors where the tubing is compressed, and kinks in the tubing). We hypothesized that particulates and debris within our input reagents may be contributing to the blockages and high pressures that we sometimes observed. We tested pre-filtering all reagents as a solution to reduce the observable (through our logging data) mid-assay pressure rise. This step was standardized across all subsequent experimental protocols. Filtering all solutions using 40 μm cell strainers (MTC Bio SureStrain, Millipore Sigma) as they were pipetted into the final reservoir tubes (immediately prior to connection to the fluidic control system) resulted in a reduction in observed assay pressures (333.0 ± 289.5 mbar (n = 5 experiments) prior to cell strainer use and 182.2 ± 177.5 mbar (n = 10 experiments) after cell strainer use).

In an additional iteration of the PDCA cycle shown in Figure 3(c) we hypothesized that pressure increases could also result from insufficient cleaning of protein residues following assays. In earlier assay trials, we had used a standard procedure of rinsing all reservoir lines with ultrapure water, rinsing one line with 2% RBS-35 detergent ultrapure water (60 minutes at 80 μL/min) to remove residues from the flow sensors and flushing the line with ultrapure water to rinse away residual RBS, rinsing all lines with isopropanol (IPA), and drying all lines with air. After discussion with the fluidic control system vendor, an extended protocol was adopted, particularly for addressing protein residues: this included adding a 2% Alconox Liquinox detergent rinse (3 minutes at 120 μL/min) for all reservoir lines that delivered protein reagents (followed by another thorough water rinse of those lines at 120 μL/min for 45-60 mins), as well as increasing the concentration of the RBS-35 to 10%. The extended cleaning procedure, in combination with pre-filtering solutions, led to decreased average supplied pressure (335 ± 263.5 mbar before extended cleaning and 110.7 ± 15.4 mbar after extended cleaning). Thus, the adoption of the extended cleaning procedure reduced assay variability by standardizing pressures, and was subsequently standardized across all assays showcasing our use of ‘Act’ from the PDCA cycle. We have included a flow chart illustrating our fluidic system cleaning procedures in SI Section 4.

These results illustrate how the PDCA workflow was beneficial in quickly identifying problems and standardizing our assays to improve replicability. Problems can quickly arise during assays (e.g., particulates which quickly form blockages in the fluidic path, inadequate reagent quantity, uncoupling of SiP chip from laser) and monitoring them can allow for in-assay intervention, thus improving our assay yield. Secondly, applying the PDCA workflow to our assays led to progressively greater standardization of conditions in successive assays, lending to more robust detection of IL-8 and a lower incidence of failed assays.

### 3.3 Overcoming challenges with diminished amplification using the Open-Narrow-Close method

Many biomolecules relevant for diagnostic or prognostic purposes exist at picomolar or nanomolar concentrations and can have relatively low molecular weights. For a platform like SiP biosensors that transduces refractive index change (related to the mass of the bound species), this can be a limiting factor for direct detection of analyte binding [29]. One approach that can improve sensitivity involves the use of enzymatic amplification, where additional mass in the form of an insoluble precipitate can be produced continuously and at a rate proportional to the concentration of the analyte of interest [30]. This is typically implemented by immobilizing an enzyme (e.g., HRP) atop an antigen-antibody sandwich and supplying the substrate of that enzyme for a predetermined duration or reaction time [31]. We elected to follow this approach to design an assay permitting quantification of low ng/mL concentrations of IL-8, a sub-10 kDa immune-modulating cytokine with potential utility as a broad inflammatory biomarker. We used a sandwich immunoassay employing passively adsorbed anti-IL-8 capture antibody alongside an HRP-conjugated secondary detection antibody with 4CN as the aqueous-phase enzyme substrate [32]. The assay stages are visualized in Figure 4(a). And further technical details are presented in SI Sections 2.3 and 2.4.

**Figure 4.**
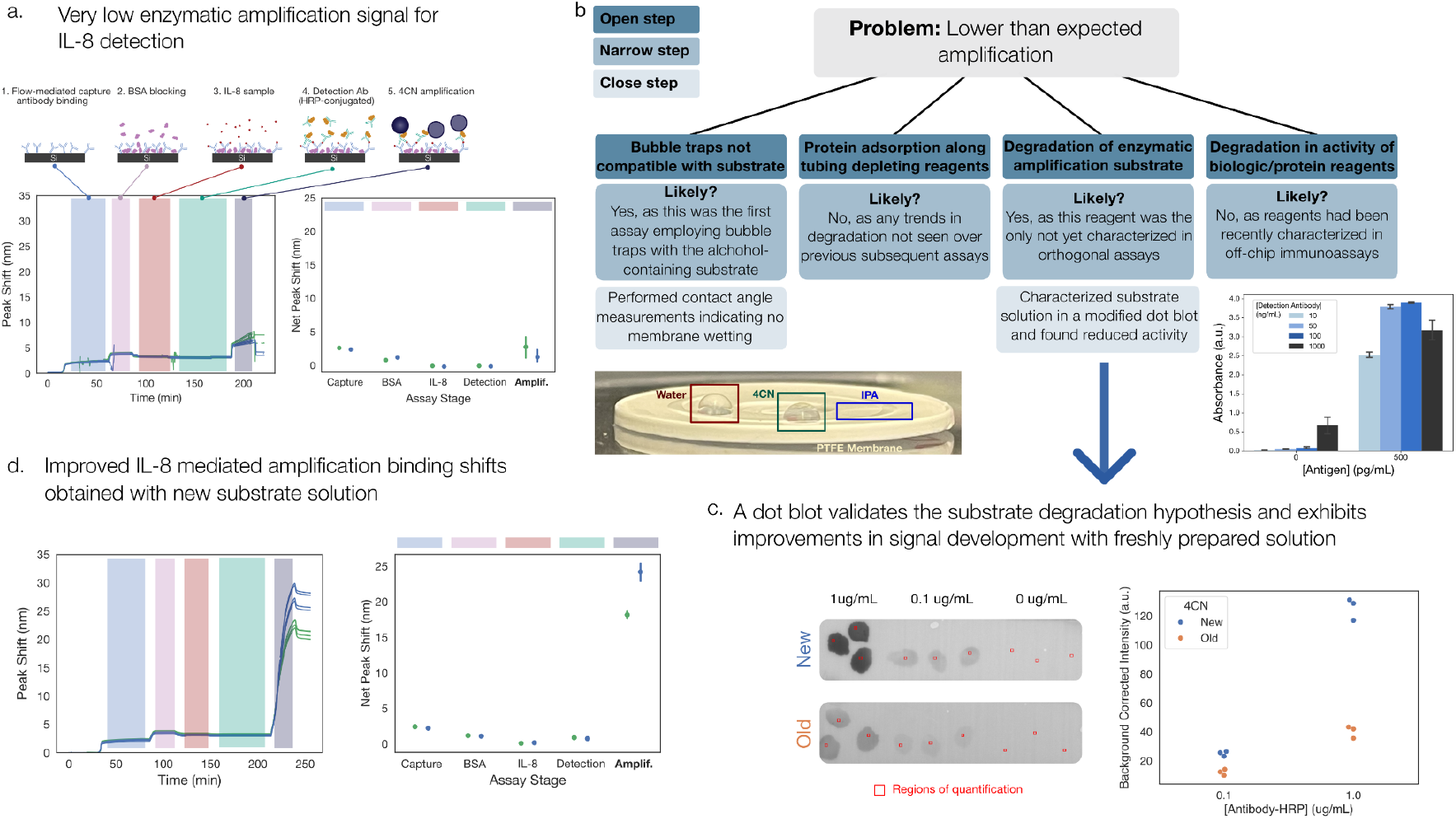
Applying the Open-Narrow-Close framework to troubleshooting low amplification signal. **(a)** A representative silicon photonics assay depicting far lower than expected amplification signal (expected values based on earlier assays employing this sandwich assay were > 20 nm). Dots represent calculated shifts based on pre- and post-stage buffer rinses with error bars depicting standard deviation across four parallel sensor replicates. Individual lines in resonance peak shift vs. time plots depict sensors and colors represent channel replicates. **(b)** The Open-Narrow-Close methodology was applied to generate hypotheses to explain the observed phenomenon, assign strengths of said hypotheses, and prioritize orthogonal testing of those determined to be strong based on available evidence. Below the left hypothesis is a photograph of equal volumes of distilled, deionized water, the 4-chloro-1-naphthol (4CN) substrate solution, and isopropanol (IPA) dropped onto bubble trap membranes illustrating 4CN beading atop the membrane similar to water and not wetting it, in contrast to IPA, which penetrates completely into the membrane and does not leave a droplet. Below the right hypothesis we show validation of all reagents used in the SiP sandwich assay for IL-8 detection (except the substrate solution) using an ELISA with enzymatic TMB readout. The bar graph plots the mean absorbance (error bars depicting one standard deviation), showing the expected dose-dependent binding and signal development with those reagents. **(c)** Results from a dot blot and corresponding densitometry quantification depict reduced signal development when the 4CN previously employed (‘Old’) is used as compared to freshly dissolved 4CN obtained in tablet form (‘New’), supporting the third hypothesis. Dots in the densitometry plot represent mean inverted grayscale image intensities over regions-of-interest outlined in red. **(d)** Representative silicon photonics assay results depicting use of the ‘New’ 4CN with substantially improved signal associated with the enzymatic amplification stage; dots represent calculated shifts based on pre- and post-stage buffer rinses with error bars depicting standard deviation across four sensor replicates. Individual lines in resonance peak shift vs. time sensorgrams depict sensors and colors represent channel replicates.

Initial experiments employing this protocol yielded resonance peak shifts across the enzymatic amplification assay stage in excess of 20 nm (example in Figure S3). In a later string of assays, however, we began observing markedly reduced amplification (representative example shown in Figure 4(a)).

At this juncture, the systematic troubleshooting framework that we felt was most applicable to identifying the cause of amplification signal loss was the ONC method [20] [33] [34], which is well-suited to scenarios like this one where a larger volume of potential hypotheses to explain a particular phenomenon may be present. We present an overview of the application of ONC to this challenge in Figure 4(b). A simple strategy is to identify any changes that were implemented before the issue appeared. These may range from deliberate protocol adjustments to simple aging of reagents and use of the testing system for other experiments. For this case of enzymatic amplification signal loss, we depict and discuss the four strongest hypotheses we generated (Figure 4(b)).

The first hypothesis focused on the microfluidic polytetrafluoroethylene (PTFE) bubble-traps (Precigenome, San Jose) that we had recently incorporated into our testing setup to mitigate bubble-related instability; we hypothesized that the hydrophobic PTFE membranes that enable gas transport out of fluidic lines may be being wetted by the enzymatic amplification substrate solution containing 17% methanol. This wetting process could result in depletion or mixing of the substrate before delivery to the HRP on the sensor surface, reducing its activity. The second hypothesis centered around the microfluidic PEEK tubing carrying assay reagents from fluidic reservoirs to the sensor surface; we considered that HRP-conjugated proteins used in our assays may have adsorbed onto the interior surface of this tubing in previous experiments, and that adsorbed HRP may have retained enough activity to deplete our 4CN substrate before it was able to react as intended at the sensor surface. Our third and fourth hypotheses were both based on degradation (e.g., due to age) of our reagents: the enzymatic substrate solution and the protein reagents, respectively.

The second stage of the ONC framework (Narrow) comprises filtering and triaging the list of generated hypotheses to most efficiently proceed with problem solving and testing. This efficiency of hypothesis testing is especially relevant to assay development for SiP sensors due to the degree of resource investment that such platforms can require per experiment in early R&D phases, which is even more critical in an academic research environment where resources are often limited. Assessing the likelihood that a given hypothesis is correct (in other words, how worthwhile resource investment into its testing may be) is a critical aspect of this troubleshooting framework. This process is aided considerably by incorporating insights from as many sources as possible: from less resource-intensive orthogonal assays to scientific principles and relevant literature.

We determined that it was unlikely that species adsorption onto tubing explained the stark difference in amplification between previous assays and what was now being observed; the absence of any evident gradual decay in binding signal over previous assays, coupled with the inclusion of BSA-blocking and post-assay multi-step fluidic system cleaning protocols led us to conclude this hypothesis was not worthwhile to prioritize for further testing. We further determined that the fourth hypothesis was unlikely to have been the cause of the near total loss of amplification signal through the use of orthogonal assays (ELISA). We regularly use ELISAs to characterize new products and lots of binding reagents (Figure 4(b), further details in SI Section 2.5 and SI Section 3), and we had recent data showing good activity of the same stocks of protein reagents when using another substrate (TMB) for readout.

The final stage of the ONC framework (Close) is testing those hypotheses determined to be likely root causes. To determine whether the recently incorporated bubble traps may be interacting with the methanol-containing substrate solution, we performed a crude contact angle assessment comparing water (which served as a negative control as it is confirmed to not wet the membrane and be fully compatible), IPA (which served as a positive control as it is confirmed to not be compatible), and the exact substrate solution delivered during our on-chip immunoassay. As depicted in the photograph in Figure 4(b), 4CN beads much like water on the PTFE bubble trap membrane and does not penetrate into the membrane like the IPA. This simple experiment showed that it is likely that the substrate solution is able to traverse the PTFE bubble trap membrane on its way to the SIP sensor in a manner similar to aqueous solution. This result suggested that membrane penetration was not the root cause of the low amplification signal. Finally, in the assessment of the substrate solution degradation hypothesis, we employed a modified dot blot protocol (details in SI Section 2.4) to isolate and test the 4CN substrate solution. First, the HRP-conjugated secondary antibody used in the silicon photonics assay was dropped onto PVDF membranes in triplicate and at two concentrations. After an incubation and wash, the membranes were submerged in equal volumes of either the 4CN we had so far been using, which was obtained and stored as recommended by the vendor in liquid format for three-to-four months into its one-year guaranteed period (‘Old’), or freshly obtained and dissolved 4CN originating in solid/tablet form (‘New’). As illustrated in Figure 4(c), densitometry analysis indicated an approximately five-fold increase in reaction product development when comparing the ‘New’ 4CN with the ‘Old’; when the ‘New’ 4CN was then employed on the silicon photonics platform, signal was recovered to the magnitudes expected from our previous assays prior to encountering this issue Figure 4(d).

In summary, using a structured problem-solving methodology enabled us to navigate an especially complex and multifaceted technical workflow to identify the source of a critical issue hampering experimental progress. Just as importantly, it helped us do so in a more streamlined fashion that reduced troubleshooting time and wasted effort and resources. The utility of employing orthogonal assays is particularly pronounced here, both for efficient resource allocation but also by allowing researchers to isolate variables – the significance of which cannot be underscored enough for these types of label-free biosensor assays, where the outcome of an individual experiment is subject to myriad influences.

### 3.4 Elucidating contributors to variability using fishbone analysis

Even after implementing the workflow changes described in Section 3.3, we continued to observe high inter-assay variability in our enzymatic amplification signal (Figure 5(a)). Because many factors can contribute to variability in SiP (and other label-free) biosensing assays [10], we applied an established graphical framework (the fishbone diagram) to systematically map these factors (Figure 5(b)). This framework provided an organized structure to guide root cause analysis by highlighting interdependencies across assay components, supporting both identification and prioritization of high-impact contributors. The fishbone diagram particularly serves as an excellent tool for the “open” phase of ONC by helping to visualize all potential sources of variability before targeted investigation.

**Figure 5.**
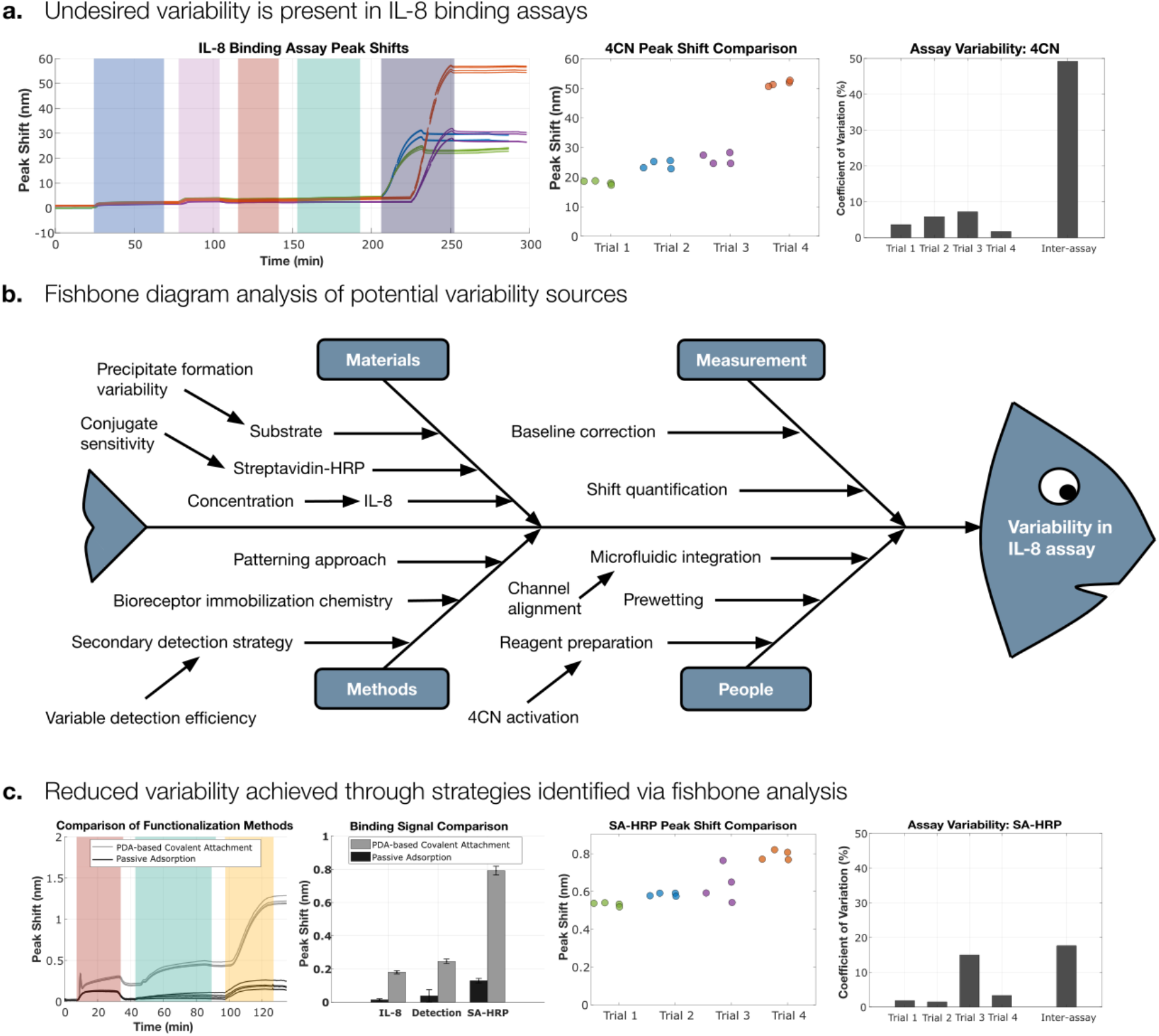
Use of a fishbone diagram to illustrate potential sources of signal variability in the IL-8 binding assay and help identify strategies to improve replicability. **(a)** High inter-assay variability (inter-assay CV 49.2%) was observed at the enzymatic amplification stage of IL-8 assays, motivating investigation into contributing factors. Line colours (green, blue, purple, orange) represent different channel replicates, allowing visualization of replicate-specific sources of variability. **(b)** A fishbone diagram analysis was performed to map hypothesized sources of unexpected variability across major assay components between consistent protocols. Branches represent underlying problems that could contribute to inconsistent signals, such as bioreceptor immobilization chemistry (e.g., weak or variable binding with passive adsorption), patterning approach (e.g., variability introduced by flow-mediated functionalization), secondary detection strategy (e.g., reduced activity from direct HRP conjugation), precipitate formation (e.g., differing substrate kinetics or stability across experiments), and 4CN activation (e.g., inconsistencies in in-house tablet preparation and activation with H_2_O_2_). These branches do not represent that different protocols were applied within replicates; rather, they reflect hypothesized mechanisms by which a given protocol (e.g., passive adsorption) could produce variability. **(c)** Mitigation strategies were implemented to increase assay signal without the use of enzymatic amplification: (1) using a biotinylated detection antibody with SA-HRP in place of the HRP-conjugated detection antibody, and (2) employing polydopamine (PDA)-based covalent capture antibody spotting instead of flow-based passive adsorption. (Left) The comparison of functionalization methods includes two peak shift plots: the grey lines represent PDA-based covalent capture antibody spotting, while the black lines represent flow-based passive adsorption. The first step in the peak shift plot corresponds to IL-8 binding (red overlay), the second to detection antibody (teal overlay), and the third to SA-HRP (yellow overlay). (Left-middle) The bar graph shows that the PDA-based approach produced higher peak shifts, allowing more reliable quantification. (Right-middle) Notably, the SA-HRP shifts across four trials were consistently above 0.5 nm, eliminating the need for enzymatic amplification. (Right) Overall, combining PDA-based capture antibody immobilization with biotinylated detection and SA-HRP reduced inter-assay variability (inter-assay CV 17.7%) relative to pre-optimization assays and removed the requirement for enzymatic amplification.

Whereas frameworks such as the Five Whys and PDCA are often used to resolve single-point failures, the fishbone diagram allowed us to consider and test a wider range of potential contributors that may work together to cause the problem. In our case, the fishbone diagram was simplified from the conventional six categories (materials, methods, people, environment, measurement and equipment) to four categories (materials, methods, people and measurement), focusing the analysis on the most relevant sources of variability in our setup (SI Section 5). We explored the use of the Five Whys, PDCA, and ONC to probe potential causes in greater depth, adapting the choice of framework to where it was most informative.

We first attempted to isolate variability by assay stage, testing a higher IL-8 concentration (10 µg/mL) with the goal to quantify binding at the capture antibody, IL-8, detection antibody, and enzymatic amplification stages to help identify at which assay stage the variability originated. However, at that time the IL-8 signal still remained below the quantifiable limit. Despite this, variability was observed across the two channel-replicates at both the HRP-conjugated detection antibody and enzymatic amplification stages, indicating that amplification was not solely responsible (Figure S6).

From here, we hypothesized that inter-experimental variability in nonspecific signal contributed substantially to overall variability. To investigate, we directly tested for nonspecific signal on-chip and confirmed its presence (Figure S7). To further dissect this effect, we designed an ELISA comparing multiple blocking strategies and conditions, using both the HRP-conjugated antibody originally employed in our assays and a biotinylated detection antibody matched with our capture antibody in a vendor-provided ELISA kit. This comparison revealed that the biotinylated antibody, when paired with a SA-HRP step, produced a stronger signal with lower nonspecific binding (Figure S4). Based on these findings, we pivoted to using the biotinylated antibody in our IL-8 assays.

This adjustment to the assay design exemplifies how the Five Whys guided our RCA process. Regular variability in the enzymatic amplification stage prompted us to ask:

- Why is there variability? One contributor may be nonspecific binding of the detection antibody.
- Why is there non-specific binding? Poor compatibility between the antibody pair.
- Why is the compatibility poor? Inadequate validation within our assay context.
- Why was validation inadequate? Absence of systematic vendor correspondence and pre-screening of matched pairs during assay design.

Addressing this root cause led us to implement two key changes for our future assay development: (i) correspondence with vendors regarding matched antibody pairs during assay design, and (ii) pre-validation of antibody pairs in ELISA format (specifically testing for nonspecific signal) prior to transfer into the on-chip assay.

Despite the switch to the more compatible biotinylated-antibody and SA-HRP, inter-assay variability remained high (CV = 77%, n = 4 channel replicates, Figure S8). We therefore turned to strategies from the literature and implemented a series of iterative PDCA cycles to further mitigate variability.

PDCA 1: Because passive adsorption-mediated bioreceptor attachment can suffer from instability and does not permit regeneration (bioreceptors are stripped during harsh regeneration conditions), we adopted a more robust covalent functionalization strategy leveraging polydopamine [35]. Covalent attachment has been reported to enhance sensor stability and reproducibility [35], [36], and thus was expected to improve assay replicability. Concurrently, we sought to reduce variability introduced by substrate preparation and handling by testing a commercial 1-Step 4CN substrate solution (SI Section 2), a stabilized, ready-to-use formulation containing both 4CN and hydrogen peroxide; this substrate has been previously used in SiP assays [37]. This eliminated the need to pause the assay for 4CN activation, block channels, and exchange freshly prepared substrate, thereby minimizing user-dependent variability. This pilot assay produced higher signals at all stages, enabling easier quantification. However, the surface could not be effectively regenerated (92% signal loss between the first and second assay rounds; 0.4% regeneration efficiency, Figure S9).

PDCA 2: Seeking to facilitate regeneration, we next tested a 1-Step precipitate-forming TMB-blotting substrate solution (in contrast to conventional soluble TMB formulations). Although supplied ready-to-use and not requiring hydrogen peroxide addition, it was not pursued further due to the poor amplification signal observed (Figure S9). However, as with the 1-Step 4CN trial, after implementing covalent functionalization, the SA-HRP stage showed a ~5-fold increase relative to passive adsorption (Figure 5(c) left-middle), and all assay stages produced quantifiable signal.

PDCA 3: Based on this outcome, we proceeded to use SA-HRP as the final signal amplification step in combination with the sandwich assay. This approach avoided the regeneration challenges associated with enzymatic amplification (Figure S9). Guided by these findings, we implemented PDA-based antibody immobilization together with a biotinylated detection antibody and SA-HRP, as the need for enzymatic amplification was effectively eliminated. In this optimized format, the SA-HRP shift alone provided ~4× amplification of the IL-8 binding signal (Figure 5(c), middle), yielding sufficient signal to reliably detect the target concentration (3.125 ng/mL), while simplifying the workflow and reducing variability from an inter-assay CV of 49.2% to 17.7% (Figure 5(a) & 5(c), right).

These results demonstrate how the fishbone diagram guided a systematic root cause analysis, enabling targeted interventions to mitigate inter-assay variability. Implementing a high-level ONC strategy, we leveraged the fishbone diagram for the “open” phase to map potential sources of variability, and applied the Five Whys to pinpoint causes of nonspecific binding followed by iterative PDCA cycles to optimize substrate handling, bioreceptor immobilization, and signal amplification during the “narrow” and “close” phases. Together, these interventions simplified the assay, reduced variability, and enabled robust, quantifiable IL-8 detection, highlighting the utility of structured RCA frameworks for enhancing complex biosensing assays.

## 4. Conclusion

Assay failure is an inevitable part of the assay development process, particularly in interdisciplinary environments like academic laboratories that conduct novel research as well as training of highly qualified personnel. Rather than treating failed experiments as wasted effort, our experience demonstrates that developing a simple, three-question workflow for continuous improvement and systematically applying troubleshooting strategies and RCA frameworks can transform setbacks into valuable learning opportunities. By embedding structured reflection into every experimental cycle, we have been able to iteratively refine our assays and our team’s collective practices.

The implementation of our workflow has had a measurable impact on our team’s assay yield and throughput. For example, in early binding assays performed before implementing this process, only ~45% of assays produced high quality, reliable data (n = 29 assays from 2021 until August 2023). In contrast, after introducing structured RCA approaches and iteratively improving our methods, our assays achieved a success rate of ~90% (n = 22 assays in summer 2025), defined as experiments that executed successfully and produced resulting data that were reliable and within expected ranges.

Through our case studies, we also reflected on the relative strengths of the different frameworks incorporated into our workflow. The Five Whys is a straightforward, effective first-line RCA framework that is lightweight and intuitively guides researchers beyond simple problem-solving to find the underlying root cause of the issue. In the example that we shared, simply replacing the malfunctioning valve would have solved the problem, but digging deeper to consult with the vendor, analyze our protocols, and update our training, documentation, and analysis approaches allowed us to mitigate and reduce the impact of future equipment failures. The Plan-Do-Check-Act cycle for continuous improvement invites researchers to take a series of iterative small steps to solve problems and improve experiments. The Open-Narrow-Close framework is well-suited to situations where multiple interacting variables could plausibly contribute to assay failure, providing structure to hypothesis generation and prioritization. Finally the fishbone diagram is most valuable when helping visualize system-wide interdependencies and stimulate broad, inclusive problem-solving. Together, these methods provide a flexible toolkit, allowing us to select the right framework or combination of frameworks depending on the nature and complexity of the problem. To aid researchers in selecting an appropriate set of frameworks to address a given problem, Figure 6 presents an example of how all four frameworks can be used together and summarizes the relative strengths of each framework.

**Figure 6.**
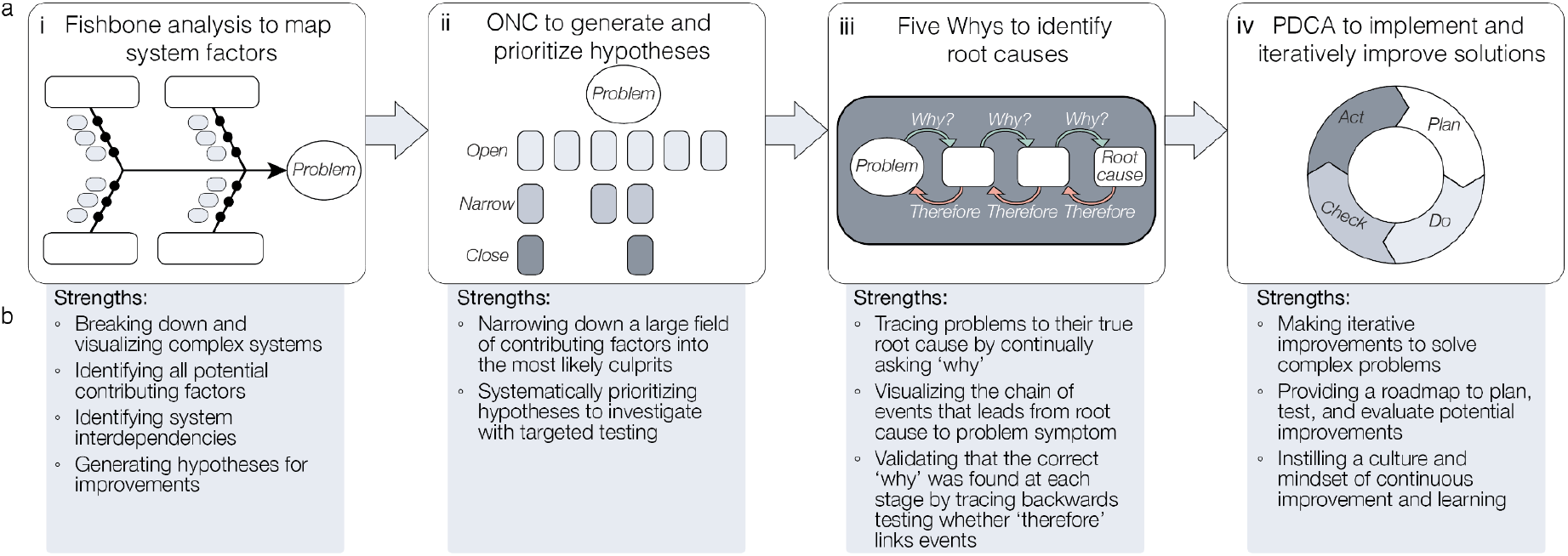
Comparing and combining the frameworks described here. **(a)** A potential RCA process flow combining all four frameworks described here. **(b)** Summary of the strengths of each framework to aid in choosing the appropriate framework or set of frameworks for a given problem.

Despite the benefits, our workflow has limitations. The specific system components, logging strategies and troubleshooting priorities we describe are tailored to silicon photonic biosensors and our specific testing setup, and translation to other platforms will require adaptation. We demonstrate the integration of four example frameworks for problem-solving and RCA in this work; however, future work will involve continuing to iteratively improve the workflow, for example by implementing additional methods and frameworks into the third stage of our workflow (e.g., Fault Tree Analysis (FTA) [38], [39] and Failure Modes and Effects Analysis (FMEA) [40] [41]). Furthermore, it is worth highlighting that although lightweight adaptations of these frameworks lower barriers to adoption and can provide considerable benefit, they cannot replace more formalized RCA methods and experimental design methodologies such as Design of Experiments (DOE).

Looking forward, there are several opportunities to further improve reproducibility, collaboration, and efficiency in biosensor assay development. While we report a simple, document-based method for designing and summarizing experiments, building in the use of standardized protocol repositories like protocols.io [42] or electronic lab notebooks [43] could enhance continuity across research teams and projects and promote replicability, data integrity, and the use of data management best practices. Integration of DOE principles [44] into assay optimization could provide more rigorous, statistically guided improvements. By building on the foundations of lightweight troubleshooting and RCA, these additional tools could help academic teams continue to bridge the gap between exploratory research environments and the robustness of industrial workflows.

Label-free biosensor assay development is a complex, interdisciplinary challenge in which things will go wrong: equipment will fail, researchers will make mistakes, and reagents will not perform as expected. This work illustrates how reframing assay failure as a structured learning opportunity can drive continuous improvement in assay yield and throughput by mitigating both the probability of these failures and their impacts. In designing our workflow, we noted that an important consideration for future adoption is researcher engagement and buy-in. For trainees eager to move quickly, detailed logging and structured troubleshooting may initially feel unnecessary, especially when their assays seem to be working. In academic settings, the culture often emphasizes innovation, while the challenges of achieving reproducibility and manufacturability at scale are seen as problems for industry. Yet, as we show in these examples, these practices can provide immediate value by accelerating troubleshooting when challenges inevitably arise, saving time and effort in the long run. At the same time, introducing such habits early also prepares trainees for the rigor of translational or industrial environments, where reproducibility and reliability are paramount. It further readies the research itself for translation and collaboration with industry, accelerating the impact of the work. Our lightweight frameworks are intended to support both goals, helping students advance their current projects more effectively while also building skills that will serve them well in future careers, but their success ultimately depends on students embracing these practices as part of their research process. We have navigated this balance by prioritizing simplicity and accessibility in the development of our workflow and shared specific examples of its benefits. By sharing both our challenges and the solutions we developed, we hope to contribute to a culture of continuous improvement in academic research (and other resource-limited environments like startup companies) by providing a roadmap for other teams navigating the complexities of label-free biosensor assay development to find the best balance of workflows for their teams.

## Supporting information

Supplemental protocol for IL-8 assay

Supplemental reagent prep sheet for IL-8 assay

## Author contributions

**Conceptualization**, S.C., S.M.G., A.R., K.C.C.; **Data curation**, S.C., S.M.G., A.R., M.W., K.N.; **Formal analysis**, S.C., S.M.G., A.R., M.W., K.N.; **Funding acquisition**, S.M.G., L.C., S.S., and K.C.C.; **Investigation**, S.C., S.M.G., A.R., M.W., K.N., L.P., Y.H.; **Methodology**, S.C., S.M.G., A.R., M.W., K.N., L.P., Y.H.; **Project administration**, S.C., S.M.G., L.C., S.S., K.C.C.; **Software**, S.M.G. and A.R.; **Resources**, S.M.G., L.C., S.S., and K.C.C.; **Supervision**, S.M.G., A.R., L.C., S.S., and K.C.C.; **Validation**, S.C., S.M.G., A.R., M.W., K.N.; **Visualization**, S.C., S.M.G., A.R., M.W., K.N.; **Writing - original draft**, S.C., S.M.G., A.R., M.W., K.N.; **Writing - review & editing**, S.C., S.M.G., A.R., M.W., K.N., L.P., L.C., S.S., K.C.C..

## Data availability

Data will be made available upon request.

## Acknowledgements

The authors are grateful to work and live on land that is the traditional, ancestral, and unceded territory of the Coast Salish Peoples, including the territories of the Musqueam, Squamish, and Tsleil-Waututh First Nations. This manuscript reports the results of several years of improvements in our team’s experimental workflows, which were enabled by all members of our interdisciplinary team. The authors would like to express their immense gratitude to all of the members of our biosensors team from 2021-2025, whose work, feedback, and suggestions made these improvements possible. The authors acknowledge specific contributions from Mohammed Al-Qadasi, for the suggestion to include the table to log our experiment checks within our experimental protocol documents, and from So Jung Kim, for suggestions for specific checks to be included. The authors are also grateful for assistance with assay preparation and execution from Stephen Kioussis, for data acquisition and analysis software contributions from Yifei Liu, Piramon Tisapramotkul, and Mohammed Al-Qadasi, as well as for contributions to photonic and microfluidic design and fabrication from Ben Cohen-Kleinstein, Dr. Matthew Mitchell, and Kowsar Heydari. The authors acknowledge project and fellowship support from the Schmidt Science Polymaths Program, Optica Foundation Challenge (026724), School of Biomedical Engineering Trainee Research Awards program, Mitacs Inc. (IT30425), UBC Expansion Funding for Canada’s Digital Technology Supercluster (CDTS) projects, Natural Sciences and Engineering Research Council of Canada (NSERC) (RGPIN-2020-04798), Killam Trusts, Silicon Electronics-Photonics Integrated Circuits Fabrication (SIEPICfab) consortium, CMC Microsystems, and Canada’s National Design Network. We also acknowledge undergraduate research funding supplements from the UBC Work Learn International Undergraduate Research Awards, Faculty of Medicine Multidisciplinary Research Program in Medicine, Centre for Blood Research Undergraduate Summer Research Program, NSERC Undergraduate Student Research Awards, and School of Biomedical Engineering Synergy Research Program. The authors acknowledge contributions from Applied Nanotools Inc., the Stewart Blusson Quantum Matter Institute Advanced Nanofabrication Facility and machine shop, the research lab of Prof. Hugh Kim (including Steven Zhexuan Jiang for assistance with dot blot and ELISA readout), Kizhakkedathu Research Group, Bizzotto Research Lab, and Overall Lab.

## Supplementary information

Supplementary documents are attached separately.

## References

[1] A. Chieng, Z. Wan, and S. Wang, “Recent Advances in Real-Time Label-Free Detection of Small Molecules.,” Biosensors (Basel), vol. 14, no. 2, Feb. 2024, doi: 10.3390/bios14020080.

[2] M. Nirschl, F. Reuter, and J. Vörös, “Review of transducer principles for label-free biomolecular interaction analysis.,” Biosensors (Basel), vol. 1, no. 3, pp. 70–92, Jul. 2011, doi: 10.3390/bios1030070.

[3] A. Bonyár, “Label-Free Nucleic Acid Biosensing Using Nanomaterial-Based Localized Surface Plasmon Resonance Imaging: A Review,” ACS Appl. Nano Mater., vol. 3, no. 9, pp. 8506–8521, Sep. 2020, doi: 10.1021/acsanm.0c01457.

[4] A. L. Washburn, L. C. Gunn, and R. C. Bailey, “Label-free quantitation of a cancer biomarker in complex media using silicon photonic microring resonators.,” Anal. Chem., vol. 81, no. 22, pp. 9499–9506, Nov. 2009, doi: 10.1021/ac902006p.

[5] E. Luan, H. Shoman, D. M. Ratner, K. C. Cheung, and L. Chrostowski, “Silicon Photonic Biosensors Using Label-Free Detection.,” Sensors, vol. 18, no. 10, Oct. 2018, doi: 10.3390/s18103519.

[6] C. Dhote, A. Singh, and S. Kumar, “Silicon Photonics Sensors for Bio-Photonic Applications - A Review,” IEEE Sens. J., pp. 1–1, 2022, doi: 10.1109/JSEN.2022.3199663.

[7] E. Zeidan, C. L. Kepley, C. Sayes, and M. G. Sandros, “Surface plasmon resonance: a label-free tool for cellular analysis.,” Nanomedicine (Lond), vol. 10, no. 11, pp. 1833–1846, 2015, doi: 10.2217/nnm.15.31.

[8] C. V. Agu, R. L. Cook, W. Martelly, L. R. Gushgari, M. Mohan, and B. Takulapalli, “Multiplexed proteomic biosensor platform for label-free real-time simultaneous kinetic screening of thousands of protein interactions.,” Commun. Biol., vol. 8, no. 1, p. 468, Mar. 2025, doi: 10.1038/s42003-025-07844-z.

[9] T. S. Bronder, A. Poghossian, M. P. Jessing, M. Keusgen, and M. J. Schöning, “Surface regeneration and reusability of label-free DNA biosensors based on weak polyelectrolyte-modified capacitive field-effect structures.,” Biosens. Bioelectron., vol. 126, pp. 510–517, Feb. 2019, doi: 10.1016/j.bios.2018.11.019.

[10] L. S. Puumala et al., “Resonating with replicability: factors shaping assay yield and variability in microfluidics-integrated silicon photonic biosensors,” BioRxiv, Jul. 2025, doi: 10.1101/2025.07.16.664198.

[11] M. Barsalou, “Determining which of the classic seven quality tools are in the quality practitioner’s RCA tool kit,” Cogent Engineering, vol. 10, no. 1, Dec. 2023, doi: 10.1080/23311916.2023.2199516.

[12] D. Tahmasbi, H. Shirali, S. Sajad Mousavi Nejad Souq, and M. Eslampanah, “Diagnosis and root cause analysis of bearing failure using vibration analysis techniques,” Eng. Fail. Anal., vol. 158, p. 107954, Apr. 2024, doi: 10.1016/j.engfailanal.2023.107954.

[13] G. Singh, R. H. Patel, S. Vaqar, and J. Boster, “Root cause analysis and medical error prevention,” in StatPearls, Treasure Island (FL): StatPearls Publishing, 2025.

[14] S. Shin, D. Kang, and R. S. Walmsley, “Post-endoscopy esophageal adenocarcinoma and root cause analysis in Auckland, New Zealand,” Endosc. Int. Open, Aug. 2025, doi: 10.1055/a-2676-3883.

[15] G. Aryaduta and R. Rusindiyanto, “Risk mitigation strategy for CNC plasma cutting machine damage using the house of risk method and root cause analysis,” JLMA, vol. 6, no. 4, pp. 951–969, Aug. 2025, doi: 10.37899/journallamultiapp.v6i4.2202.

[16] K. B. Percarpio, B. V. Watts, and W. B. Weeks, “The effectiveness of root cause analysis: what does the literature tell us?,” Jt Comm J Qual Patient Saf, vol. 34, no. 7, pp. 391–398, Jul. 2008, doi: 10.1016/S1553-7250(08)34049-5.

[17] O. Serrat, “The five whys technique,” in Knowledge Solutions, Singapore: Springer Singapore, 2017, pp. 307–310.

[18] N. Tague, The Quality Toolbox, 2nd ed. ASQ Quality Press, 2004, p. 585.

[19] M. Best and D. Neuhauser, “Walter A Shewhart, 1924, and the Hawthorne factory.,” Qual. Saf. Health Care, vol. 15, no. 2, pp. 142–143, Apr. 2006, doi: 10.1136/qshc.2006.018093.

[20] Interaction Institute for Social Change, “Core Models of Facilitative Research,” 2014, Accessed: Sep. 09, 2025. [Online]. Available: http://tiach.pbworks.com/w/file/fetch/90446663/Handout%20-%20Facilitative%20Leadership%2012.9.14.pdf.

[21] A. Kumah et al., “Cause-and-Effect (Fishbone) Diagram: A Tool for Generating and Organizing Quality Improvement Ideas.,” Glob. J. Qual. Saf. Healthc., vol. 7, no. 2, pp. 85–87, May 2024, doi: 10.36401/JQSH-23-42.

[22] K. Ishikawa, Guide to Quality Control, 2nd ed. Tokyo, 1986.

[23] L. May and T. Runyon, “Labscrum: A case study for agility in academic research labs,” May 2019, doi: 10.31234/osf.io/zg4ub.

[24] International Baccalaureate Organization, “Middle Years Programme Sciences Guide,” International Baccalaureate Organization (UK) Ltd, 2019.

[25] D. A. Kolb, “Experiential learning: Experience as the source of learning and development,” J. Organ. Behav., vol. 8, no. 4, pp. 21–38, 1984, doi: 10.1002/job.4030080408.

[26] T. H. Morris, “Experiential learning – a systematic review and revision of Kolb’s model,” Interactive Learning Environments, vol. 28, no. 8, pp. 1064–1077, Jan. 2019, doi: 10.1080/10494820.2019.1570279.

[27] E. Ries, “The Five Whys for Start-Ups,” 2010. https://hbr.org/2010/04/the-five-whys-for-startups (accessed Sep. 10, 2025).

[28] X. Zhong et al., “A descriptive study on clinical department managers’ cognition of the Plan-Do-Check-Act cycle and factors influencing their cognition.,” BMC Med. Educ., vol. 23, no. 1, p. 294, May 2023, doi: 10.1186/s12909-023-04293-2.

[29] M. S. Luchansky, A. L. Washburn, M. S. McClellan, and R. C. Bailey, “Sensitive on-chip detection of a protein biomarker in human serum and plasma over an extended dynamic range using silicon photonic microring resonators and sub-micron beads.,” Lab Chip, vol. 11, no. 12, pp. 2042–2044, Jun. 2011, doi: 10.1039/c1lc20231f.

[30] B. S. Munge et al., “Nanostructured immunosensor for attomolar detection of cancer biomarker interleukin-8 using massively labeled superparamagnetic particles.,” Angew. Chem. Int. Ed, vol. 50, no. 34, pp. 7915–7918, Aug. 2011, doi: 10.1002/anie.201102941.

[31] J. T. Kindt, M. S. Luchansky, A. J. Qavi, S.-H. Lee, and R. C. Bailey, “Subpicogram per milliliter detection of interleukins using silicon photonic microring resonators and an enzymatic signal enhancement strategy.,” Anal. Chem., vol. 85, no. 22, pp. 10653–10657, Nov. 2013, doi: 10.1021/ac402972d.

[32] G. Volpin, M. Cohen, M. Assaf, T. Meir, R. Katz, and S. Pollack, “Cytokine levels (IL-4, IL-6, IL-8 and TGFβ) as potential biomarkers of systemic inflammatory response in trauma patients.,” Int. Orthop., vol. 38, no. 6, pp. 1303–1309, Jun. 2014, doi: 10.1007/s00264-013-2261-2.

[33] L. A. Zampetakis, L. Tsironis, and V. Moustakis, “Creativity development in engineering education: the case of mind mapping,” Journal of Mgmt Development, vol. 26, no. 4, pp. 370–380, Apr. 2007, doi: 10.1108/02621710710740110.

[34] P. A. Heslin, “Better than brainstorming? Potential contextual boundary conditions to brainwriting for idea generation in organizations,” J. Occup. Organ. Psychol., vol. 82, no. 1, pp. 129–145, Mar. 2009, doi: 10.1348/096317908X285642.

[35] S. Bakshi et al., “Bio-inspired polydopamine layer as a versatile functionalisation protocol for silicon-based photonic biosensors.,” Talanta, vol. 268, no. Pt 1, p. 125300, Feb. 2024, doi: 10.1016/j.talanta.2023.125300.

[36] L. S. Puumala et al., “Biofunctionalization of multiplexed silicon photonic biosensors.,” Biosensors (Basel), vol. 13, no. 1, Dec. 2022, doi: 10.3390/bios13010053.

[37] J. H. Wade, A. T. Alsop, N. R. Vertin, H. Yang, M. D. Johnson, and R. C. Bailey, “Rapid, multiplexed phosphoprotein profiling using silicon photonic sensor arrays.,” ACS Cent. Sci., vol. 1, no. 7, pp. 374–382, Oct. 2015, doi: 10.1021/acscentsci.5b00250.

[38] C. G. Siontorou and F. A. Batzias, “A methodological combined framework for roadmapping biosensor research: a fault tree analysis approach within a strategic technology evaluation frame.,” Crit. Rev. Biotechnol., vol. 34, no. 1, pp. 31–55, Mar. 2014, doi: 10.3109/07388551.2013.790339.

[39] C. S. Carlson, “Introduction to fault tree analysis (FTA),” in Effective FMEAs: achieving safe, reliable, and economical products and processes using failure mode and effects analysis, Wiley, 2012, pp. 316–327.

[40] R. J. Mikulak, R. McDermott, and M. Beauregard, The Basics of FMEA, 2nd ed. Productivity Press, 2017, p. 90.

[41] J. F. van Leeuwen et al., “Risk analysis by FMEA as an element of analytical validation.,” J. Pharm. Biomed. Anal., vol. 50, no. 5, pp. 1085–1087, Dec. 2009, doi: 10.1016/j.jpba.2009.06.049.

[42] L. Teytelman, A. Stoliartchouk, L. Kindler, and B. L. Hurwitz, “Protocols.io: virtual communities for protocol development and discussion.,” PLoS Biol., vol. 14, no. 8, p. e1002538, Aug. 2016, doi: 10.1371/journal.pbio.1002538.

[43] S. G. Higgins, A. A. Nogiwa-Valdez, and M. M. Stevens, “Considerations for implementing electronic laboratory notebooks in an academic research environment.,” Nat. Protoc., vol. 17, no. 2, pp. 179–189, Feb. 2022, doi: 10.1038/s41596-021-00645-8.

[44] Jiju. Antony, Design of experiments for engineers and scientists. Oxford, 2003.

