## Supplementary figures for "What Could Go Wrong? Promoting Success by Planning for Failure in Label-Free Biosensor Assay Development"

#### 1. Experimental cycle

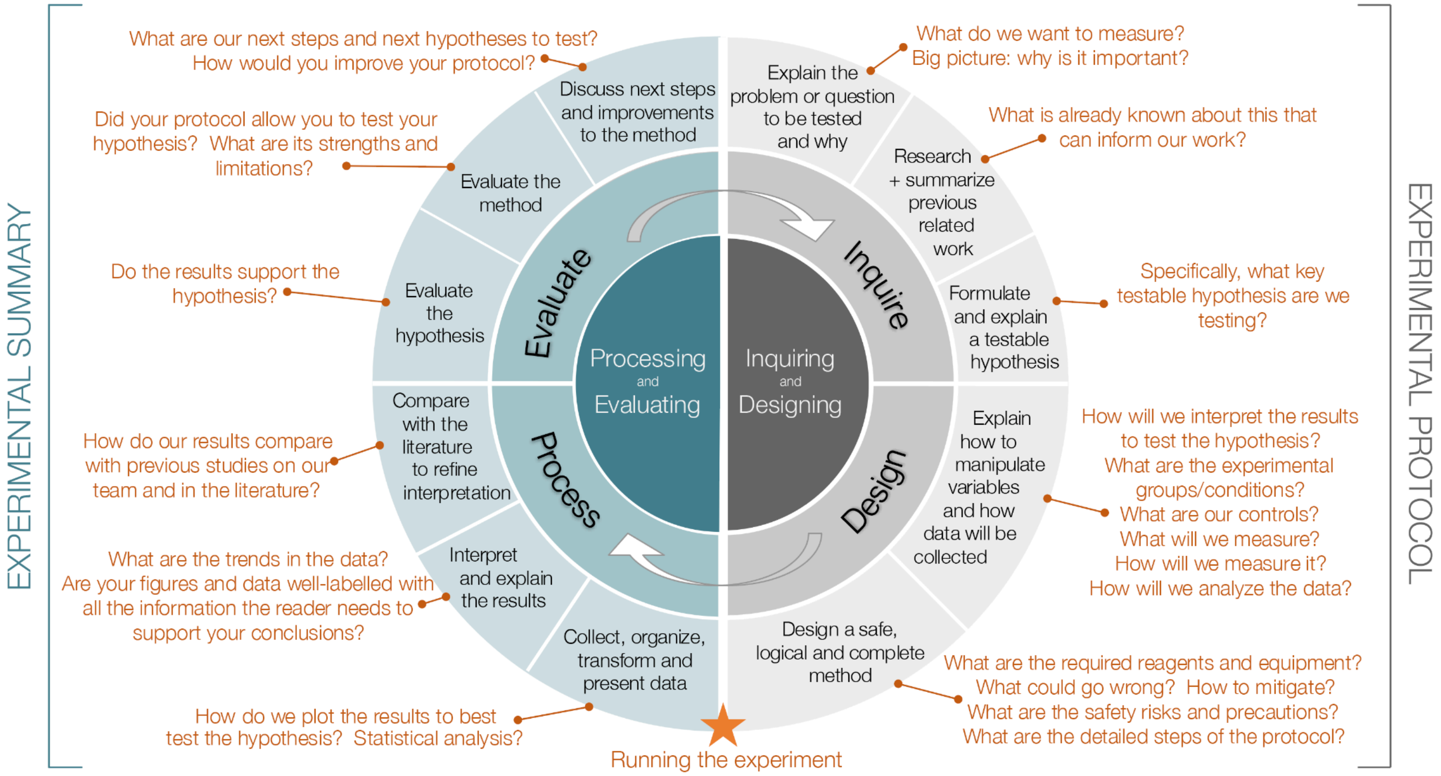


**Figure S1**. Overview of the experimental cycle, probing questions, and documentation that our team uses for each experiment as a part of our workflow. This cycle diagram was adapted by our team using resources provided by i-Biology.net and International Baccalaureate [1], [2].

#### 2. Technical methods

##### 2.1 On-chip binding assays

To detect IL-8, our team used a sandwich assay format similar to that previously described by Kindt, *et al*. [3]. Our assay involved: (1) functionalizing the SiP sensors using monoclonal antibodies; (2) delivering a sample containing the IL-8 cytokine, which binds to the capture antibody on the sensor surface; (3) binding a detection antibody (either biotinylated or horseradish peroxidase (HRP)-conjugated) to the bound IL-8; (4) (if a biotinylated detection antibody was used) binding streptavidin-HRP (SA-HRP) to the sensor surface. We implemented the assay using a photonic-fluidic testing setup and methods as previously described [4]. This work employed photonic circuits containing SWG ring resonator biosensors with ring radius $R$ = 30 µm, coupling gaps $g_c$ = [500, 550] nm, grating period $\Lambda$ = 250 nm, duty cycles $\delta$ = [0.65, 0.7], waveguide width $w$ = 500 nm, and waveguide thickness $t$ = 220 nm, as previously described [5]. The photonic circuits were fabricated on silicon-on-insulator (SOI) wafers by Applied Nanotools, Inc. (ANT, Edmonton, AB, Canada) using their NanoSOI Fabrication Service Silicon Device Layer process (employing 100 keV electron beam lithography and reactive ion etching [6]). The fabricated photonic circuits were integrated with microfluidic gaskets fabricated by casting poly(dimethylsiloxane) (PDMS) in 3D-printed moulds (using a ProFluidics 285D printer and MasterMould resin (CadWorks 3D) as previously described [4], [5], [7]).

For this work, the capture antibody was either (1) passively adsorbed to the sensor surface using flow-mediated functionalization at a concentration of 20 µg/mL in PBS or (2) covalently attached to the surface using a PDA-mediated functionalization approach involving pipette spotting as previously described, at a concentration of 240 µg/mL [4]. All remaining assay stages were delivered through the microfluidic device under constant flow at 20-30 µL/min using a Fluigent LineUp fluidic control system as previously described [4]. In several assays using covalent antibody attachment, we performed sensor regeneration using flow of a pH 2.2 glycine-HCl solution [8]–[11], disrupting the antibody-antigen interaction and allowing the sensor to be used for multiple rounds of binding.

The on-chip IL-8 immunoassays described here used the following specific reagents: IL-8 capture antibody (included in ELISA kit R&D Systems DY208-05, lots ASJ3822051, ASJ3824031); IL-8 (included in ELISA kit R&D Systems DY208-05, lots ASJ3822051, ASJ3824031); IL-8 detection antibody (R&D Systems BAF208, lots UM2322052, UM221031), high-sensitivity streptavidin-HRP (Pierce 21130 lot XJ361814). For assays using covalent capture antibody attachment, an initial rinse with an acidic sensor regeneration buffer was used to remove non-covalently attached functionalization and blocking reagents from the sensor surface prior to the binding assay. This regeneration buffer included final concentrations of 10 mM glycine (Sigma-Aldrich G8790-100G, lot SLCG2930) and 160 mM NaCl (Fisher S271-3, lot 166221), adjusted to pH 2.2 using 1M HCl stock (prepared using ultrapure water and HCl concentrate: Sigma-Aldrich 258148-2.5L-GL, PCode: 4102988462, Source: MKCS8144), with balance of ASTM Type I water from a Nanopure Diamond system.

An overview of the stages of the IL-8 binding assay without enzymatic amplification described here is presented in Figure S2 below.

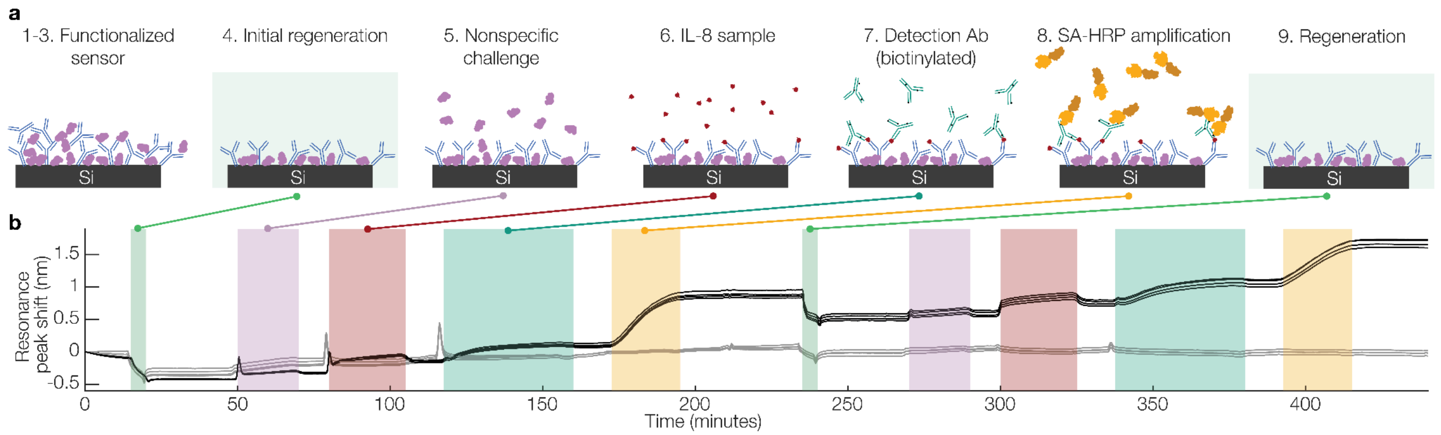


**Figure S2**. Overview of the on-chip sandwich immunoassay developed to detect IL-8. **(a)** cross-sectional views of the sensor waveguides depicting the assay stages. (1-3) the sensor surface is first functionalized with capture antibody and blocked with BSA, either using a flow-mediated passive adsorption process or a covalent attachment process using polydopamine-mediated attachment and pipette spotting. (4) an initial rinse with acidic regeneration buffer removes non-covalently attached antibody and blocking agent. (5) A 1 mg/mL solution of BSA in PBS is delivered to the sensor to test for nonspecific binding. (6) a sample made up of IL-8 in assay running buffer (either 0 or 3.125 ng/mL) is delivered to the sensor, and IL-8 specifically attaches to the capture antibody. (7) a biotinylated IL-8 detection antibody (1 µg/mL in assay running buffer) is delivered to the sensor surface and binds to the immobilized IL-8. (8) streptavidin-horseradish peroxidase (SA-HRP) (2 µg/mL in assay running buffer) is delivered to the surface and amplifies the sandwich assay signal. (9) the sensor is regenerated using a pH 2.2 glycine-HCl solution to disrupt the antibody-antigen interaction and prepare for another binding cycle. Between each assay stage the sensor is rinsed with assay running buffer (0.1 mg/mL BSA in PBS, 30 µL/min). **(b)** example sensorgram depicting the stages of the IL-8-detection immunoassay. Two cycles of detection are shown. The sensors in a microfluidic channel to which 3.125 ng/mL IL-8 was delivered during the sample stage are coloured black, while those in a negative control channel (0 ng/mL IL-8) are coloured grey.

#### 2.2 Leakage testing

The 10-port microfluidic selector valve is a key component of the fluidic control system used in our assays (Fluigent LineUp including the M-Switch selector valve). It is an 11-port/10-position microfluidic bidirectional valve that our team uses for delivery of different solutions to our chip and permits selection from 10 input fluidic reservoirs for delivery. The peripheral ports are numbered from 1 to 10 and are connected to one central port, and the switch is actuated by a motor that drives a rotor [12]. To test for leakage from the valve, our team developed a protocol in which red food dye (20% Scott-Bathgate Red Food Dye in ultrapure water) was placed in one of the ten reagent reservoirs, and the other nine filled with ultrapure water. The dye-containing reservoir was primed first at 750 mbar until the colour was seen in the effluent. The water-containing reservoirs were primed next at the same pressure for 2 minutes. After ensuring that all the fluidic lines were filled and the dye was washed out of the system, we began the testing. Water was delivered from each reservoir sequentially until (1) colour appeared in the effluent in the case of a leakage pathway, or (2) 3 minutes of remaining clear in the case no leakage pathway exists. If a leakage pathway was present between the dye-containing and tested reservoirs, additional time was given to allow the effluent to run clear before switching to the next. Effluent was collected with a white Kimwipe, aiding in visualization of the colour change. Images were taken of each Kimwipe and the qualitative intensity of colour was recorded in a table. This priming and testing process was repeated with the food dye in each reservoir position, and flowing from each of the nine others to check for leakage pathways. During testing, the connections between the reservoir and the valve were inspected for evidence of leakage as well. In some cases, it was possible to visualize backflow from the dye-containing reservoir into a water reservoir.

#### 2.3 On-chip binding assays with enzymatic amplification

To amplify binding assay signals, HRP-conjugated antibodies or SA-HRP were delivered over the sensors under continuous flow, followed by delivery of a chromogenic substrate solution under flow conditions. The primary substrate employed was 4-chloro-1-naphthol (4CN), which HRP catalyzes to form an insoluble coloured precipitate localized on the sensor surface, enhancing the surface refractive index (RI) change. For 4CN assays, the substrate solutions were prepared as described in Supplementary Information (SI) Section 2.4. Due to the unstable nature of the substrate solution, it was loaded into the microfluidic reservoir immediately after preparation.

On-chip IL-8 immunoassays employing enzymatic amplification utilized the following reagents: IL-8 capture antibody (included in ELISA kit R&D Systems DY208-05, lot P320482); bovine serum albumin (Sigma-Aldrich, A7906-50G, lot SLCL7265) IL-8 (included in ELISA kit R&D Systems DY208-05, lot P320482); HRP-conjugated secondary detection antibody (Sino Biological 10098-MM18-H, lot HO15JA2803), 4-chloro-1-napthol (4CN) solution (Sigma-Aldrich, C8302-100ML, lot SLCD6361) or tablets (Sigma-Aldrich C6788-5TAB) dissolved in methanol and tris-buffered saline. Hydrogen peroxide (Sigma-Aldrich H1009-5ML, lot MKCR9645) was added to 4CN at a final concentration of 0.01% (v/v) immediate prior to use and loaded into the assay fluidic delivery system during the buffer rinse prior to the amplification assay stage. The assay described below employed 4CN solution, while the 4CN used assays described in the main text is specified in respective sections.

An overview of the stages of the IL-8 binding assay with enzymatic amplification described here is presented in Figure S3 below.

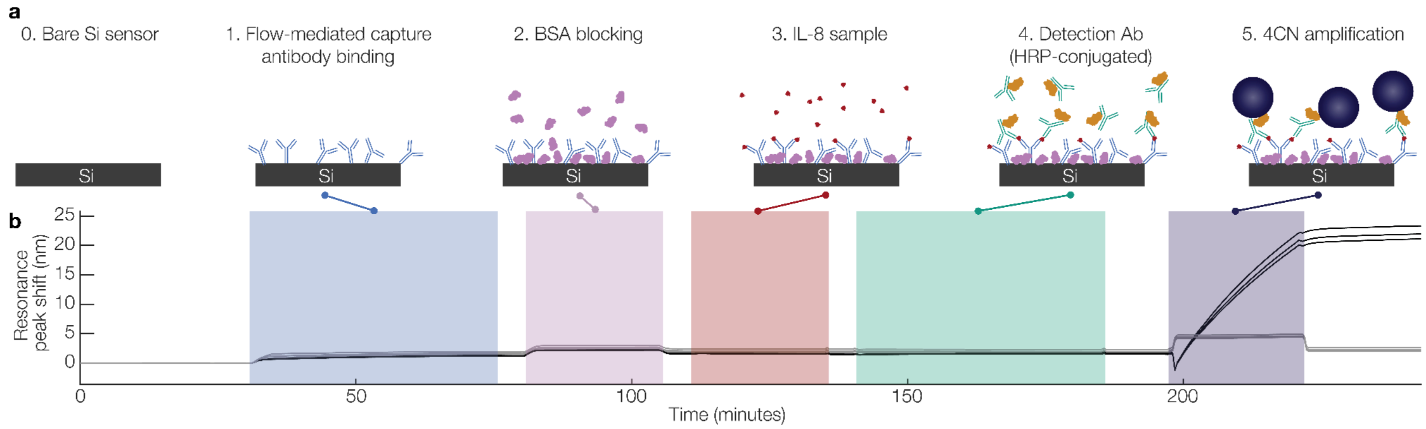


**Figure S3**. Overview of the on-chip sandwich immunoassay developed to detect IL-8 using enzymatic amplification. **(a)** cross-sectional views of the sensor waveguides depicting the assay stages. (0) the bare sensor chip (pre-functionalization) is integrated with microfluidics and PBS is flowed through the channels (30 µL/min). (1) capture antibody solution is supplied to the chip (30 µL/min, 45 mins) to functionalize the surface of the sensors for specific binding. (2) A 10 mg/mL solution of BSA in PBS is delivered to the sensor to block the surface to reduce nonspecific binding (30 µL/min, 25 mins). (3) a sample made up of IL-8 in assay running buffer (either 0 or 3.125 ng/mL) is delivered to the sensor (30 µL/min, 25 mins), and IL-8 specifically attaches to the capture antibody. (4) an HRP-conjugated IL-8 detection antibody (0.5 µg/mL in assay running buffer) is delivered to the sensor surface and binds to the immobilized IL-8. (5) a 4CN substrate solution is delivered to mediate formation of an insoluble enzymatic product (precipitate) that drastically increases detected signal (30 µL/min, 25 mins). Between each assay stage the sensor is rinsed with PBS (before BSA) or assay running buffer (0.1 mg/mL BSA in PBS, 30 µL/min, stages following BSA). **(b)** example sensorgram depicting the stages of the IL-8-detection immunoassay. The sensors in a microfluidic channel to which 3.125 ng/mL IL-8 was delivered during the sample stage are coloured black, while those in a negative control channel (0 ng/mL IL-8) are coloured grey.

In later experiments, a commercial 1-Step 4CN substrate solution (Thermo Scientific 34012, lot YG381483) was tested to reduce assay complexity and minimize variability associated with manual reconstitution and peroxide addition. As a proof-of-concept, a 1-Step precipitate-forming (unlike the typical TMB formulations that produce a soluble product) 3,3′,5,5′-tetramethylbenzidine (TMB)-blotting substrate solution (Thermo Scientific 34018, lot ZA382449) was also tested in a single trial to explore use of an alternate HRP substrate. TMB is supplied ready-to-use and does not require hydrogen peroxide addition, but was not pursued further in this study due to poor amplification signal as described in SI Section 4.

##### 2.4 Dot blots

Dot blot assays were conducted to compare performance of newly prepared substrate solutions made from 4CN tablets with previously used vendor-supplied liquid 4CN stocks. Polyvinylidene fluoride (PVDF) membranes were activated via immersion in a sequence of methanol, water, and TBS baths, then spotted in triplicate with 10 µL volumes of HRP-conjugated secondary antibodies at 1 µg/mL, 0.1 µg/mL, or 0 µg/mL (negative control). After incubation for 1 hour at room temperature, membranes were submerged in 4CN solutions prepared under different conditions: "Old" liquid 4CN , or "New" 4CN prepared from solid tablets, freshly dissolved in methanol and diluted in TBS; both solutions were activated with hydrogen peroxide immediately before use (further details provided in SI Section 2.3. Reactions were allowed to proceed for 5 minutes before quenching with ultrapure water. Membranes were imaged with a gel documentation system, and densitometry analysis was performed to compare product intensities across substrate formulations by quantifying regions of interest in the spotted areas from the membrane images, and performing background subtraction against regions spotted with the vehicle.

##### 2.5 ELISA

To characterize the reagents used in our binding assays in a conventional, less resource intensive format, we incorporated ELISAs. ELISAs were used to validate assay specificity, compare antibody detection strategies, and contextualize on-chip observations. A representative protocol, developed following optimization trials and vendor-provided guidance, is described here. ELISAs were performed on polystyrene 96-well plates to validate an HRP-conjugated IgG antibody as well as compare its performance with a biotinylated antibody supplied as part of a commercial ELISA kit (included in ELISA kit R&D Systems DY208-05, lot P320482). 100 µL capture antibody was dispensed into wells and incubated overnight at room temperature; the following day, plates were washed three times with PBS + 0.05% Tween-20 (PBST) and blocked for 1 hour. In one experiment, alternative blocking agents were employed (reagent details below) and tested against 2% BSA in PBST as shown in Figure S4. Following blocking, recombinant (R&D Systems DY206-5, lot P320482) and native (sourced from Calu-3 cells cultured in microfluidic chips, as described previously) IL-8 antigen samples, along with negative controls, were added and incubated for 2 hours at room temperature [7], [13]. After washing, secondary antibody solutions were added; when employing biotinylated detection antibody, a 10 ng/mL solution was dispensed and incubated for two hours, followed by washes and subsequent addition of a 40-fold dilution of SA-HRP which was incubated for 20 minutes. When employing the HRP-conjugated secondary antibody for IL-8, depending on the particular assay, different concentrations were assessed (ranging from 10-1000 ng/mL, specified where results are presented) and also incubated for 2 hours. All HRP-containing reagents were protected from light. Plates were then washed and incubated with TMB ELISA Substrate (R&D Systems DY 999B lot P348181) for 15 minutes prior to quenching with stop solution (R&D Systems DY 994 lot MKCJ1597). Absorbance was read at 450 nm with a 540 nm reference using a plate reader (Molecular Devices, San Jose).

#### 3. Orthogonal Assays to Characterize Reagents and Support Decision-Making

Results from initial characterization and concentration-titration of HRP-conjugated antibody alongside recombinant IL-8 and anti-IL-8 capture antibody from a commercial ELISA kit are included in Figure 4(b) of the main text. Included below in Figure S4 are results from an experiment where HRP-conjugated detection antibody was employed in parallel to the biotinylated detection antibody included as part of a commercial ELISA kit; the objective of this experiment was to compare performance between the antibodies with respect to background signal and sensitivity, while also assessing the ability of different commercial blocking agents to mitigate background signal. The blocking agents tested included StartingBlock (Thermofisher 37578, lot XH353524) and BlockAid (Thermofisher B10710, lot 2406544). As illustrated in Figure S4, substantial background signal presents when the HRP-conjugated detection antibody is used with samples in a vehicle of complete culture medium, a phenomenon not seen with biotinylated antibody. The biotinylated antibody, furthermore, exhibits more robust signal when detecting both recombinant and native antigen when compared to the HRP-conjugated alternative. Neither of the commercial blocking agents assessed provided significant benefit over BSA in PBST in their ability to mitigate non-specific background while preserving positive control signal magnitude.

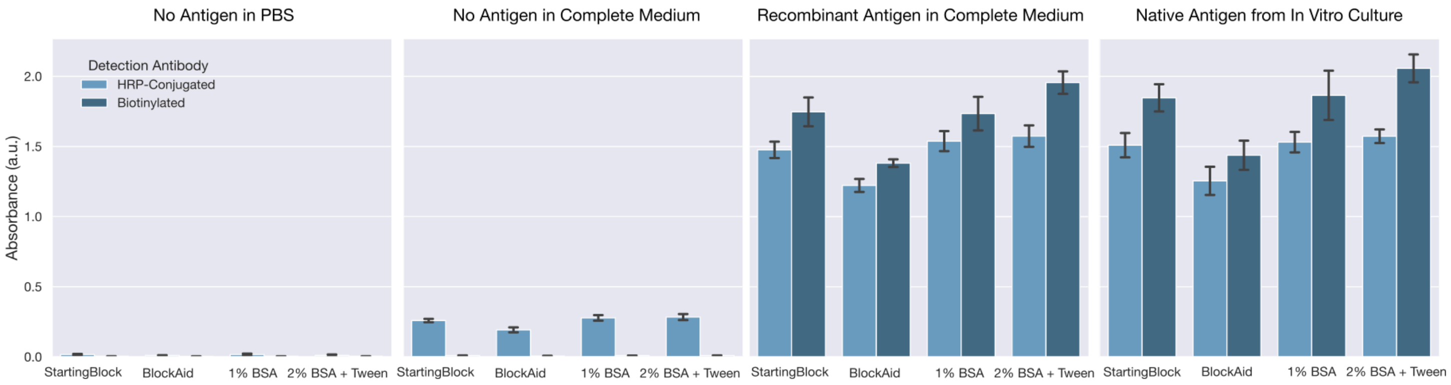


**Figure S4.** Detection antibody and blocking agent comparison across assay matrices. These bar plots depict the signal generated by a 10 ng/mL concentration of a biotinylated detection antibody alongside an HRP-conjugated detection antibody across vehicles of PBS and complete culture medium, as well as complete culture medium spiked with recombinant antigen and native antigen sourced from Calu-3-laden in vitro cell cultures. Both recombinant and native antigen were present in solution at 500 pg/mL; in-vitro samples with ELISA-measured IL-8 concentrations were pooled to achieve a 500 pg/mL working concentration, and recombinant IL-8 was spiked into complete culture medium at 500 pg/mL. Assays were performed with three technical replicates per condition and error bars represent one standard deviation across those replicates; bar height represents mean absorbance measured by a SpectraMax plate reader at 450 nm with reference subtraction at 540 nm. For the HRP-conjugated antibody, BlockAid reduced background signal in culture medium but also attenuated true signal. By contrast, background signal with the biotinylated antibody was negligible under all blocking conditions.

These results led us to ultimately forgo use of the HRP-conjugated detection antibody in favor of the biotinylated detection antibody, despite it requiring an additional streptavidin-HRP step, due to the markedly improved performance it exhibited in-well. These results, furthermore, enabled us to decide that use of a more costly and proprietary commercial blocking formulation was not worthwhile at that time.

#### 4. Overview of extended fluidic control system cleaning procedure

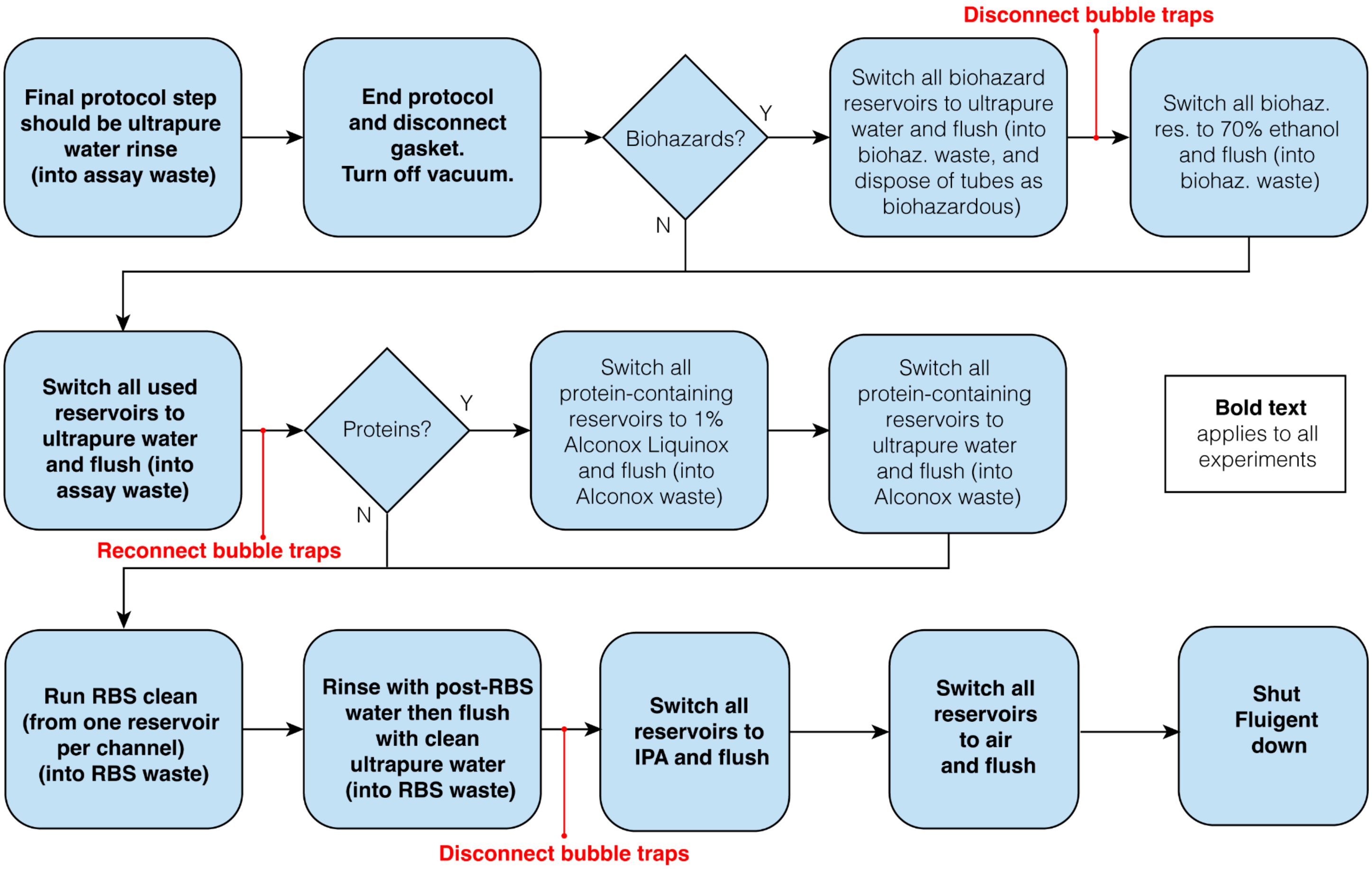


**Figure S5**. Overview of the extended cleaning protocol for all experiments using the fluidic control system. Flowchart outlines steps depending on the contents of the assay and whether biohazardous and/or protein-containing reagents were used. Biohazardous lines need to be flushed with 70% Ethanol for decontamination, and protein-containing lines need to be cleaned with 1% Alconox Liquinox. Bubble traps are disconnected and reconnected during alcohol-delivery steps to prevent these organic solvent solutions from wetting the bubble trap membrane (which renders it ineffective and contributes to other assay issues like reagents entering vacuum lines instead of being delivered to the microfluidic device). Steps written in bold text applied to all experiments.

#### 5. Supporting Data for Fishbone Analysis and Inter-assay Variability Mitigation

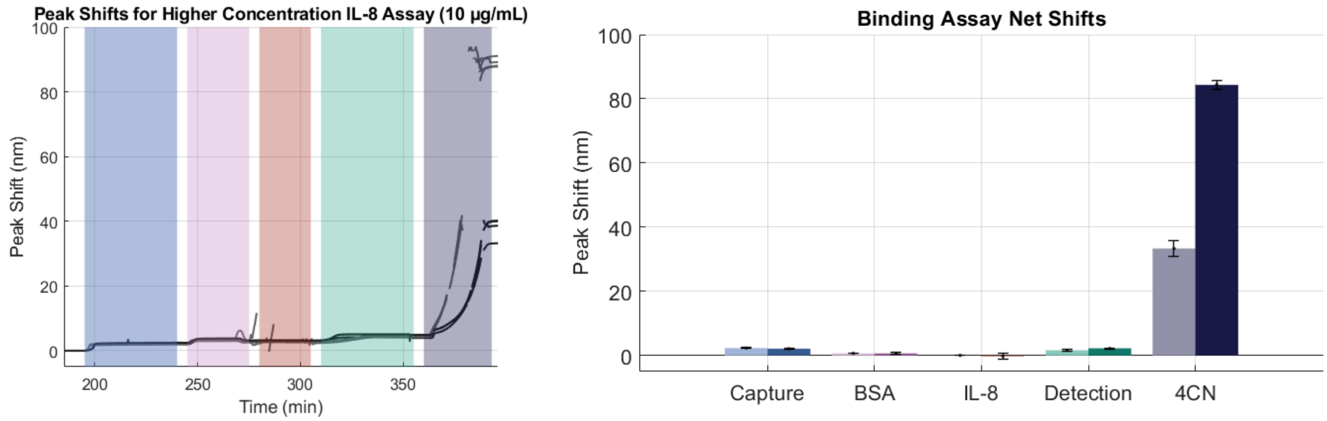


**Figure S6.** Peak shifts for higher concentration IL-8 binding assay (10 µg/mL). (Left) Time-resolved peak shift traces from two channel-replicates (gray and black) are shown, with assay stages overlaid: capture antibody (blue), BSA blocking (purple), IL-8 antigen (red), detection antibody (teal), and enzymatic amplification with 4CN (indigo). (Right) The capture antibody shift averaged 2.32 ± 0.19 nm (ch1) and 2.12 ± 0.14 nm (ch2), the BSA blocking step shift averaged 0.67 ± 0.04 nm (ch1) and 0.67 ± 0.29 nm (ch2), the IL-8 antigen shift was 0.013 ± 0.002 nm (ch1) and –0.20 ± 1.00 nm (ch2), the detection antibody produced clear shifts of 1.62 ± 0.16 nm (ch1) and 2.22 ± 0.15 nm (ch2), while enzymatic amplification yielded large responses (33.4 ± 2.46 nm, ch1; 84.3 ± 1.45 nm, ch2). Net shifts associated with IL-8 binding, extracted from only those peaks exhibiting stability and continuity, averaged below 20 pm in both channels (negative in the top channel), indicating that the IL-8 concentration employed was insufficient to clearly visualize direct binding. By contrast, direct binding of the HRP-conjugated detection antibody was observable, averaging >1.5 nm in both channels. The ratio of shifts between detection antibody binding and IL-8 binding greatly exceeded expectations from the ratio of their molecular weights; for example, a 13 pm IL-8 shift in the bottom channel would predict ~220 pm of detection antibody signal assuming 1:1 binding of detection antibody and IL-8 (13 pm × 150 kDa/8.9 kDa), but 1620 pm was observed. This discrepancy may reflect non-specific binding of detection antibody, which was deployed at 20-fold higher concentration than in previous assays. Although the enzymatic amplification produced a large signal magnitude, the associated shift rate and sweep range exceeded reliable tracking limits.

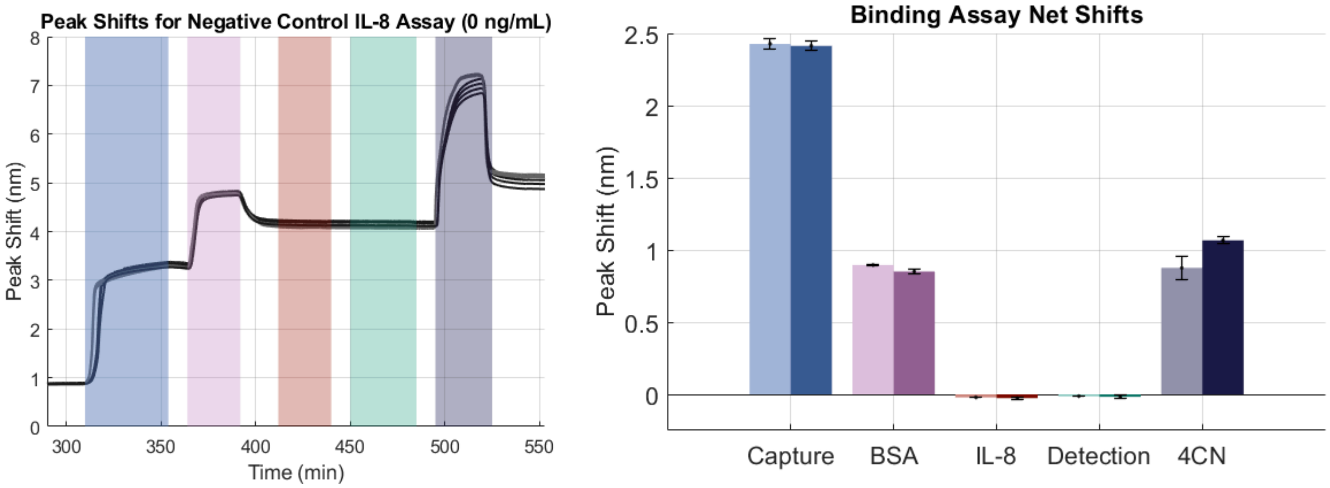


**Figure S7.** Peak shifts for a negative control IL-8 binding assay (0 ng/mL). (Left) Time-resolved peak shift traces from two channel replicates (gray and black) are shown, with assay stages overlaid: capture antibody (blue), BSA blocking (purple), IL-8 antigen (red), detection antibody (teal), and enzymatic amplification with 4CN (indigo). (Right) The capture antibody shift averaged 2.43 ± 0.04 nm (ch1) and 2.41 ± 0.03 nm (ch2), the BSA blocking step shift averaged 0.90 ± 0.01 nm (ch1) and 0.85 ± 0.01 nm (ch2), the IL-8 antigen shift was −0.021 ± 0.001 nm (ch1) and −0.026 ± 0.009 nm (ch2), and the detection antibody shift was −0.013 ± 0.001 nm (ch1) and −0.017 ± 0.013 nm (ch2). These negative IL-8 and detection antibody shifts reflect values beneath the quantifiable limit, consistent with the absence of antigen in this assay. Enzymatic amplification with 4CN produced modest shifts of 0.88 ± 0.08 nm (ch1) and 1.07 ± 0.03 nm (ch2). We would expect negligible amplification shifts if no detection antibody bound to the chip in the absence of IL-8. The observed signal indicates that some non-specific binding did occur.

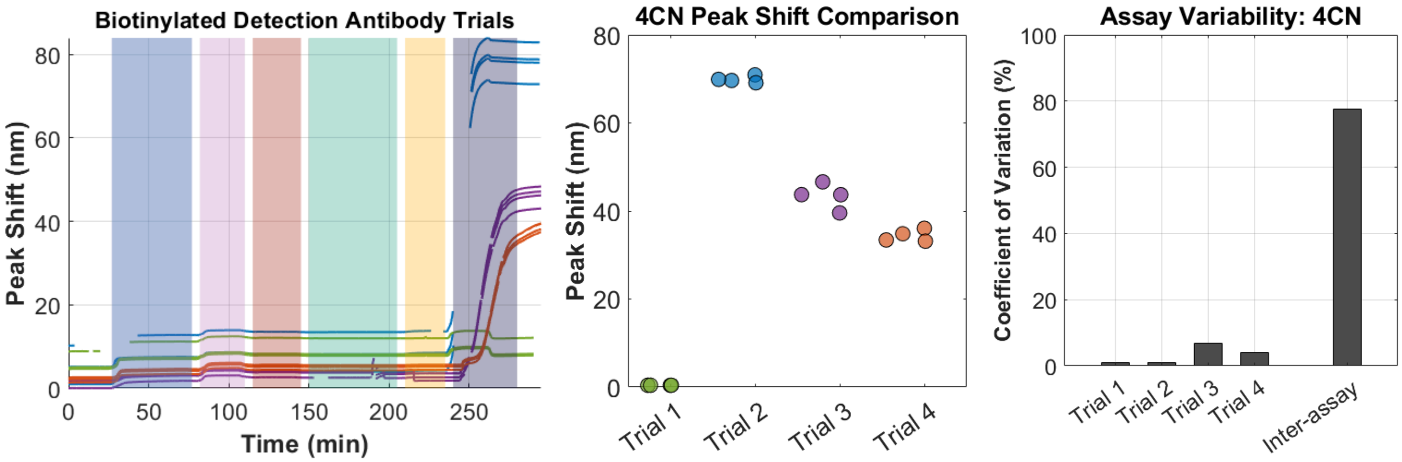


**Figure S8.** Inter-assay variability in biotinylated detection antibody trials with SA-HRP and 4CN amplification.
(Left) Time-resolved peak shift traces for four channel replicates are shown (green, blue, purple, orange), with assay stages overlaid: capture antibody (blue), BSA blocking (purple), IL-8 antigen (red), detection antibody (teal), SA-HRP (yellow), and enzymatic amplification with 4CN (indigo). (Middle) Beeswarm plot of 4CN peak shifts for each sensor (four sensors per channel) across the four channel replicates, highlighting the spread in individual sensor responses. (Right) Bar plot showing intra-assay variability for each trial and the overall inter-assay CV (%) across the four trials (77%), illustrating that substantial inter-assay variability persists despite switching from the HRP-conjugated antibody to the more compatible biotinylated detection antibody and SA-HRP. This variability may reflect factors such as limited sample size (n = 4), variability in flow-mediated sensor functionalization that relied on passive adsorption of antibody, sensor-to-sensor differences, or persistent non-specific binding effects.

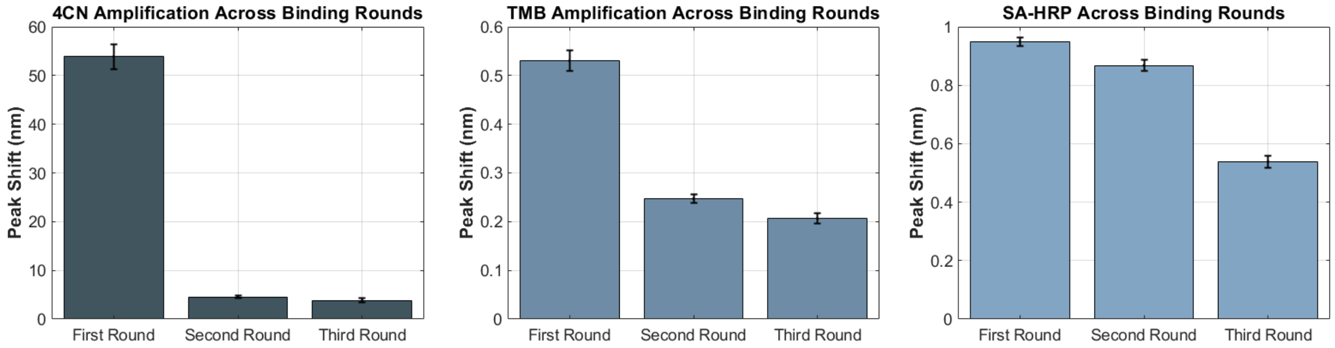


**Figure S9.** Cycle-to-cycle binding signal highlighting the regeneration performance across assay formats. This figure shows the binding signal for on-chip IL-8 assays comprising three binding cycles with attempted sensor regeneration using low-pH glycine–HCl regeneration after each binding cycle. Three assay formats are compared: 1-step 4CN, 1-step precipitate-forming (blotting) TMB, and SA-HRP without enzymatic amplification. Bars represent signals measured after the first, second, and third regeneration cycles, illustrating retention of functional and accessible capture antibody and assay performance. For 4CN, we observed 91.6% signal loss between the first and second round and 15.9% between the second and third, corresponding to a 92.9% overall signal loss by the third round. For TMB, we observed 53.4% signal loss between the first and second round and 16.6% between the second and third, yielding a 61.1% overall signal loss. Because low-pH glycine–HCl provided little regeneration for TMB, an additional high-pH step using 50mM NaOH (pH ≈ 13) in 1M NaCl was included [14], but performance remained poor. Exploration of the TMB substrate was ultimately abandoned due to low amplification shifts (~0.5 nm) and the fact that SA-HRP alone produced similar amplification. For SA-HRP without enzymatic amplification, regeneration was comparatively higher, with 8.6% signal loss between the first and second round and 37.9% between the second and third, corresponding to 43.2% overall signal loss. One experimental replicate is shown for each assay format, with error bars depicting one standard deviation of the signals from n = 4 sensors. We hypothesize that the tested regeneration solutions were insufficient to remove the precipitate formed during enzymatic amplification despite their ability to disrupt the antibody-antigen binding interactions when enzymatic amplification was not used. Although additional regeneration optimization would be beneficial, the cycle-to-cycle signal decrease in the assay that did not employ enzymatic amplification (right) was reasonable.

### References

[1] “Experimental Cycle [and other diagrams] | i-Biology,” 2014. https://i-biology.net/myp/experimental-cycle/ (accessed Sep. 11, 2025).

[2] International Baccalaureate Organization, “Middle Years Programme Sciences Guide,” International Baccalaureate Organization (UK) Ltd, 2019.

[3] J. T. Kindt, M. S. Luchansky, A. J. Qavi, S.-H. Lee, and R. C. Bailey, “Subpicogram per milliliter detection of interleukins using silicon photonic microring resonators and an enzymatic signal enhancement strategy.,” *Anal. Chem.*, vol. 85, no. 22, pp. 10653–10657, Nov. 2013, doi: 10.1021/ac402972d.

[4] L. S. Puumala *et al.*, “Resonating with replicability: factors shaping assay yield and variability in microfluidics-integrated silicon photonic biosensors,” *BioRxiv*, Jul. 2025, doi: 10.1101/2025.07.16.664198.

[5] L. S. Puumala *et al.*, “An Optimization Framework for Silicon Photonic Evanescent-Field Biosensors Using Sub-Wavelength Gratings.,” *Biosensors (Basel)*, vol. 12, no. 10, Oct. 2022, doi: 10.3390/bios12100840.

[6] “NanoSOI Design Center | Applied Nanotools Inc.” https://www.appliednt.com/nanosoi/sys/resources/specs/ (accessed Aug. 27, 2022).

[7] T. C. Cameron *et al.*, “PDMS Organ-On-Chip Design and Fabrication: Strategies for Improving Fluidic Integration and Chip Robustness of Rapidly Prototyped Microfluidic In Vitro Models.,” *Micromachines (Basel)*, vol. 13, no. 10, Sep. 2022, doi: 10.3390/mi13101573.

[8] J.-Y. Byeon, F. T. Limpoco, and R. C. Bailey, “Efficient bioconjugation of protein capture agents to biosensor surfaces using aniline-catalyzed hydrazone ligation.,” *Langmuir*, vol. 26, no. 19, pp. 15430–15435, Oct. 2010, doi: 10.1021/la1021824.

[9] M. S. Luchansky and R. C. Bailey, “Silicon photonic microring resonators for quantitative cytokine detection and T-cell secretion analysis.,” *Anal. Chem.*, vol. 82, no. 5, pp. 1975–1981, Mar. 2010, doi: 10.1021/ac902725q.

[10] A. L. Washburn, L. C. Gunn, and R. C. Bailey, “Label-free quantitation of a cancer biomarker in complex media using

silicon photonic microring resonators.," *Anal. Chem.*, vol. 81, no. 22, pp. 9499–9506, Nov. 2009, doi: 10.1021/ac902006p.

[11] A. L. Washburn, M. S. Luchansky, A. L. Bowman, and R. C. Bailey, "Quantitative, label-free detection of five protein biomarkers using multiplexed arrays of silicon photonic microring resonators.," *Anal. Chem.*, vol. 82, no. 1, pp. 69–72, Jan. 2010, doi: 10.1021/ac902451b.

[12] Fluigent, "Microfluidic Bidirectional Valve - Fluigent," 2021. https://www.fluigent.com/research/instruments/microfluidic-valves/m-switch/ (accessed Sep. 10, 2025).

[13] A. Randhawa, "Exploring the microfluidic organ-on-chip platform for aerosol exposure study," *University of British Columbia*, 2022, doi: 10.14288/1.0421355.

[14] C. Cao and S. J. Sim, "Signal enhancement of surface plasmon resonance immunoassay using enzyme precipitation-functionalized gold nanoparticles: a femto molar level measurement of anti-glutamic acid decarboxylase antibody.," *Biosens. Bioelectron.*, vol. 22, no. 9–10, pp. 1874–1880, Apr. 2007, doi: 10.1016/j.bios.2006.07.021.
