## Supplemental protocol for IL-8 assay for "What Could Go Wrong? Promoting Success by Planning for Failure in Label-Free Biosensor Assay Development"

IL-8 biosensing assay trial X

### GOALS

The objective of this experiment is to troubleshoot ongoing issues with low IL-8 signals. The experiment is designed with two channels. Both channels will use BSA blocking in PBS. The assay will involve exposing the sensor to a 0 ng/mL IL-8 negative control solution (C1), followed by five exposures to a 3.125 ng/mL IL-8 sample (C5).

- Top fluidic channel (Fluigent #1)
  - C1 (0 ng/ml IL-8) → C5→ C5→ C5 → C5 → C5(3.125 ng/ml IL-8)
- Bottom fluidic channel (Fluigent #2),
  - C1 (0 ng/ml IL-8) → C5→ C5→ C5 → C5 → C5(3.125 ng/ml IL-8)

We will deliver exactly the same concentration in the same order into both channels, in an effort to test whether the inter-channel variability has been improved by the installation of the new Fluigent M-switch on Jan. 13, 2025 due to leakage that was observed with the previous M-switch. We will use sensor chips with our multiplexed SiP sensor architecture for this assay.

Questions we hope to answer through this experiment are listed below:

1. Will fluidic channel #2 perform the same as #1 after replacing the M-switch (eliminating the leakage between reservoirs), and with fresh reagents?
2. Do we see replicable detection shifts across 5 binding cycles of the same concentration?

### SUMMARY OF PREVIOUS RESULTS

1. Results from previous trial:
   a. Significantly lower binding shifts (~85-330% lower) in channel 2 than in channel 1
   b. 20% residual signal in binding round with 0 ng/mL IL-8, indicating a significant amount of non-specific binding
   c. Possible wetting of bubble trap membrane in channel 1. Liquid was being drawn into the vacuum line, suggesting that the actual flow rate was lower than expected for the first binding round until the vacuum was turned off.
2. Results from leakage testing:
   a. Identified multiple variable leakage pathways in Fluigent M-switch #2
   b. After replacing M-switch, another round of leakage testing was conducted and no leakage pathways were identified.

### HYPOTHESES AND EXPECTATIONS

1. Fluigent channel #2 will yield the same binding shifts compared to Fluigent channel #1.
2. Using the new detection antibody and Strep-HRP, as well as a Fluigent system cleaned with Alconox, will help to enhance the binding shifts.
3. Combining cell culture medium with BSA for blocking will not cause a significant change of Strep-HRP binding.
4. Including a reference signal and sensor will allow us to remove unwanted sensorgram effects.
5. Well-aligned channels will reduce noise in sensogram signals.

### CONTROLS/EXPERIMENTAL GROUPS

We will use 2 microfluidic channels and 6 detection cycles. Both fluidic channels will serve as experimental replicates and will, therefore, be exposed to identical samples throughout the experiment. There will be one negative control sample detection cycle in both channels at the beginning of the assay, followed by five IL-8 sample detection cycles. Analyte detection will be performed in complete cell culture medium (CCM)

We will use 2 functionalisation conditions:

- IL-8 capture antibody spots on upstream and downstream sensor groups (240 µg/mL IL-8 cAb in spotting buffer)
- BSA control spot on middle sensor group (20 mg/mL BSA in spotting buffer)

Each channel contains 6 functionalized sensors, split between the three functionalisation spots.

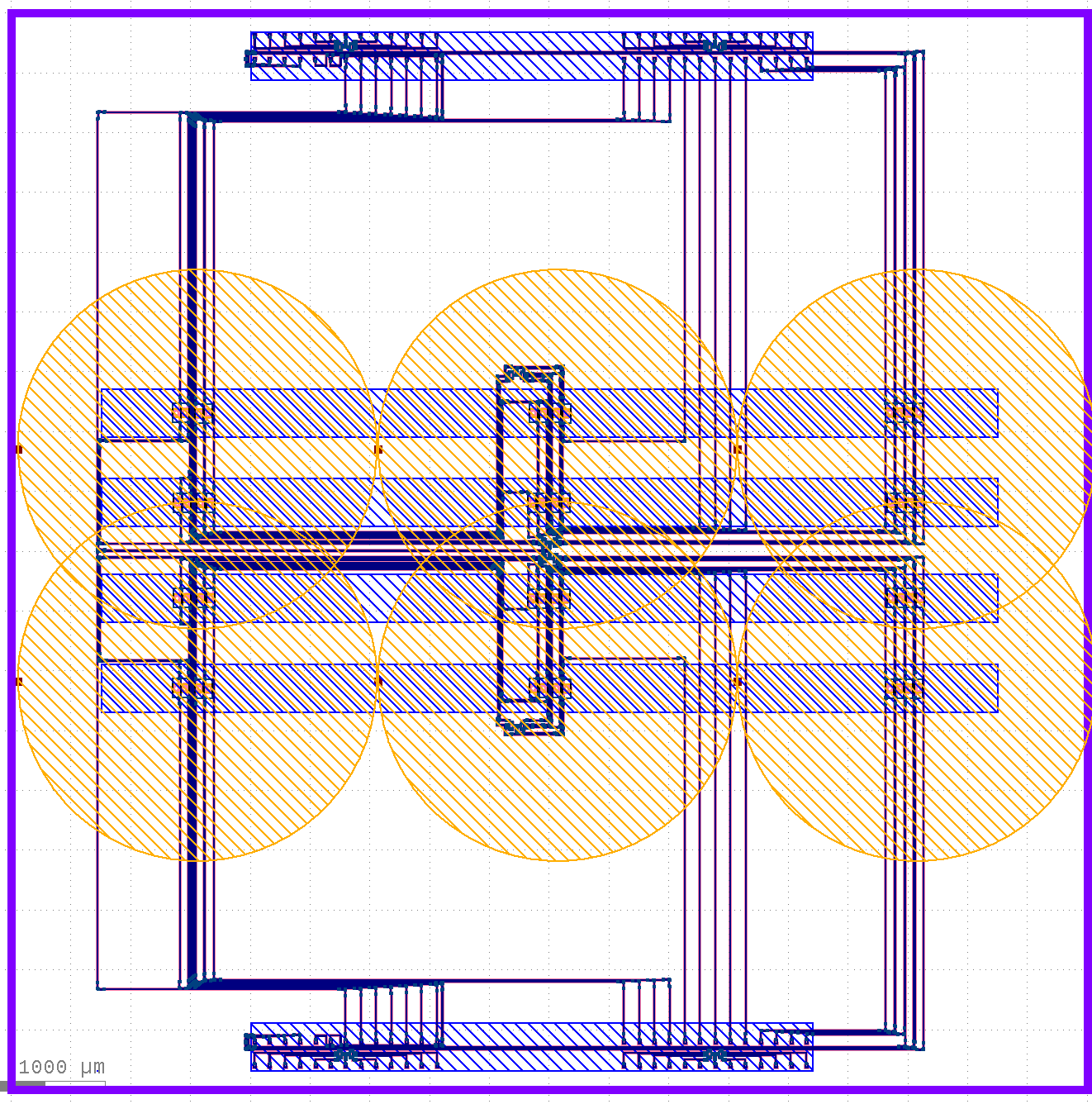

µFluidic 1b

µFluidic 1a

µFluidic 2b

µFluidic 2a

**IL-8 cAb droplet**

**Ref Droplet**

**IL-8 cAb droplet**

Overview of assay conditions used for both fluidic channels:

- Bubble mitigation conditions
  - Gasket treatment: overnight gasket degassing + plasma treatment
  - Pre-wetting fluid: 0.3 mM Triton X-100 in PBS
  - Bubble traps? Yes
- Blocking: 20 mg/mL BSA in PBS
- Running buffer: 0.1 mg/mL BSA in PBS (PBS-BSA)
- Sample:
  - Cycle 1: negative control C1 (0 ng/ml IL-8 in CCM)
  - Cycles 2–6: sample C5 (3.125 ng/ml IL-8 in CCM)
- Amplification: 1µg/mL IL-8 detection antibody + 2 µg/mL SA-HRP in running buffer
- Bulk RI: PBS-dilution-based bulk refractive index after assay

### HYPOTHESIS-TESTING STRATEGY

To test all of our defined hypotheses, an IL-8 binding assay will be performed on the functionalized sensors. We will design the assay and perform quantification as described below in order to test each outlined hypothesis.

1. **Hypothesis 1:** Fluigent channel #2 will yield the same binding shifts compared to Fluigent channel #1.
   a. **Hypothesis-testing strategy:** Signals obtained in channels 1 and 2 will be compared by running a T-test. Data from replicate trials will be collected and analyzed to assess signal reproducibility, variability and consistency by calculating the coefficient of variation (CV).
2. **Hypothesis 2:** Using the new detection antibody and Strep-HRP, as well as a Fluigent system cleaned with Alconox, will help to enhance the binding shifts.
   a. **Hypothesis-testing strategy:** The signals for the C5 sample cycles obtained in this assay will be compared to previous assays involving the detection of the same analyte concentration. Signal reproducibility, variability and consistency will be analyzed by calculating the coefficient of variation (CV) across different assays.
3. **Hypothesis 3:** Combining cell culture medium with BSA for blocking will not cause a significant change of Strep-HRP binding.
   a. **Hypothesis-testing strategy:** The chip will be incubated in BSA and PBS for blocking after surface functionalization, and the binding shift results from the first round (zero concentration) will be compared to the first round in channel 1 of Trial IIIc.
4. **Hypothesis 4:** Including a reference signal and sensor will allow us to remove unwanted sensorgram effects.
   a. **Hypothesis-testing strategy:** The upstream and downstream droplets will contain capture antibody. The middle will act as a reference/control droplet containing BSA instead.
5. **Hypothesis 5:** Well-aligned channels will reduce noise in sensogram signals.
   a. **Hypothesis-testing strategy:** Images will be taken before and after alignment, checking that the rings are entirely aligned with each channel.

### REQUIRED REAGENTS AND EQUIPMENT

#### Required supplies

##### Setup:

- LSI Maple Leaf Stage
- Fluigent System

##### Instrumentation:

- Aven Tools microscope
- N77 detector
- C-band Laser
- 12-channel hybrid fibre array
- Abbe refractometer

##### Tools:

- Soft-tip tweezers
- 4-40 Screws (for mounting the chip and gasket to the plate)
- Acrylic washers
- Mounting plate

##### Consumables and equipment for PDA coating and offline functionalization:

- Scoopula
- Pipettes
- Weigh paper
- Analytical scale
- 1 × 100 mL glass beaker
- Serological pipette (50 mL) and pipette aid
- 1 × magnetic stir bars
- Magnetic stir plate
- Stir bar retriever
- 1 × Small (~60 mm diameter) glass crystallisation dish
- Transfer pipettes
- Waste beaker
- Waste bottle
- Cleanroom wipe
- Soft tip tweezers
- 1 × 12-well plate (Thermo Nunclon Delta 150628)

##### Consumables for Assay:

- ANT Multiplexed Sensing Chip
- ANT Multiplexed Sensing Gasket
- Tygon and PEEK tubing
- Petri dish
- Falcon and Eppendorf tubes for reagent preparation
- 1 × 250 mL glass bottle
- 1 × 20 mL glass scintillation vial
- 40 µm cell strainers
- Pipettes

##### Reagents:

| Reagent | Vendor | Product number | Lot number | Stock concentration | Aliquot info | Notes |
| --- | --- | --- | --- | --- | --- | --- |
| Dopamine hydrochloride | Sigma | H8502-5G | BCCJ9540 | Powder | – |  |
| Tris buffer, 0.5 M, pH 8.6 | Thermo Scientific | AAJ62287AP | P24K523 | 1× | – |  |
| Nitrogen Gas, 99.998% purity | Linde | NI 4.8-T | – | – | – |  |
| Ultrapure water, ASTM Type I (ddW) | Nanopure Diamond system | – | – | – | – |  |
| Phosphate buffered saline (PBS), pH 7.4, 1X | Gibco | 10010-023 | 2842947 | 1× | – |  |
| Glycerol | Fisher Scientific | G33-4 | 116960 | 100% | – |  |
| Triton X-100 | Sigma Aldrich | T8787-100ML | SLCJ6163 | 100% | – |  |
| BSA | Sigma | A706-50G | Lot: 0000282927  Source: 0000282927 | 200 mg/mL in 1× PBS | Aliquoted: 2024-08-09  Vol: 600 µL |  |
| IL-8 Capture Antibody | R&D Systems | DY208-05 | ASJ3824031 | 480 ug/mL | Reconstituted 2025-02-19  useable within 8 weeks (expire 2025-04-16) | IL-8 ELISA Kit, picked up 2025-02-03, Lot No. P437281. Part No. 890804 |
| IL-8 Standard | R&D Systems | DY208-05 | P390129 | 120 ng/mL | Reconstituted 2025-02-19  useable within 4 weeks (expire 2025-03-19) | IL-8 ELISA Kit, picked up 2025-02-03, Lot No. P437281, Part No. 890806 |
| IL-8 Detection | R&D Systems | BAF208 | UM2322052 | 200 ug/mL | Aliquoted 2025-02-19 useable within 6 months | 50 ug -> 250 uL  12 months from date of receipt, -20 to -70 °C as supplied.  1 month, 2 to 8 °C under sterile conditions after reconstitution.  6 months, -20 to -70 °C under sterile conditions after reconstitution.  Reconstitute at 0.2 mg/mL in sterile PBS.  62.5uL per aliquot for a total of 4 aliquots! |
| Streptavidin-HRP | 21130 | 21130 - Pierce™ High Sensitivity Streptavidin-HRP - 1.00 | ZF394944 | 1.0-1.1 mg/mL | – | Pierce, High-Sensitivity. Picked up 2024-11-26, LOT: ZF394944.  useable within 12 months |
| RBS 35 Detergent | Thermo Scientific | 27950 | WB3191271 | 100% | – |  |
| DMEM | Sigma Aldrich | D6421-500ML | RNBM3386 | – | – |  |
| FBS | Gibco | 12483020 | 2357664RP | – | 5 mL aliquots |  |
| L-glutamine | Sigma Aldrich | G7513-100ML | RNBM3694 | 200 mM | 1 mL and 10 mL aliquots |  |
| Antibiotic-  Antimycotic | Thermo Scientific | 15240-062 | 2585910 | 100X | 1 mL and 10 mL aliquots |  |

### SAFETY

- Appropriate PPE must be worn when running this protocol: **gloves, a lab coat, safety goggles, loose-fitting long pants (no ankle visible), and closed-toed shoes**.
  - Do not touch any designated gloved surfaces without gloves.
  - Do not touch your face, clothes, or any personal items (e.g., phone, mask) with your gloves.
  - Do not place your phone or any other personal items on any gloved surfaces.
- Chemical safety: refer to relevant SDS documents
  - Hydrochloric acid
    - HCl is highly corrosive. It can cause serious chemical burns to skin, as well as serious eye damage if not handled appropriately.
    - Always wear the following PPE when handling HCl solutions: lab goggles, lab coat, nitrile gloves, long pants, closed-toe splash-resistant shoes.
    - Handle concentrated solutions in a fume hood.
    - Make sure as little skin as possible is exposed when handling HCl, especially concentrated solutions.
      - If your lab coat sleeves tend to ride up, consider trucking them into your gloves or taping them to your gloves while handling HCl.
  - Triton X-100 solution
    - The product is chemically stable under standard ambient conditions (room temperature).
    - The solution will form an explosive mixture with air on intense heating.
    - Violent reactions are possible with:
      - Strong oxidizing agents
      - Strong acids
    - Strong heating is to be avoided.
    - Wash skin thoroughly after handling.
    - Do not eat, drink or smoke when using this product.
    - Avoid release to the environment.
    - IF SWALLOWED: Call a POISON CENTER/ doctor if you feel unwell. Rinse mouth.
    - IF ON SKIN: Wash with plenty of water.
    - IF IN EYES: Rinse cautiously with water for several minutes. Remove contact lenses, if present and easy to do. Continue rinsing. Immediately call a POISON CENTER/ doctor.
    - If skin irritation occurs: Get medical advice/ attention.
    - Take off contaminated clothing and wash it before reuse.
    - Collect spillage.
    - Dispose of contents/ container to an approved waste disposal plant.
  - RBS-35 Concentrate solution
    - Category 2: Skin corrosion & eye irritation
    - SKIN CONTACT: Wash off immediately with plenty of water for at least 15 minutes. Remove and wash contaminated clothing and gloves, including the inside, before re-use. Immediate medical attention is required.
    - EYE CONTACT: Rinse immediately with plenty of water, also under the eyelids, for at least 15 minutes. Immediate medical attention is required.
    - INGESTION: Never give anything by mouth to an unconscious person. Do not induce vomiting without medical advice. Get medical attention if symptoms occur.
    - Inhalation: Remove to fresh air. If not breathing, give artificial respiration. If symptoms persist, call a physician.
    - Collect waste and dispose of as hazardous waste.
  - Cell culture medium: DMEM, FBS, L-glytamine, and anti-anti
    - Individual components are non-hazardous, but cell culture media is a potential biohazard
    - SKIN CONTACT: Wash off immediately with plenty of water for at least 15 minutes. Remove and wash contaminated clothing and gloves, including the inside, before re-use.
    - EYE CONTACT: Rinse immediately with plenty of water, also under the eyelids, for at least 15 minutes. Remove contact lenses, if present and easy to do. Continue rinsing.
    - INHALATION: Remove to fresh air. If not breathing, give artificial respiration. If symptoms persist, call a physician.
    - Collect waste in a separate container and dispose of using bleach.
      - Add bleach to a final concentration > 10% in the container and then pour into the bleached waste container.
      - Neutralize all bleach solutions with very high pH (11-14) prior to sink disposal.
        - Bleach can be quite damaging to plumbing if used in excess or if inappropriately disposed of.
      - Use safe and practical bleach neutralizers: sodium metabisulphite (Na_2_S_2_O_5_), sodium bisulphite (NaHSO_3_), sodium sulphite (Na_2_SO_3_), sodium thiosulphate (Na_2_S_2_O_3_), 3% hydrogen peroxide (H_2_O_2_).
        - The use of ¼ to 1 teaspoon of solid neutralizer is typically sufficient to neutralize 1-4L of volume of 10% bleach solution.

### RISK MITIGATION

- Damage to SiP chip during PDA deposition
  - Ensure that the crystallisation dish with the dopamine solution is placed on the stir plate before adding the SiP chip and stir bar
  - Add the stir bar first. Move the dish such that the magnet holds the stir bar in place somewhat off-centre in the dish without risk of colliding with the dish walls
  - Add the SiP chip. Position it so it is as far away from the stir bar as possible.
  - When starting to stir the solution, begin at a very low speed and gradually increase to a low but steady stir speed.
- Capture antibody spots run into each other during functionalization
  - Spot carefully with 1 uL at a time, do not eject to the second stop. Wait ~1 minute between depositing each droplet to assess antibody spreading
- Ensure the gasket screws are adequately tightened to the “sweet spot” to prevent leakage but avoid over tightening and causing channel distortion.
- Carefully examine the Fluigent protocols and reservoir set-up to ensure that the fluids are being injected into the gasket at the expected and desired times.
- Ensure that the Fluigent lines are properly primed with the correct solutions before the Bulk RI and assay protocols are run.
- Reservoir contamination
  - Prepare new reagents before every replicate test.
  - Flush all protein-containing lines with 2% Alconox solution after each experiment.
- Accidental exposure of the functionalized surface to reagents that could damage it:
  - Prime the Fluigent system prior to connecting the gasket.
  - Examine the reservoir setup and predict which reservoirs will be traversed during the experiment (since the M-switch takes the shortest path by default). Adjust reservoir setup and M-switch direction in the Fluigent protocol if needed.
- Impurities and Fluigent blockages
  - Make sure that reservoir tubes do not touch anything (including your gloves) when changing reservoirs.
  - If the outside of the reservoir tubes need to be cleaned, use a wash bottle or new Falcon tube to rinse the outside of the tubing with clean ddW and wipe if needed using a lint-free cleanroom wipe only (not Kimwipe or paper towel).
  - Rinse tubing and gasket with >1 mL water post-experiment, then push air through the system.
  - Clean the system and tubing with RBS after each experiment
    - Ensure an RBS clean run has been run before commencing
  - Check Fluigent pressures/flow rates before use and during and after priming to make sure that blockages are not likely to impact the assay
  - Detach tubing from set-up (carefully preventing tubing tips from contacting surfaces) and store in clean petri-dishes
  - If blockages occur, first try backflow into a waste reservoir (double-check that it is empty when you start the backflow!), then try to isolate the problem to a length of tubing and switch it out if needed. Follow our blockage troubleshooting instructions (several slides) to solve the issue.
- Inconsistent freeze-thaw cycles impacting replicability
  - Prepare and freeze IL-8 aliquots 1-2 days prior to assay
- Inaccurate flow rate measurements
  - Perform the RBS clean after each experiment, ensure RBS clean has been performed prior to assay
  - Understand that flow rate sensor accuracy is low at flow rates below 10 μL/min
- Running out of reagents
  - Double-check the flow rates in your Fluigent protocol and calculate the required amount of fluid, then double-check the amount of fluid remaining in each reservoir just before starting the protocol
  - Include a buffer amount of extra fluid in each reservoir in case of slight inaccuracies in flow rate measurements
- Inconsistent reagent concentrations
  - Ensure correct volumes of reagents and buffer solutions are used.
  - Ensure reagents are properly and adequately mixed (triturate at least 10× with a pipette volume of at least ⅓ the prepared volume of the reagent.
- Bubble nucleation within fluidic lines and connectors
  - Minimise microfluidic connections in pre-wetting set-up
  - Ensure PEEK tubing connecting to each of the bubble traps is inserted directly into the microfluidic channel inlets without using any fluidic connectors
- Bubbles in Triton X-100 detergent
  - Gently and slowly invert Triton X-100 solution after aliquoting to prevent large bubbles from forming around the meniscus
  - If bubbles have formed in Triton X-100 solution, let the solution sit for an hour to give time for bubbles to dissipate
- Potential for injury when handling hydrochloric acid
  - HCl is highly corrosive. It can cause serious chemical burns to skin, as well as serious eye damage if not handled appropriately.
  - Always wear the following PPE when handling HCl solutions: lab goggles, lab coat, nitrile gloves, long pants, closed-toe splash-resistant shoes.
  - Make sure as little skin as possible is exposed when handling HCl, especially concentrated solutions.
    - If your lab coat sleeves tend to ride up, consider trucking them into your gloves or taping them to your gloves while handling HCl.

### DETAILED PROTOCOL

#### Day 0: preparation

##### Gasket casting

Fabricate the PDMS gasket, as described in the PDMS fabrication SOP.

- **Record oven temperature using an internal oven thermometer: _℃**
- **Record exact cure time: _ h**

#### Day 1: preparation

##### Gasket preparation

- Remove gasket from oven, cut PDMS out of mould, punch all I/O holes, expand bolt holes to oval if necessary, and clean gasket well with tape.
- Leave the gasket in the desiccator overnight to remove dissolved air and reduce the chance of bubble nucleation.
- Note the chip and gasket information in the table below.

|  | Chip information | Gasket information |
| --- | --- | --- |
| Wafer/mould number |  |  |
| Chip number or gasket preparation date |  |  |

##### Setup preparation

- Check that we have all necessary functional pipettes at the biosensor bay.
- Check that we have pipette tips of the proper type to fit the new pipettes and ensure accurate pipetting.
- Check that we have all necessary waste reservoirs ready for use.
- Gather necessary tubes and other consumables.
- Prepare outlet tubing.

##### Reagent preparation

- Label the sides and top and prepare all tubes for:
  - Functionalization
  - Reagent prep
  - Priming
  - Assay
- Prepare and aliquot all non-protein reagents and pipette required buffer volumes into protein solution aliquots to reduce prep time on day 1.
- Remove an aliquot of the regeneration solution from the 4ºC fridge and leave at room temperature overnight to discourage bubble formation.
- Prepare and measure bulk RI of bulk RI aliquots.
  - Filter all solutions as they are pipetted into each reservoir, by placing the cell strainer on top of the tube as the tubes are filled, and pipetting at an angle so the solutions run down the sides of the tubes to reduce the bubble introduction into the fluid.
- Prepare colour-coded tape labels for the holders of the reservoirs that need to be refilled and the aliquots that will refill them, to help avoid confusion during the refill process.

##### Fluigent prep and RBS clean (if not done since last assay)

1. Draft the Fluigent protocol for the experiment
2. Ensure Fluigent outlets are going into RBS waste tubes.
3. Flow a solution of 10% RBS through Reservoir 1 of both channels of the Fluigent at 80 μL/min for 1 hour.
4. Replace the RBS solutions in the first reservoir of each channel with the post RBS Ultrapure water.
5. Flow the post RBS Ultrapure water at 500 mBar for at least 1 hour.
6. Check the pressures to see if they are stable or increasing

##### Chip and fluidics configuration

| Backwards GC/FA# from layout summary | Our grating coupler and FA # (FA coming from top) | Detector Slot (Data Channel) | Group 1 | | | | Group 2 | | | |
| --- | --- | --- | --- | --- | --- | --- | --- | --- | --- | --- |
|  |  |  | Fluigent channel | Gasket channel | Code | Spot | Fluigent channel | Gasket channel | Code | Spot |
| **1** | **13** | **12 (N77 4)** | 2 | Top (closer to GCs) | R4C11 | Upstream spot 1 | **2** | **Top** | **R4C3** | **Down- stream spot 3** |
| 2 | 12 | 11 (N77 3) | 1 | Bottom (further from GCs) | R3C11 | Upstream spot 1 | 1 | Bottom | R3C3 | Down- stream spot 3 |
| **3** | **11** | **10 (N77 2)** | 2 | Top | R4C12 | Upstream spot 1 | **2** | **Top** | **R4C4** | **Down- stream spot 3** |
| 4 | 10 | 9 (N77 1) | 1 | Bottom | R3C12 | Upstream spot 1 | 1 | Bottom | R3C4 | Down- stream spot 3 |
| **5** | **9** | **8** | 2 | Top | R4C6 | Middle spot 2 | **2** | **Top** | **R4C8** | **Middle spot 2** |
| 6 | 8 | 7 | 1 | Bottom | R3C6 | Middle spot 2 | 1 | Bottom | R3C8 | Middle spot 2 |
| 7 | 7 | N/A, Laser Input |  |  |  |  |  |  |  |  |
| **8** | **6** | **6** | 2 | Top | R4C1 | Downstream spot 3 | **2** | **Top** | **R4C7** | **Middle spot 2** |
| 9 | 5 | 5 | 2 | Top | R4C2 | Downstream spot 3 | 1 | Bottom | R3C7 | Middle spot 2 |
| **10** | **4** | **4** | 1 | Bottom | R3C1 | Downstream spot 3 | **2** | **Top** | **R4C9** | **Upstream spot 1** |
| 11 | 3 | 3 | 1 | Bottom | R3C2 | Downstream spot 3 | 1 | Bottom | R3C9 | Upstream spot 1 |
| **12** | **2** | **2** | 1 | Bottom | R3C5 | Middle spot 2 | **2** | **Top** | **R4C10** | **Upstream spot 1** |
| 13 | 1 | 1 | 2 | Top | R4C5 | Middle spot 2 | 1 | Bottom | R3C10 | Upstream spot 1 |

##### Characterise the bare chip to assess its optical performance

Aligning the fiber array: follow our setup documentation to couple to the loopback of group 1

1. Check and, if needed, adjust the levelness of the stage.
2. Place the photonic chip on the mounting plate and then onto the stage.
3. Align the camera views to the FA.
4. Open the instrument control in PyOptomip so that you can control the stage and the laser.
5. Bring the FA and the photonic chip close together in the X,Y, and Z coordinates. Begin with small steps to make sure the PyOptomip is working correctly and the stage is moving in the direction you expect.
6. Once the Photonic chip is close to the FA, use the cameras and very small step sizes to align the grating couplers of the Group 1 alignment structures to the FA as closely as possible.
   a. Double-check the stage rotation to make sure that the line of grating couplers and the fibre array are parallel.
7. Use the fine align function in PyOptomip to further improve the detected power.

Measure spectra:

1. Once aligned, use the sweep function in PyOptomip to measure the loopback spectrum of Group 1. Save the spectrum in .png and .fig formats. Add images to the table below and save the data to Google Drive.
2. Move forward **200 µm** to the devices
3. Pipette water droplets on the resonators
4. Acquire a sweep and save the image and data to Google Drive. Add the images to the table below.
5. Repeat to characterize next group 2 (move by **3090 μm** **in small steps** to move between devices (300, 300, 300, 300, 300, 300, 300, 300, 300, 300, 90 μm). Each movement, check that the FA and the chip are not moving closer together in Z.
6. Repeat measurements if we run into issues

*Spectra of Groups 1-4 before surface modification*

| **Loopback Powers (0 dBm input power)** | |
| --- | --- |
| Group 1 | Group 2 |
| **Resonance Spectra (0 dBm input power)** | |
| Group 1 | Group 2 |

Be sure to note the chip information in the Day 1 Chip/Gasket information table

##### PDA-coat the SiP chip

1. Clean the chip(s) with acetone, IPA, and ddW
   a. Fill three crystallization dishes with
      i. Dish 1: ~30 mL of acetone
      ii. Dish 2: ~30 mL of isopropanol (IPA)
      iii. Dish 3: ~100 mL of ultrapure water (ddW)
   b. Use the solvent-resistant PELCO tweezers to transfer the chip(s) into the acetone bath (dish 1). Agitate gently for ~30 seconds.
   c. Swiftly transfer the chip(s) to the IPA bath (dish 2), maintaining some liquid acetone on the chip surface during this transfer to avoid acetone drying on the surface, which could leave residue. Agitate gently for ~30 seconds.
   d. Swiftly transfer the chip(s) to the ddW bath (dish 3), maintaining some liquid IPA on the chip surface during this transfer. Agitate gently for ~30 seconds.
   e. Remove the chip(s) from the ddW bath and dry with N_2_ gas.
2. Weigh ≥60 mg of dopamine HCl onto a piece of weigh paper. Pour the dopamine into a 100-mL beaker.
3. Record the actual mass of dopamine in the table below.
4. Calculate the volume of Tris buffer required to achieve a 2 mg/mL solution concentration (≥30 mL).

*Dopamine solution preparation for SiP chip modification*

| Actual mass of dopamine HCl, *m* (mg) | Required volume of Tris ($V=\frac{m}{2 mg/mL}$) |
| --- | --- |

5. Use a serological pipette to transfer the required volume of Tris into the beaker with the dopamine.
6. Working swiftly, add a stir bar to the beaker, cover with parafilm to minimize exposure to air, and stir at ~250-300 rpm on a stir plate until the dopamine is fully dissolved.
7. Using a serological pipette, transfer 30 mL of the dopamine solution into a clean glass crystallization dish. Discard the excess solution in a labeled waste bottle.
8. Place the dish on a magnetic stir plate.
9. Drop a small pill-sized magnetic stir bar into the dish. Position the dish such that the stir bar is held off-center to allow plenty of room for the chip(s).
10. Turn on the stir plate and adjust the speed such that the stir bar spins cleanly on an axis without jumping around the dish, while keeping the stir speed relatively low to avoid excessive turbulence.
11. Transfer the chip(s) into the crystallization dish using clean soft-tip tweezers. Place the chip(s) as far away from the stir bar as possible (see photo below).

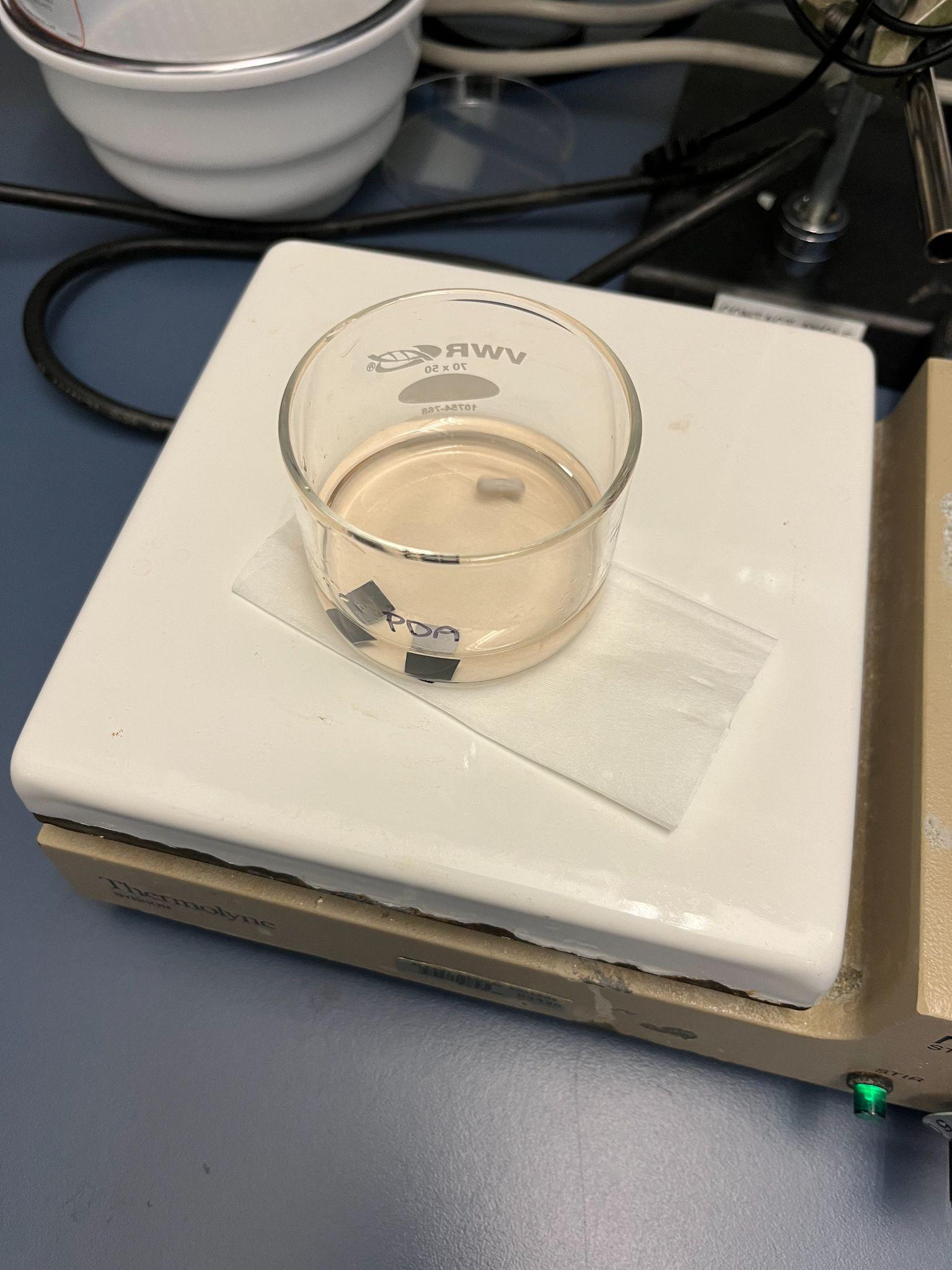

12. Do not cover the reaction vessel. Leave to react at room temperature for 30 minutes, stirring gently the entire time. During this time, the solution will become increasingly dark in colour, indicating oxidation of the dopamine HCl.
13. Rinse and dry the chip(s).For each chip,
    a. Remove the chip from the dopamine solution using tweezers.
    b. Thoroughly rinse the chip(s) with ddW:
       i. Hold the chip with tweezers above a beaker and rinse using ~10 mL of ddW dispensed from a squeeze bottle.
       ii. Place the chip on a piece of cleanroom wipe. Hold the chip steady with tweezers and dry the surface with nitrogen gas.
           1. When drying, make sure to prevent water droplets from touching the tweezers and re-contacting the chip surface.
       iii. Place the chip in a safe place (e.g., a small cleanroom wipe-lined petri dish) until ready for use.
14. Dispose of all liquid waste in the labeled PDA solution waste container.

##### Characterise the PDA-coated chip

Aligning the fiber array: follow our setup documentation to couple to the loopback of group 1

1. Check and, if needed, adjust the levelness of the stage.
2. Place the photonic chip on the mounting plate and then onto the stage.
3. Align the camera views to the FA.
4. Open the instrument control in PyOptomip so that you can control the stage and the laser.
5. Bring the FA and the photonic chip close together in the X,Y, and Z coordinates. Begin with small steps to make sure the PyOptomip is working correctly and the stage is moving in the direction you expect.
6. Once the Photonic chip is close to the FA, use the cameras and very small step sizes to align the grating couplers of the Group 1 alignment structures to the FA as closely as possible.
   a. Double-check the stage rotation to make sure that the line of grating couplers and the fibre array are parallel.
7. Use the fine align function in PyOptomip to further improve the detected power.

Measure spectra:

15. Once aligned, use the sweep function in PyOptomip to measure the loopback spectrum of Group 1. Save the spectrum in .png and .fig formats. Add images to the table below and save the data to Google Drive.
16. Move forward **200 µm** to the devices
17. Pipette water droplets on the resonators
18. Acquire a sweep and save the image and data to Google Drive. Add the images to the table below.
19. Repeat to characterize next group 2 (**move by 3090 μm in small steps** to move between devices (300, 300, 300, 300, 300, 300, 300, 300, 300, 300, 90 μm). Each movement, check that the FA and the chip are not moving closer together in Z.
20. Repeat measurements if we run into issues
21. If needed, raise the FA in Z, turn the chip around, and repeat steps 1-7 for Groups 3 and 4

Store the coated chip(s) in the nitrogen storage box or in a dark cupboard overnight.

*Spectra of groups 1-2 after PDA coating*

| **Loopback Powers (0 dBm input power)** | |
| --- | --- |
| Group 1 | Group 2 |
| **Resonance Spectra (0 dBm input power)** | |
| Group 1 | Group 2 |

#### Day 2: run assay

Update setup sign-in form and experiment tracking sheet.

Follow our Setup Training SOP for all steps.

##### Antibody Spotting:

Reagent preparation details and calculations can be found in the Reagent Prep and Protocol spreadsheet. Prepare non-protein components of functionalization reagents before the day of the assay. Add in protein components on the day of the assay.

*Functionalization reagent prep checklist:*

| Solution | Non-protein components prepared | Protein added | Tubes | Vol. per tube | Stock (S) or functionalization reagent (R)? |
| --- | --- | --- | --- | --- | --- |
| 2X spotting buffer stock solution (0.015% Triton X-100, 30% glycerol in PBS) | Prepared during [Trial 1](https://docs.google.com/spreadsheets/d/1RwReaedv423ecuZpca4e2hZWsXUwbLkr9jwWSpzPcTI/edit?gid=659327382#gid=659327382). | | | | |
| Capture (IL-8) spotting solution |  |  | 1 × 600-µL microcentrifuge tube | 5 µL | R |
| Control (BSA) spotting solution |  |  | 1 × 600-µL microcentrifuge tube | 10 µL | R |
| 200 mg/mL BSA stock in PBS | Prepared on 2024-08-09. See aliquot log for details. | | | | |
| 20 mg/mL BSA blocking solution in PBS |  |  | 1 × 5-mL Eppendorf tube | 3.5 mL | R |

###### Prepare the antibody spotting solution

Refer to the reagent prep spreadsheet. Fill in checklists as you complete each step.

*Capture antibody spotting solution preparation*

| **IL-8 capture antibody spotting solution** | | **Added?** |
| --- | --- | --- |
| Conc. IL-8 cAb stock (µg/mL) | 480 |  |
| Conc. spotting buffer stock (x) | 2 |  |
| Req'd conc. IL-8 cAb (µg/mL) | 240 |  |
| Req'd conc. spotting buffer stock (x) | 1 |  |
| Req'd tot. vol IL-8 cAb spotting sol'n (µL) | 5 |  |
| **Prepare 0.6 mL tube** | | FALSE |
| Vol. IL-8 cAb stock to add (µL) | 2.50 | FALSE |
| Vol. spotting buffer stock to add (µL) | 2.50 | FALSE |
| Vol. PBS to add (µL) | 0.00 | FALSE |
| Triturate | | FALSE |

*Reference BSA spotting solution preparation*

| **Reference BSA spotting solution** | | **Added?** |
| --- | --- | --- |
| Conc. BSA stock (mg/mL) | 200 |  |
| Conc. spotting buffer stock (x) | 2 |  |
| Req'd conc. BSA (mg/mL) | 20 |  |
| Req'd conc. spotting buffer stock (x) | 1 |  |
| Req'd tot. vol BSA spotting sol'n (µL) | 10 |  |
| **Prepare 0.6 mL tube** | | FALSE |
| Vol. BSA stock to add (µL) | 1.00 | FALSE |
| Vol. spotting buffer stock to add (µL) | 5.00 | FALSE |
| Vol. PBS to add (µL) | 4.00 | FALSE |
| Triturate | | FALSE |

Prepare the spotting solution and blocking control solution according to the tables above:

1. Use the buffer stock previously made.
2. Prepare the spotting solutions.
   a. Label two 600 µL microcentrifuge tubes with the solution names.
   b. Add the required spotting buffer followed by the required protein stock solution volumes. Triturate thoroughly.

###### Spot the SiP chip with antibody solution and incubate

1. Line a small petri dish with a few rounds of cleanroom wipe. Dampen the cleanroom wipe with ddW. Add enough ddW to saturate the cleanroom wipe without causing any pooling of liquid.
2. Use soft tip tweezers to transfer the SIP chip into the petri dish, sensor side facing up.
3. Fill a p2 pipette with 1 µL of antibody spotting solution.
4. Carefully deposit the capture antibody solution over the upstream sensors, using the Aven Tools microscope to help position the droplet dispensing. Try as much as possible not to touch the chip with the pipette tip.
   a. Press on the plunger partway, until there is a small droplet at the end of the pipette tip.
   b. Carefully bring the pipette tip down to the chip until the droplet touches the chip surface.
   c. If all sensors are not covered and there is room, you can try to add additional volume to cover the sensors.
5. Repeat step 4 for the downstream sensors (capture antibody spotting solution) and middle sensors (BSA reference spotting solution).
6. Take an image with the Aven Tools microscope to record droplet positions. 📷
   a. **Time of spotting**: **_ am**
   b. **Volume spotted on upstream group 1: _ μL**
   c. **Volume spotted on middle group 2: _ μL**
   d. **Volume spotted on downstream group 3: _ μL**
7. Close the petri dish, and leave to incubate for 1 hour at room temperature. Take another image every ~10 minutes using the Aven Tools microscope. 📷
   a. If droplets start to grow towards each other, try opening the petri dish to reduce humidity. Continue monitoring to make sure the droplets do not dry out.
8. Take an image with the Aven Tools microscope of the droplets covering the sensor groups at the end of the incubation, to look for any changes in droplet size or position.
   a. **Start time of rinse: _ am**
9. After the 1-hour incubation time is complete, rinse the chip with PBS
   a. Use soft tip tweezers to hold the chip above a clean petri dish. Use a 1 mL pipette to rinse the chip surface with 3 mL of PBS.
   b. Fill two wells of a 12-well plate with 4 mL of PBS.
   c. Transfer the chip to the well. Leave for 3 minutes with periodic gentle agitation.
   d. Transfer the chip to the second well. Leave for 3 minutes with periodic gentle agitation.

##### Blocking:

Refer to the reagent prep spreadsheet. Fill in checklists as you complete each step.

###### Prepare blocking solution

*BSA blocking solution preparation details*

| **BSA blocking solution** | | **Added?** |
| --- | --- | --- |
| Conc. BSA stock (mg/mL) | 200 |  |
| Req'd conc. BSA (mg/mL) | 20 |  |
| Req'd tot. vol. BSA blocking sol'n (µL) | 3500 |  |
| **Prepare 5 mL tube** | | FALSE |
| Vol. BSA stock to add (µL) | 350 | FALSE |
| Vol. PBS to add (µL) | 3150 | FALSE |
| Triturate | | FALSE |

Prepare the BSA blocking solution according to the table above.

1. Prepare the 20 mg/mL BSA blocking solution (do this while the spotting solutions are being incubated on the chip).
   a. Label a 600-µL microcentrifuge tube for the blocking solution.
   b. Thaw a BSA stock solution aliquot and triturate before use.
   c. Add the required volume of PBS to the tube, followed by the 200 mg/mL BSA stock solution. Triturate thoroughly.

###### Block the sensor surface

1. Add the BSA blocking solution to a well of the 12-well plate.
2. Transfer the chip to the BSA solution to incubate for 1 hour at room temperature.
   a. Incubation start time: **_ am**
3. After the incubation time is complete, rinse the chip: **_ am**
   a. Use soft tip tweezers to hold the chip above a clean petri dish. Use a 1 mL pipette to rinse the chip surface with 3 mL of PBS.
   b. Fill two wells of a 12-well plate with 4 mL of PBS.
   c. Transfer the chip to the first well. Leave for 5 minutes with periodic gentle agitation.
   d. Transfer the chip to the second well. Leave for 5 minutes with periodic gentle agitation.
4. Take note of the following
   a. Ambient temperature: **_ C**
   b. Ambient humidity: **_%**

##### Binding Assay

###### Reservoir setup

Calculations and further details about reagent preparation and reservoir setup can be found in the reagent prep spreadsheet.

| Reservoir # | Reagent type | Ch. 1 solution | Ch. 2 solution | Priming | Required vol. per ch. before excess (µL) | Req'd vol. per ch. with excess for assay (µL) | Check fill/refill before assay start to at least (µL) | Refill after 2nd binding cycle to at least... (µL) |
| --- | --- | --- | --- | --- | --- | --- | --- | --- |
| 1 | Ultrapure water | Ultrapure water | Ultrapure water | Ultrapure water | 2550 | 3550 | 3550 |  |
| 2a | Streptavidin-HRP | Streptavidin-HRP, 2 µg/mL | Streptavidin-HRP, 2 µg/mL | Running buffer | 3000 | 3200 | 3200 |  |
| 2b | Bulk RI: PBS, 0.5x | PBS, 0.5x | PBS, 0.5x | – | 2400 | 3400 | – |  |
| 3a | IL-8 sample | IL-8, 3.125 ng/mL in CCM | IL-8, 3.125 ng/mL in CCM | Cell culture medium | 2500 | 2700 | 2700 |  |
| 3b | Bulk RI: PBS, 1x | PBS, 1x | PBS, 1x | – | 1200 | 2200 | – |  |
| 4a | Triton X-100 pre-wetting solution | Triton X-100, 0.3 mM | Triton X-100, 0.3 mM | Triton X-100 pre-wetting solution | 360 | 1360 | – |  |
| 4b | Running buffer | PBS-BSA | PBS-BSA | Running buffer | 18900 | 19900 | 13000 | 13000 |
| 5 | BSA/CCM challenge | BSA, 1 mg/mL in CCM | BSA, 1 mg/mL in CCM | BSA/CCM challenge | 3600 | 4600 | 4600 |  |
| 6 | IL-8 detection antibody | IL-8 detection antibody, 1 µg/mL | IL-8 detection antibody, 1 µg/mL | Running buffer | 5400 | 5600 | 5600 |  |
| 7 | CCM | CCM | CCM | Cell culture medium | 500 | 700 | 700 |  |
| 8 | Unused in assay | PBS, 1x | PBS, 1x | PBS, 1x | 0 | 500 | 500 |  |
| 9 | Unused in assay | PBS, 1x | PBS, 1x | PBS, 1x | 0 | 500 | 500 |  |
| 10 | Regeneration solution | 10 mM glycine, 160 mM NaCl in water, pH 2.2 | 10 mM glycine, 160 mM NaCl in water, pH 2.2 | Regeneration solution | 1050 | 1250 | 1250 |  |

##### Reagent preparation

Refer to the reagent prep spreadsheet. Fill in checklists as you complete each step. Prepare non-protein components of binding assay reagents before the day of the assay. Add in protein components on the day of the assay (this may be done while the chip is being blocked with BSA).

*Triton X-100 solution preparation details*

| **Triton X-100 pre-wetting solution** | | **Added?** |
| --- | --- | --- |
| Mw, Triton X-100 (g/mol) | 625 |  |
| Density, Triton X-100 (g/mL) | 1.07 |  |
| Conc. Triton X-100 stock (mM) | 30 | *Use 30 mM stock |
| Req'd conc. Triton X-100 in prewetting sol'n (mM) | 0.3 |  |
| Req'd vol. per Triton X-100 prewetting sol'n channel (µL) | 2860 |  |
| # of channels | 2 |  |
| Req'd tot. vol. Triton X-100 prewetting sol'n (µL) | 5720 |  |
| **Prepare 15 mL prep tube** | | FALSE |
| **Prepare 2 × 15 mL assay tubes** | | FALSE |
| Vol. Triton X-100 stock to add (µL) | 57 | FALSE |
| Vol. PBS to add (µL) | 5663 | FALSE |
| Triturate | | FALSE |
| Vol. to split | 2860 | FALSE |

*PBS-BSA running buffer preparation details*

| **Running buffer (PBS-BSA)** | | **Added?** |  |
| --- | --- | --- | --- |
| Stock conc., BSA (mg/mL) | 200 |  |  |
| Req. conc., BSA (mg/mL) | 0.1 |  |  |
| Req'd vol. per running buffer channel, including priming (mL) | 19.9 | *Includes priming |  |
| Additional req'd vol. per channel for priming (mL) | 2 | 2 reservoirs/ch, 1 mL priming vol |  |
| # of channels | 2 |  |  |
| Additional req'd vol. for other solution prep (mL) | 17.531 |  |  |
| Tot. req'd vol. (mL) | 61.331 |  |  |
| Vol. to prep (mL) | 80 |  |  |
| Prepare 2 × 50 mL prep tubes | | FALSE |  |
| Prepare 2 × 15 mL assay tubes | | FALSE |  |
| Prepare 4 × 15 mL priming tubes | | FALSE |  |
| Prepare in 2 vessels: | | Added tube 1? | Added tube 2? |
| Vol. BSA stock to add (µL) | 20 | FALSE | FALSE |
| Vol. PBS to add (mL) | 39.980 | FALSE | FALSE |
| Pour one into the other to mix (x2) | | FALSE | FALSE |

*CCM preparation details*

| **Cell culture medium (CCM)** | | **Added?** |
| --- | --- | --- |
| Stock conc., FBS (x) | 1 |  |
| Stock conc., anti-anti (x) | 100 |  |
| Stock conc., L-glutamine (mM) | 200 |  |
| Req'd conc., FBS (x) | 0.1 |  |
| Req'd conc., anti-anti (x) | 1 |  |
| Req'd conc., L-glutamine (mM) | 2 |  |
| Req'd vol. per CCM channel, including priming (mL) | 1.7 |  |
| Additional req'd vol. per channel for priming (mL) | 1 | 1 reservoir/ch, 1 mL priming vol |
| # of channels | 2 |  |
| Additional req'd vol. for other solution prep (mL) | 17.398 | IL-8 + BSA/CCM challenge |
| Tot. req'd vol. (mL) | 22.798 |  |
| Vol. to prep (mL) | 50 |  |
| **Prepare 50 mL prep tube** | | FALSE |
| **Prepare 2 × 2 mL assay tubes** | | FALSE |
| Vol. FBS stock to add (mL) | 5 | FALSE |
| Vol. anti-anti stock to add (mL) | 0.5 | FALSE |
| Vol. L-glutamine stock to add (mL) | 0.5 | FALSE |
| Vol. DMEM to add (mL) | 44 | FALSE |
| Triturate | | FALSE |
| Vol. to split/filter (mL) | 1.7 | FALSE |

*BSA/CCM challenge solution preparation details*

| **BSA/CCM challenge** | | **Added?** |
| --- | --- | --- |
| Stock conc., BSA (mg/mL) | 200 |  |
| Req'd conc., BSA (mg/mL) | 1 |  |
| Req'd vol. per channel (µL) | 6100 |  |
| # of channels | 2 |  |
| Tot. req'd vol. (µL) | 12200 |  |
| **Prepare 15 mL prep tube** | | FALSE |
| **Prepare 2 × 15 mL assay tubes** | | FALSE |
| Vol. BSA stock to add (µL) | 61.0 | FALSE |
| Vol. CCM to add (µL) | 12139.0 | FALSE |
| Triturate | | FALSE |
| Vol. to split (µL) | 6100 | FALSE |

*IL-8 sample solution preparation details*

| **IL-8** | | **Added?** |
| --- | --- | --- |
| Stock conc., IL-8 (ng/mL) | 120 |  |
| Req'd conc., IL-8 (ng/mL) | 3.125 |  |
| Req'd vol. per channel (µL) | 2700 |  |
| # of channels | 2 |  |
| Tot. req'd vol. (µL) | 5400 |  |
| **Prepare 15 mL prep tube** | | FALSE |
| **Prepare 2 × 15 mL assay tubes** | | FALSE |
| Vol. IL-8 stock to add (µL) | 141 | FALSE |
| Vol. CCM to add (µL) | 5259 | FALSE |
| Triturate | | FALSE |
| Vol. To split/filter (µL) | 2700 | FALSE |

*IL-8 detection antibody solution preparation details*

| **IL-8 detection antibody** | | **Added?** |
| --- | --- | --- |
| Stock conc., IL-8 detection antibody (µg/mL) | 200 |  |
| Req'd conc., detection antibody (µg/mL) | 1 |  |
| Req'd vol. per channel (µL) | 5600 |  |
| # of channels | 2 |  |
| Tot. req'd vol. (µL) | 11200 |  |
| **Prepare 15 mL prep tube** | | FALSE |
| **Prepare 2 × 15 mL assay tubes** | | FALSE |
| Vol. IL-8 detection antibody stock to add (µL) | 56 | FALSE |
| Vol. PBS-BSA to add (µL) | 11144 | FALSE |
| Triturate | | FALSE |
| Vol. to split (µL) | 5600 | FALSE |

*SA-HRP solution preparation details*

| **Streptavidin-HRP** | | **Added?** |
| --- | --- | --- |
| Stock conc., SA-HRP (µg/mL) | 1000 |  |
| Req'd conc., SA-HRP (µg/mL) | 2 |  |
| Req'd vol. per channel (µL) | 3200 |  |
| # of channels | 2 |  |
| Tot. req'd vol. (µL) | 6400 |  |
| **Prepare 15 mL prep tube** | | FALSE |
| **Prepare 2 × 15 mL assay tubes** | | FALSE |
| Vol. SA-HRP stock to add (µL) | 12.8 | FALSE |
| Vol. PBS-BSA to add (µL) | 6387.2 | FALSE |
| Triturate | | FALSE |
| Vol. to split (µL) | 3200 | FALSE |

*Regeneration solution preparation details*

| **Regeneration buffer (prepared 2024-07-04)** | | **Added?** |
| --- | --- | --- |
| Conc., glycine (mM) | 10 |  |
| Conc., NaCl (mM) | 160 |  |
| Measured pH | 2.2 |  |
| Tot. req'd vol. (both channels) (µL) | 4500 |  |
| **Prepare 15 mL prep tube** | | FALSE |
| **Prepare 2 × 2 mL assay tubes** | | FALSE |
| Req'd vol. per channel (µL) | 2250 | FALSE |

*Bulk RI solution RIs and preparation details*

| **Bulk RI solutions** | | | | | | | | | |
| --- | --- | --- | --- | --- | --- | --- | --- | --- | --- |
| PBS conc. (x) | Vol. to prepare (mL) | Vol. 1x PBS to add (mL) | Vol. water to add (mL) | Meas. RI 1 | Meas. RI 2 | Meas. RI 3 | Ave. meas. RI | Step size (RIU) | Prepared? |
| 1 | 50 | 50 | 0 |  |  |  | #DIV/0! | #DIV/0! | FALSE |
| 0.5 | 50 | 25 | 25 |  |  |  | #DIV/0! | #DIV/0! | FALSE |
| 0 | 50 | 0 | 50 |  |  |  | #DIV/0! | -- | FALSE |

1. Prepare the binding assay reagents according to the tables above. Filter all solutions through 40 μm cell strainers as they are pipetted into the reservoir tubes.
   a. Prepare the Triton X-100 pre-wetting solution
      i. Add the required volume of PBS to a labelled 15-mL Falcon tube.
      ii. Add the required volume of 30 mM Triton X-100 stock solution.
      iii. Triturate thoroughly.
      iv. Split & filter the solution into the prepared assay reservoir tubes.
   b. Prepare the PBS-BSA (0.1 mg/mL BSA) running buffer solution.
      i. Add the required amount of PBS to the 50 mL tubes.
      ii. Add in the required volume of BSA stock.
      iii. Triturate thoroughly.
      iv. Filter the solution into the prepared priming & assay reservoir tubes.
   c. Prepare the CCM
      i. Add the required amount of DMEM to a 50-mL tube.
      ii. Add the required amount of FBS stock.
      iii. Add the required amount of anti-anti stock.
      iv. Add the required amount of L-glutamine stock.
      v. Triturate thoroughly.
      vi. Filter the solution into the prepared priming & assay reservoir tubes.
   d. Prepare the BSA/CCM challenge solution.
      i. Add the required amount of CCM to a 15-mL tube.
      ii. Add the required amount of BSA stock.
      iii. Triturate thoroughly.
      iv. Split & filter the solution into the prepared assay reservoir tubes.
   e. Prepare IL-8 solution
      i. Add the required amount of CCM to a 15 mL tube.
      ii. Add in the required volume of IL-8 stock.
      iii. Triturate thoroughly.
      iv. Split & filter the solution into the prepared assay reservoir tubes.
   f. Prepare IL-8 detection antibody solution
      i. Add the required amount of PBS-BSA running buffer to a 15 mL tube.
      ii. Add in the required volume of IL-8 detection antibody stock.
      iii. Triturate thoroughly.
      iv. Split & filter the solution into the prepared assay reservoir tubes.
   g. Prepare SA-HRP amplification solution
      i. Add the required amount of PBS-BSA running buffer to a 15 mL tube.
      ii. Add in the required volume of IL-8 detection antibody stock.
      iii. Triturate thoroughly.
      iv. Split & filter the solution into the prepared assay reservoir tubes.
   h. Prepare the regeneration solution.
      i. Filter 1.5 mL of the existing regeneration solution from the aliquot kept at RT overnight into the labelled assay reservoir tubes.
   i. Prepare the bulk RI solutions
      i. Use the refractometer to measure the refractive index of each of the bulk RI solutions and record the results in the table above.
      ii. Filter the solutions into the prepared assay reservoir tubes.
2. Zero the pressure on the Fluigent and block the outlet tubing.
3. Connect the fluidic reservoirs to the priming solutions as described in the reservoir setup table.

###### Prime the fluidic channels

1. Do not connect the bubble traps until after priming, in case there is residual IPA in the lines from the previous ramp-down that could wet the bubble trap membranes.
2. Insert the effluent PEEK tubing for each channel in separate non-hazardous waste tubes.
3. Double-check all reservoirs:
   - Is the correct priming solution in each reservoir position?
   - Is there enough volume with excess remaining in each reservoir after priming?
   - Are there any tiny bubbles trapped at the bottom of the reservoir tubes? If so, tap the tube (e.g., with a marker or scissors) to release the bubbles. Try not to tap hard enough to damage the tubes or cause the liquid to splash up to the cap.
4. Run the priming protocol on Oxygen so that each reservoir that will be used has flow at 750 mBar for 2 mins.
   - When preparing the priming protocol, prime the Triton X-100 reservoirs separately (make the automated priming protocol go from reservoir 3→1, 10→5 and then run a second protocol to prime reservoir 4 (Triton). Move the PEEK tubing outlets of each channel from the non-hazardous waste tubes to Triton X-100 waste tubes when priming reservoir 4.
   - Keep an eye on the measured flow rates to make sure that the tubing is effectively primed.
     - Flow rate should be saturated at 120 μL/min.
     - During flow of each reservoir we should see a characteristic pattern of high flow (fluid in the M-switch-to-flow-sensor tubing) → low flow (air bubble that filled reservoir-to-M-switch tubing) → high flow (after the fluid from the reservoir reaches the flow sensor).
   - After priming, double-check the Fluigent log file to make sure that the correct flow rates were reached.
5. Set flow rate to 1-5 uL/min after priming the Triton X-100, to allow small crevices in the bubble trap lines to be better wetted and to reduce Triton waste.
6. Connect the bubble traps to the appropriate Fluigent channel line(s). Use the coloured tape indicators on each tubing connection to avoid confusion.
7. Wait until the fluid fills the bubble trap lines, while setting up the system to start prewetting. Increase the flow rate if necessary.
8. Turn on the bubble trap vacuum. Watch for fluid in the bubble trap vacuum lines. If fluid is observed, this means that the bubble trap membrane has been wetted, is compromised, and needs to be replaced.
9. Start pre-wetting the gasket.

##### Prepare chip-gasket assembly

1. Take an image of the functionalized photonic chip with the Aven Tools microscope prior to assembly. 📷
2. Leave the gasket in the desiccator under vacuum until just prior to use.
3. Immediately before mounting the gasket on the chip (within ~30 mins or prewetting), plasma-treat the gasket.
   a. Plasma-treat the gasket with air plasma using the main lab plasma chamber for 1 min 15 sec at 400 mTorr and high power.
   b. **Pressure**: **_ mTorr**
   c. **Time**: **_ pm**
4. Working under the Aven Tools microscope, mount the gasket onto the mounting plate over top of the photonic chip ensuring proper alignment of the channels, and place the mounted gasket on the Maple Leaf stage setup.
5. Tighten the bolts to the “sweet-spot” (gently tightening) of making the gasket secure and leak-free but without causing too much channel distortion.
6. Take a set of images of the assembled gasket and chip using the Aven Tools microscope. 📷
   a. Take an overall image of the chip and aligned gasket 📷
   b. Take zoomed-in photos of each of the resonator groups to show alignment 📷
   c. Take zoomed-in images of the GCs 📷
7. Rinse the grating couplers:
   a. Use a piece of scotch tape to cover the inlets of the gasket to prevent any liquid from accidentally entering.
   b. Use a 1 mL pipette to flow ~2 mL of excess regeneration solution over the grating couplers after mounting the gasket (if there are no leaks and we are careful to not let any solution splatter near the inlets, this should not affect any of the resonators).
   c. Use a 1 mL pipette to flow ~3 mL of ddW over the grating couplers.
   d. Dry the grating couplers by tilting the chip-gasket assembly and using a Kimwipe to wick up excess liquid at the edge of the chip. Use some compressed air or nitrogen to dry off any excess liquid.
   e. Inspect the grating couplers to make sure they look clean and take another set of images of the assembled gasket and chip. 📷
      i. Take an overall image of the chip and aligned gasket 📷
      ii. Take zoomed-in photos of each of the resonator groups to show alignment 📷
      iii. Take zoomed-in images of the GCs 📷

###### Pre-wetting

1. Insert dried outlet tubing (>20 mins airflow to dry) into the outlet ports, verifying that the PEEK tubing does not extend to the bottom of the PDMS (chip surface). Block the outlet tubing.
2. Double-check all reservoirs:
   - Is the correct solution in each reservoir position?
   - Is there enough volume with excess remaining in each reservoir after priming?
   - Are there any tiny bubbles trapped at the bottom of the reservoir tubes? If so, tap the tube (e.g., with a marker or scissors) to release the bubbles. Try not to tap hard enough to damage the tubes or cause the liquid to splash up to the cap.
3. Confirm that the Fluigent channels are filled with Triton X-100 prewetting solution and the outlet tubes are in a Triton waste tube and zero the pressures on both channels.
4. Hold the bubble trap outlets up to produce some gravitational backflow to make sure that we do not accidentally introduce fluid at a faster-than-intended flow rate due to gravity flow. Aim for ~5s of backflow (backflow rate is typically around 5-6 μL/min, so this duration of backflow equates to ~30-60s of slow flow at 1 μL/min before the fluid reaches the chip).
5. Then, block the Tygon tubings that connect the bubble traps to prevent further backflow while connecting the inlet tubings, and confirm that the Fluigent measures zero flow. Dry the bubble trap tubing outlets with a Kimwipe and ensure the tubing ends are clean prior to connecting to the chip.
6. Set up the PixelLink video software to take a time-lapse capture of the flow every 5s. Ensure the top camera is focused on the channels.
7. Connect the Fluigent channel 1 and Fluigent channel 2 bubble trap outlet tubing to the inlets on the two channels of the microfluidic chip. Verify that the tubings are inserted ~2 mm into the PDMS but do not extend all the way down to the chip.
8. Unblock the bubble trap and outlet tubings and immediately begin Fluigent protocol with the delivery of Triton X-100 at 1 μL/min and visualise slow wetting of channel #1 and #2 (any wait after unblocking will introduce further backflow).
9. After the microfluidic channels are wetted and the menisci are visible in the outlet Tygon tubing, increase the flow rate to 30 μl/min and flow Triton X-100 for ~5-10 minutes to ensure Triton X-100 has sufficiently flowed through the entire system. Visualise location of meniscus in the outlet Tygon tubing and watch for bubbles in the channels. Then, switch to PBS-BSA running buffer and proceed to alignment.
   - **Time the prewetting solution contacted the channels**:
     i. **Channel 1: _ pm**
     ii. **Channel 2: _ pm**

###### Align fibre array and select best resonator response

After pre-wetting,

1. Start PBS-BSA flow at 30 µL/min through both channels. Keep the outlet tubing in the Triton X-100 waste containers during this rinse step. Ensure that the PBS-BSA flows for >20 minutes to rinse out residual Triton X-100.
2. While PBS-BSA is flowing over the sensors in both channels, align the FA to the chip and select a group (from groups 1 & 2).

Aligning the Fibre Array:

1. Align the camera views to the FA.
2. Open the instrument control in PyOptomip so that you can control the stage and the laser.
3. Bring the FA and the photonic chip close together in the X,Y, and Z coordinates. Begin with small steps to make sure the PyOptomip is working correctly and the stage is moving in the direction you expect.
4. Once the Photonic chip is close to the FA, use the cameras and very small step sizes to align the grating couplers of Group 1’s alignment structures to the FA as closely as possible.
5. Use the fine align function in PyOptomip to further improve the detected power.
6. Sweep and save the alignment structure spectra
7. Move forward **200 µm** to the devices
8. Adjust as needed and then do another fine align.

Determining the Best Resonator Response:

22. Once aligned, use the sweep function in PyOptomip to determine the resonator response of the first device (device 6). Take a screenshot of the graph and attach it in the table below.
23. Repeat to characterize next group 2 (move by **3090 μm** **in small steps**
24. Lower the FA as close as possible to the chip again while being confident that you will not collide with the chip using very small step (100 µm, then 10 µm) sizes.
25. Use small 10 µm movements if needed to bring the power above -50 dBm and then perform a fine alignment. It may be necessary to raise the laser power and decrease the detectors’ initial range if we once again observe high losses from the functionalized chip’s through-port response (like in PDA Spike Trial 1).
26. Use the sweep function in PyOptomip to determine the resonator response of the device. Take a screenshot of the graph and attach it in the table below.
27. Determine which device to use by checking which device demonstrates the greatest quality factor and extinction ratios. Watch for peaks close to (within ~10 dB of) the -70 dBm detector cutoff threshold (below which, PyOptomip replaces measured power with -100 dBm) and choose devices and spectral regions that are not subject to this issue. Including cut-off peaks with -100 dBm values in the data acquisition can make the data difficult or impossible to analyse for peak-tracking as it impacts Lorentzian fitting of the peaks. Try to make sure that peaks in the measured spectra only extend down to ~-60 dBm so that the peaks remain above the threshold power even if the baseline fluctuates due to coupling issues, etc.
28. Realign the fibre array with the alignment structures of the chosen device.
29. Obtain an approximate alignment to the waveguides by moving in the positive Y direction by 337 μm.
30. Lower the FA to within a few micrometres of the chip.
31. Move in the X and Y directions in increments of 5 - 10 µm until the powers are greater than -40 dB.
32. Perform a fine align, and then a sweep (save the sweep image and .mat file).
33. Locate the three adjacent peaks with the best quality factors and greatest extinction ratios within the ~10 dB of the -70 dB cutoff threshold.
34. Perform another sweep using a range that only includes these peaks. Save the image and .mat file again. Enter this sweep range in the .toml file of multisweep.

*Spectra of groups 1 and 2 (specify which group was used, as well as the wavelength range used for sweeps)*

| Loopback Powers (0 dBm) | |
| --- | --- |
| Group 1 | Group 2 |
| Resonator spectra | |
| Group 1 | Group 2 |

###### Run the binding assay

1. Empty the non-hazardous and Triton waste tubes and move the tubing outlets into **separate assay waste containers**.
2. Double-check all reservoirs:
   - Is the correct solution in each reservoir position?
   - Is there enough volume with excess remaining in each reservoir after priming?
   - Are there any tiny bubbles trapped at the bottom of the reservoir tubes? If so, tap the tube (e.g., with a marker or scissors) to release the bubbles. Try not to tap hard enough to damage the tubes or cause the liquid to splash up to the cap.
3. Set up the binding assay Oxygen protocol according to the steps described in the table below and double-check the steps.
   - Both channels should be identical and follow this protocol.
4. Set up the Pixelink software to capture the channel region every 20 seconds for the duration of the binding assay. Ensure that the images are saved to the correct location
5. Start the binding assay Oxygen protocol. Record the start and end time of the protocol.
6. Start multisweep and ensure that sweeps are being saved in the desired folder.
7. **Time protocol started: _ pm**
8. **Refill time**:**_ pm**
   - During the refill, we will need to:
     - Refill running buffer to >13 mL
9. **Reservoir change time:_ pm**
   - During the reservoir change, we will need to:
     - Switch reservoir 2 to 0.5x PBS
     - Switch reservoir 3 to 1x PBS
10. For the refills/reservoir switches:
    - Colour code the tops of these reservoirs and the refill containers with coloured tape to avoid confusion.
    - Set a timer for 2 minutes before the M-switch switches out of PBS-BSA stability reservoir. If this timer goes off, pause the Fluigent protocol (don’t forget to restart after refill complete).
    - With 10 mins left before the next protocol step following the refill/reservoir switch wait time, start the reservoir refill/switch process:
      - In both channels, zero the pressures and block the outlet tubing and bubble trap tubing.
      - One by one, remove each reservoir from the Fluigent and refill/switch as required.
    - Double-check that we have the min. required volumes and that all reservoirs are properly attached.
    - In both channels, unblock the outlet tubing and supply a flow rate of 30 µL/min, ensuring the M-switch is still set to the desired reservoir.

###### Fluidic protocol for binding assay

Please refer to the reagent prep and protocol spreadsheet for the assay protocol.

##### Check-in table

| Experiment checklist table | | | | | | | |
| --- | --- | --- | --- | --- | --- | --- | --- |
| Check-in # | 1 | 2 | 3 | 4 | 5 | 6 | 7 |
| Time |  |  |  |  |  |  |  |
| Date |  | | | | | | |
| Monitor name |  |  |  |  |  |  |  |
| Bubbles in the channel? |  |  |  |  |  |  |  |
| Bubbles in the bubble trap tubing or outlet tubing? |  |  |  |  |  |  |  |
| Bubbles in the reservoirs |  |  |  |  |  |  |  |
| Liquid in the bubble trap vacuum lines or vacuum trap? |  |  |  |  |  |  |  |
| Flow rate at outlet tubes? |  |  |  |  |  |  |  |
| Optical coupling ok? |  |  |  |  |  |  |  |
| Fluigent flow ok? Please note pressures. |  |  |  |  |  |  |  |
| Sufficient fluid remaining? |  |  |  |  |  |  |  |
| Leakage? |  |  |  |  |  |  |  |
| Spectrogram and peak shift plots look ok? |  |  |  |  |  |  |  |
| Multisweep running ok (saving files)? |  |  |  |  |  |  |  |
| Camera image saving ok? |  |  |  |  |  |  |  |
| Upload data to Google Drive? |  |  |  |  |  |  |  |
| Empty waste tubes if necessary |  |  |  |  |  |  |  |
| Notes:   1. | | | | | | | |

#### Day 3: Cleanup

**Switch off the setup devices**

- Ensure that MultiSweep, timelapse capture, and the Fluigent protocol have stopped.
- **Transfer the rest of the sweeps and captures that were obtained overnight.**
- **Transfer the Fluigent log files to the data folder and Google Drive**
- Run Retrospective Analysis and transfer the data.
- Restore the setup (fibre array position and cameras) to the default position for the next user.
- Turn off the stage motors.
- Turn off the N77 module.
- Turn off the TEC temperature controller (just the “TEC On/Off” button – the unit can remain powered on).
- Ensure the laser is off.
- Ensure appropriate ramp-down steps for the optical setup have been followed, as described in our setup documentation.
- Complete the biosensor setup sign-out form and setup experiment planning and tracking checklist.

**Disassemble the microfluidics**

- Disconnect the inlet PEEK tubing from the microfluidic gasket.
- Remove the gasket and mounting plate from the stage
- Allow gravity flow to empty the channels (hold up the gasket with respect to the outlet and watch the water/air interface travel the full length of the outlets to empty them). If gravity flow does not work, use a “withdrawal” syringe to empty the channel via negative pressure at the end of the outlet tube.
- Take a set of images of the chip post-assay using the Aven Tools microscope 📷
  - Take an image of the full chip 📷
  - Take zoomed-in images of each group of resonators 📷
  - Take zoomed-in images of any visible residue or other phenomena 📷
- **Disconnect the outlets from the microfluidics and store.**
- **Store the used chip safely back in the Gel-Pak.**

**Clean the Fluigent for the next users**

- Turn off the bubble trap vacuum.
- Switch all used Fluigent reservoirs to ddW. Flush all used reservoirs with ddW in sequence at **750 mBar for 3 min**.
- **Dispose of the assay waste as biohazardous waste**
- **Decontaminate potentially biohazardous lines**: If CCM was used, we decontaminate these reservoir lines with 70% ethanol:
  - All reservoirs that contained CCM are switched to ultrapure water and use live control or a protocol to flow from each at 750 mbar for 3 minutes into a biohazardous waste container.
    - Water reservoirs connected to artificial urine lines should be marked as biohazardous and disposed of after use as biohazardous waste.
    - The rinse step is necessary to prevent salt precipitation, which could occur if switching directly from a salt-containing solution like PBS to ethanol.
  - Disconnect the bubble traps as they are not compatible with alcohol.
  - All reservoirs that contained CCM are then switched to 70% ethanol and use live control or a protocol to flow from each at 750 mbar for 3 minutes into a non-hazardous waste container.
  - Switch these reservoirs to ultrapure water tubes and flow again from each at 750 mbar for 3 minutes into a non-hazardous waste container.
    - After the water rinse, reconnect the bubble traps.
- **Clean protein residues**: If solutions containing proteins were used in the assay, we rinse those lines with Alconox to remove protein residues.
  - Place the microfluidic inlet tubing (the outlet of the bubble traps) in an “Alconox waste” reservoir (empty the reservoir into the Alconox waste collection vessel if it has liquid in it).
  - Switch all reservoirs that contained protein-containing solutions at any point in the assay to 2% Alconox. Flush these reservoirs in sequence with Alconox at 750 mBar for 3 min.
  - Switch these back to ultrapure water (rinsing off any Alconox droplets from the in-reservoir FEP tubing using an ultrapure water wash bottle before connecting the water reservoir) and flush with ddW at 750 mBar for 3 min.
  - Switch the outlet back to non-hazardous waste and empty the Alconox waste into the Alconox waste vessel (NOT down the drain).
- Move the outlet tubing to an RBS waste tube.
- **Run an RBS clean to keep our flow sensors in good working order:**
  - Run either a short or overnight RBS clean:
    - Place the microfluidic inlet tubing (the outlet of the bubble traps) in an “RBS waste” reservoir (empty the reservoir into the RBS waste collection vessel if it has liquid in it).
    - For an overnight RBS clean:
      - Switch the reservoir 1 tube to 10% RBS-35 in ultrapure water, filled to 11 mL (note: the RBS clean is usually performed from one reservoir only).
      - Flow reservoir 1 at 10 μL/min overnight. 11 mL is sufficient volume to last for 16 hours (10000 μL)/(10 μL/min)*(1h/60 min). **Flow must be stopped by that time so that the system does not dry out with RBS in place**
    - For a short RBS clean:
      - Switch the reservoir 1 tube to 10% RBS in ultrapure water, filled to >6 mL.
      - Flow reservoir 1 at 80 μL/min for 45-75 mins (ideally 1h). 6 mL is sufficient volume to last for 75 mins (6000 μL)=(80 μL/min)*(75 min). **Flow must be stopped by that time so that the system does not dry out with RBS in place**
    - After the RBS clean, rinse the RBS droplets from the inner FEP tube using a wash bottle filled with ultrapure water, and switch reservoir 1 to a post-RBS ultrapure water tube, filled to 13 mL.
    - Supply > 6 mL water flow by constant flow or constant pressure (e.g., 120 μL/min for 1h).
    - Switch the outlet tubing in the waste Falcon tube from RBS waste to regular waste, and empty the RBS waste Falcon tube in the RBS waste vessel (NOT down the drain).
    - Run a priming protocol to rinse reservoir 1 with fresh, clean ultrapure water from a regular (non-post-RBS) ultrapure water reservoir (e.g., 750 mbar for 3 mins).
- If the system is to be used for another experiment, proceed to priming with the reagents for that experiment.
- If not, proceed to ramp down:
  - **Disconnect the bubble traps as they are not compatible with alcohol.**
  - Use the clean air backflow syringe to push air through the bubble trap lines from both sides until the bubble traps are empty (to prevent liquid from sitting in the lines).
  - Zero the Fluigent pressure.
  - Switch all used Fluigent reservoirs to 100% isopropyl alcohol (IPA). Flush all used reservoirs with IPA in sequence at 750 mBar for 2 min.
  - Zero the Fluigent pressure.
  - Switch all used Fluigent reservoirs to air. Flush all used reservoirs with air in sequence at 750 mBar for 2 min.
  - Turn off the Fluigent system and pump.
- Transfer all collected sensor and Fluigent data to the Biosensor Google Drive.
- Empty the compressor tank.
- Empty wastes.
- Close OxyGEN and turn off the Fluigent and pump.
- Update sign-out form and experiment tracking sheet
- Ensure all data (including Fluigent log files) has been transferred to our group’s drive and backed up.

### EXPERIMENTAL NOTES:

Day 0:

Day 1:

Day 2:

Results:
